# Epigenetic Reprogramming of Autophagy Leads to Uncovering a Novel Therapeutic Target for Mutant IDH1 Astrocytomas

**DOI:** 10.1101/2024.03.08.584091

**Authors:** Felipe J Núñez, Kaushik Banerjee, Anzar A. Mujeeb, Ava Mauser, Claire E. Tronrud, Ziwen Zhu, Sadhakshi Raghuram, Maya R. Sheth, Jorge Armando Pena Agudelo, Julio Zelaya, Ayman Taher, Padma Kadiyala, Stephen V. Carney, Maria B. Garcia-Fabiani, Andrea Comba, Mahmoud S. Alghamri, Brandon L. McClellan, Syed M. Faisal, Zeribe C. Nwosu, Hanna S. Hong, Peter Sajjakulnukit, Tingting Qin, Maureen A. Sartor, Mats Ljungman, Joshua D. Welch, Shi-Yuan Cheng, Pedro R. Lowenstein, Joerg Lahann, Costas A. Lyssiotis, Maria G. Castro

## Abstract

Mutant isocitrate dehydrogenase 1 (mIDH1) exhibits a gain of function mutation enabling 2-hydroxyglutarate (2HG) production and epigenetic reprogramming. This leads to enhanced DNA-damage response and radioresistance in mIDH1 gliomas. RNA-seq and ChIP-seq data revealed that human and mouse mIDH1 glioma neurospheres have downregulated gene ontologies (GOs) related to mitochondrial metabolism and upregulated GOs related to autophagy. Decreased mitochondrial metabolism was accompanied by decreased glycolysis, rendering autophagy a source of energy in mIDH1 gliomas. Human and mouse mutant IDH1 glioma cells exhibited increased expression of pULK1-S555 and enhanced LC3 I/II conversion, indicating augmented autophagy. Additionally, scRNA-seq data from human mIDH1 astrocytoma patients’ samples showed decreased mitochondrial metabolism and increased autophagy. We further demonstrate that inhibiting autophagy in vivo by systemic administration of synthetic protein nanoparticles encapsulating siRNA targeting Atg7 sensitized mIDH1 glioma cells to radiation-induced cell death, resulting in tumor regression, long-term survival, and immunological memory. In summary, our work uncovered that autophagy is a critical pathway for survival in mIDH1 gliomas and by blocking this pathway we can elicit radiosensitivity in vitro in human and mouse mIDH1 glioma cells, and in vivo in genetically engineered mouse models. Our data also highlights that blocking autophagy has significant potential for clinical translation.

Graphical Abstract:
Our genetically engineered mIDH1 mouse glioma model harbors IDH1^R132H^ in the context of ATRX and TP53 knockdown. The production of 2-HG elicited an epigenetic reprogramming associated with a disruption in mitochondrial activity and an enhancement of autophagy in mIDH1 glioma cells. Autophagy is a mechanism involved in cell homeostasis related with cell survival under energetic stress and DNA damage protection. Autophagy has been associated with radio resistance. The inhibition of autophagy thus radio sensitizes mIDH1 glioma cells and enhances survival of mIDH1 glioma-bearing mice, representing a novel therapeutic target for this glioma subtype with potential applicability in combined clinical strategies.

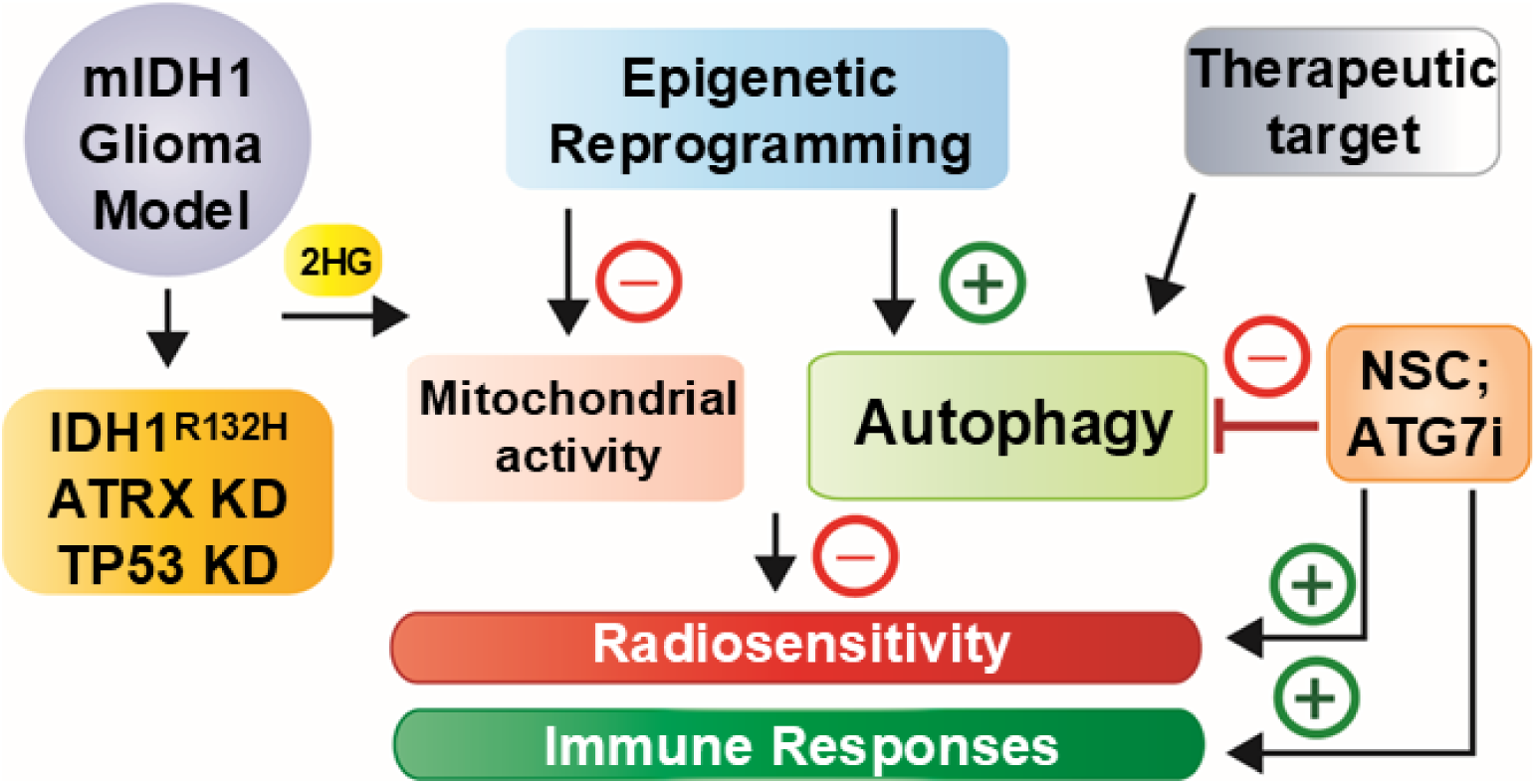

## Introduction

Gliomas are primary brain neoplasms characterized by high morbidity, recurrence, and mortality (1). This pathology includes a genetically and phenotypically heterogeneous group of CNS tumors, which are molecularly classified in different subtypes according to the presence of specific genetic lesions and epigenetic profiles (2, 3). A gain of function mutation in isocitrate dehydrogenase 1 (IDH1^R132H^; mIDH1) is a common genetic alteration in adult gliomas (4). IDH1^R132H^ catalyzes the conversion of α-ketoglutarate (α-KG) to 2-hydroxyglutarate (2HG) (5). 2HG inhibits α-ketoglutarate-dependent enzymes such as histone demethylases (KDMs) and cytosine dioxygenases [ten-eleven translocation (TET) methylcytosine dioxygenases], inducing epigenetic reprogramming of the tumor transcriptome (6). IDH1^R132H^ is associated with a better prognosis in gliomas (4, 7, 8) and impacts multiple cellular functions, including cell differentiation (9), invasion (10), immune response (11–13), and responses to cellular stress and DNA-damage (14–16). We, and others, recently demonstrated that the epigenetic reprogramming in mIDH1 results in enhanced DNA-damage responses and impaired response to radiotherapy (15–18). Furthermore, mIDH1 was shown to elicit cellular metabolic reprogramming that could modify the energetic state of mIDH1 glioma cells and the tumor microenvironment (TME) (16, 19–22). The metabolic alterations in mIDH1 gliomas have been associated with a less aggressive tumor phenotype (16, 23–25). Mutant IDH1 glioma cells lose the ability to use NADPH-reducing equivalents and gain a novel NADPH-coupled α-ketoglutarate-reducing activity (26), resulting in a change in the cellular NADPH to NADP+ ratio (5). This leads to enhanced susceptibility to changes in the cellular redox status and oxidative stimuli in mIDH1 glioma cells (27–30).

Autophagy is a conserved process that allows the recycling of intracellular components including damaged organelles, protein aggregates and cellular waste, maintaining the energetic homeostasis and protecting cells against stress (31). The autophagic pathway involves the generation and degradation of autophagic vesicles, where the cellular material is collected, contained, processed, and recycled (32). This phenomenon occurs in several steps: initiation, expansion, fusion, and degradation, and involves numerous autophagy-related genes (ATG) that participate in an orchestrated assembly, driving the autophagic flux (33, 34). This flux can be activated via cell signaling in response to stimulus such as starvation, cellular stress, or DNA-damage (32, 35, 36). This physiological process plays crucial roles during development and its deregulation is involved in disease progression, including Parkinson’s disease, diabetes, and cancer (37–41). The role of autophagy and the mechanisms underlying its regulation in mIDH1 gliomas have not been fully elucidated. For instance, Garry et al., 2014, showed that mutant IDH1/2HG can induce oxidative stress, autophagy, and apoptosis (14). In addition, Viswanath et al., 2018, showed that 2HG induces a reduction in the endoplasmic reticulum (ER) via a mechanism termed autophagy of the ER (42). They also demonstrated that autophagic flux was higher in both WT-IDH1 cells exposed to 2HG and mIDH1 cells (42).

In this study, we demonstrate that autophagy plays a critical role in mediating DNA damage response and energetic homeostasis in mouse mIDH1 glioma models and human mIDH1 gliomas co-expressing TP53 and ATRX inactivating mutations (2, 15). Through epigenetic, transcriptomic, metabolomic, and signaling pathway analysis, we show that the epigenetic changes in mIDH1 gliomas, with TP53 and ATRX loss of function mutations, modulate the expression of several genes that functionally impact mitochondrial metabolism and upregulate autophagy. We demonstrate that targeting the autophagy pathway by synthetic protein nanoparticles (SPNPs) incapsulating siRNA against ATG7 enhanced radiosensitivity in mIDH1 mouse glioma model, prolonged survival, and resulted in a sustained immunological memory (43, 44). Our findings highlight that autophagy inhibition combined with ionizing radiation (IR) represents an attractive therapeutic strategy that could be implemented for mIDH1 glioma patients.

## Results

### Transcriptomic reprogramming in mIDH1 glioma impacts autophagy and mitochondrial metabolic pathways

For this study, we used the Sleeping Beauty Transposon system to create three genetically engineered mIDH1 mouse glioma models which include the following genetic lesions: (i) NPA/NPAI: *NRAS-G12V*, sh*TP53*, sh*ATRX*, and *IDH1-R132H*; and two RAS independent mouse glioma models: (ii) CPA/CPAI: *CDKN2A* -/-, sh*TP53*, sh*ATRX*, and *IDH1-R132H*; and (iii) RPA/RPAI: *PDGFRα-D842V*, sh*TP53*, sh*ATRX*, and *IDH1-R132H*. We also employed human mIDH1 glioma cell cultures obtained from surgical biopsies and tumor tissue for our “omics” and functional studies. To understand the biological significance of the transcriptomic alteration in mIDH1 glioma, we first analyzed the RNA-seq data generated from our previously described mIDH1 mouse glioma model (NPA/NPAI; Figure 1A) (15). We specifically focused on the differential expression of genes related to cellular metabolism and energetic homeostasis. We observed that mIDH1 neurospheres (NS) had upregulated genes related to autophagy gene ontologies (Figure 1B; Supplemental Table 1). In addition, these cells have a profound downregulation of electron transport chain (ETC) complex genes (Figure 1C; Supplemental Table 2), suggesting impaired mitochondrial function and bioenergetics. This revealed a downregulation of mitochondrial metabolism pathways and an enrichment in autophagy (Figure 1D). Our data also revealed downregulation of the mitochondria respiratory chain complex, the electron transport chain, and mitochondrial oxidative phosphorylation pathways, indicating reduced mitochondrial metabolic activity in mIDH1 cells (Figure 1E-F). To confirm the differences in the gene ontologies (GOs) related to autophagy observed in our RNA-seq data, we also analyzed scRNA-seq data obtained from mIDH1 glioma patients. We found alterations in GOs related to several autophagy genes including *ULK1*, *UVRAG*, *ATG4b*, and *ATG7* (Supplemental Figure 1). Additionally, we have analyzed data from The Cancer Genome Atlas Program (TCGA) and found significant differences in mRNA expression in *UVRAG*, *ATG9b*, *ATG7*, and *ULK1* in mIDH patients when compared to WT-IDH patients (Supplemental Figure 2). To determine the effect of mIDH1 on cell metabolism, we performed mass spectrometry-based metabolomics analysis (Supplemental Figure 3). We observed that the disrupted mitochondrial respiration gene program (Figures 1G-H; and Supplemental Figures 4-5) is accompanied by the accumulation of TCA cycle metabolites (Figure 1G), suggesting suppressed pathway activity. Consistently, TCA cycle anaplerotic substrates such as aspartate and glutamate (Figure 1H and Supplemental Figure 6), also accumulated in the mIDH1 cells, supporting that mIDH1 disrupts metabolic activities in the glioma NS (Figure 1I). To validate that our metabolic findings were due to mIDH1 gliomas’ biology and not to the slower proliferative state of mIDH1 cells, we performed colorimetric assays to determine both Malate and Succinate levels in two different genetically engineered mouse glioma models under reduced growth conditions (Supplemental Figure 7). We found that there were significantly higher levels of Malate and Succinate in mIDH1 mouse glioma models when compared to their WT-IDH1 mouse glioma model counterparts, indicating that the alterations seen in mIDH1 glioma metabolism are due to the mIDH1 biology rather than differing growth rates (Supplemental Figure 7).

**Figure 1:**
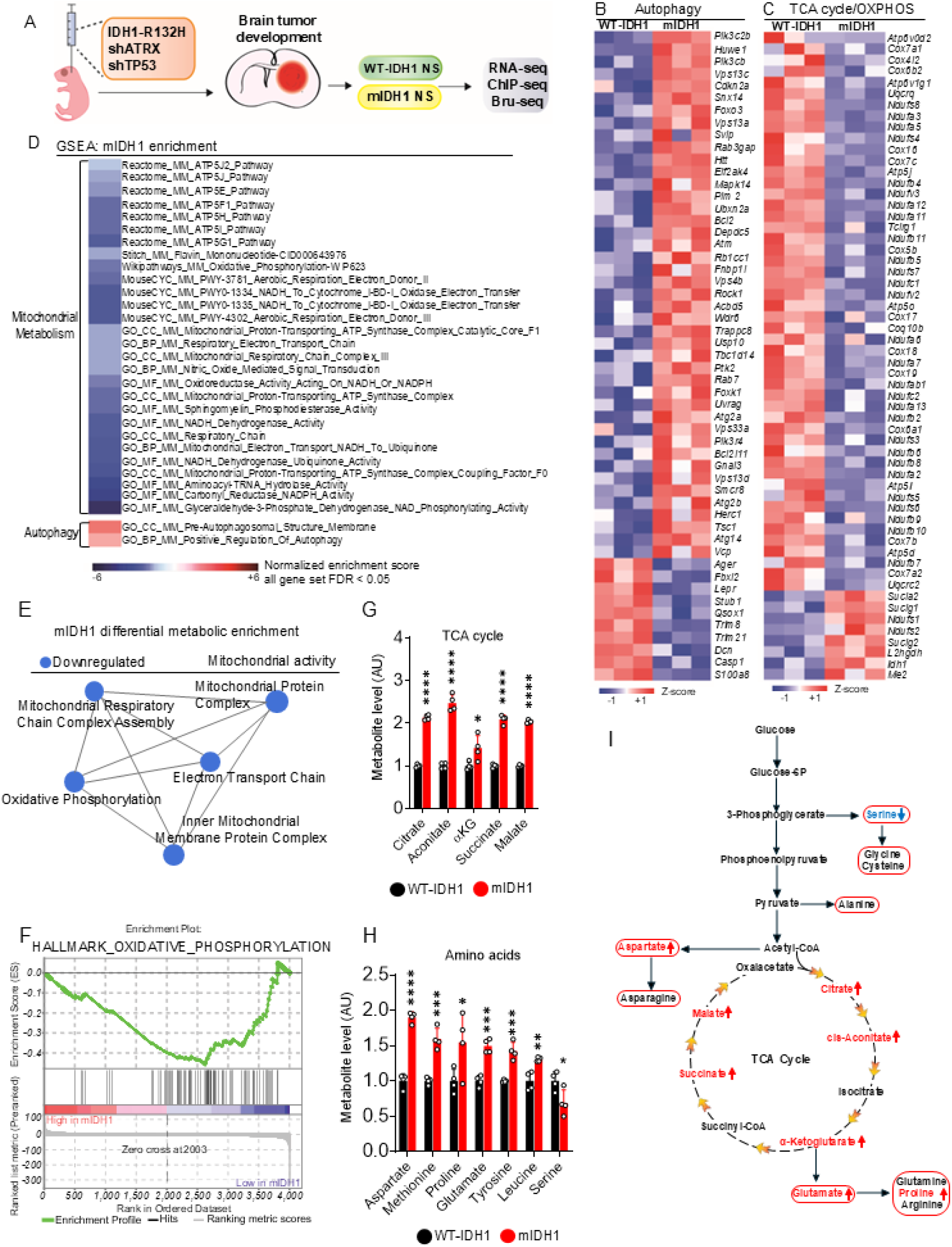
mIDH1 glioma has upregulated autophagy and disrupts mitochondrial metabolism. (**A**) Illustration of genetically engineered mIDH1 glioma models harboring ATRX and TP53 knockdown and molecular analysis performed. (**B-C**) Heatmaps showing RNA-seq gene expression pattern for (B) autophagy and (C) electron transport chain (ETC) genes in WT-IDH1 and mIDH1 NS (n = 3 biological replicates; up-regulated genes are red; down-regulated genes are blue; FDR ≤ 0.05; ≥ 1.5-fold). (**D**) GSEA comparing transcriptional changes in mIDH1 NS versus WT-IDH1. Negative/positive normalized enrichment scores (blue/red scale; FDR < 0.05) show GO terms linked to metabolism (mitochondrial, OXPHOS) (downregulated) and autophagy (upregulated) in mIDH1 NS. (**E**) Pathway enrichment map of differentially enriched (DE) genes in mIDH1 NS versus WT-IDH1. Blue nodes illustrate down-regulation in mIDH1 NS (\**P* < 0.05; overlap cutoff>0.5) related to mitochondrial metabolic activity. (**F**) Enrichment plots of gene sets significantly downregulated in mIDH1 NS with decreasing expression levels of OXPHOS genes detected by GSEA. Green curves show enrichment score and reflect degree of each gene (black lines) and is represented at bottom of ranked gene list. (**G-H**) Metabolomics profiling data showing relative levels of (G) TCA cycle intermediates and (H) amino acid levels in WT and mIDH1 NS (a.u.: arbitrary unit). Results expressed as mean±SD.\**P* < 0.05; \*\*\**P* < 0.001; \*\*\*\**P* < 0.0001; two-way ANOVA. (**I**) Illustration showing TCA cycle intermediates and amino acids disrupted in mIDH1 NS versus WT-IDH1 NS. Significantly altered amino acids depicted in red.

### Histone hypermethylation in mIDH1 glioma NS is associated with metabolism and autophagy pathways

To evaluate the impact of epigenetic reprogramming mediated by mIDH1 on tumor metabolism and autophagy, we used previously published ChIP-seq data generated through native ChIP followed by sequencing (Figure 2A) (15). To determine the impact of the mutation in IDH1 on histone marks’ hypermethylation levels, we measured H3K4me3, H3K27me3, and H3K36me3 via western blot in four different glioma cell model systems, including human mIDH1 glioma cells obtained from surgical biopsies (SF10602) (45). Mutant IDH1 glioma models tested were as follows: (i) NPA/NPAI mouse NS, (ii) RPA/RPAI mouse NS, (iii) WT-IDH1 SJGBM2/mIDH1 SJGBM2 human glioma cells, and (iv) SF10602 human glioma cells treated with and without mIDH1 inhibitor AG-881 (Supplemental Figure 8). We found that there was a significant increase in expression levels for all histone marks in mIDH1 cells when compared to WT-IDH1 glioma cells (Supplemental Figure 8). We previously demonstrated the efficacy of mIDH1 inhibitors at decreased production of 2HG in both mouse and human mIDH1 glioma cells (15, 18). We then proceeded to evaluate the differential enrichment at promoter regions of H3K4me3, a transcriptional activation mark, and of H3K27me3, a transcriptional repressive mark in mIDH1 compared to WT-IDH1 NS (Figure 2B-D). Differential H3K4me3 enriched GO terms included regulation of several cell metabolic processes (Figure 2B), resulting in an epigenetic regulation of these metabolic pathways in mIDH1 cells. GO terms enriched in H3K27me3 histone indicate epigenetic repression in mIDH1 versus WT-IDH1 glioma NS (Figure 2C). To better understand the relationship between epigenetic regulation and autophagy in mIDH1 glioma, we conducted comprehensive CUT&Run analyses that demonstrated autophagy enrichment, including differential deposition of H3K4me3 and H3K27ac marks at promoter and regulatory regions of genes which mediate autophagy (Figure 2D). Of note, we observed differential peaks for H3K4me3 around the promoter and genomic regions of the autophagy related gene *ATG9b* (46, 47), as well as differential peaks for H3K27ac around enhancer regions in mIDH1 NS when compared to WT-IDH1 NS (Figure 2E). Consistent with our previous results, our Bru-seq analysis performed on mIDH1 NS showed a higher transcription rate for *ATG9b* (> 2-fold) versus WT-IDH1 NS (Figure 2F). Further analysis revealed that mIDH1 cells express significantly higher levels of ATG9b protein in both mouse and human mIDH1 glioma cells (Figure 2G-H, Supplemental Figure 9). These results indicate that mIDH1, through epigenetic reprograming alters metabolic and autophagic pathways in glioma cells (Figure 2I).

**Figure 2:**
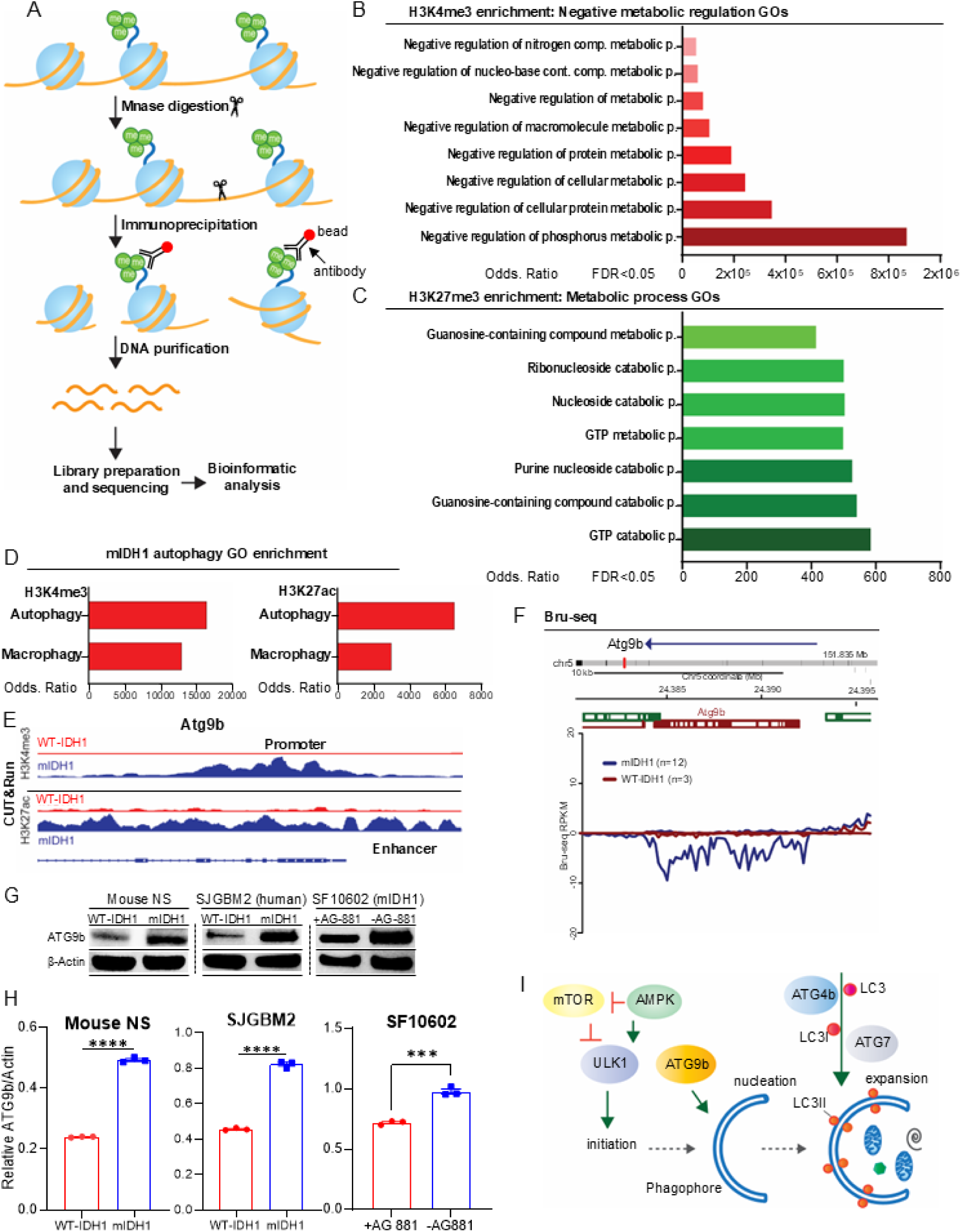
Epigenetic reprogramming in mIDH1 glioma decreases metabolism and enhances autophagy in mIDH1 glioma cells. (**A**) Native H3K4me3 ChIP-seq diagram performed on mIDH1 NS and WT-IDH1 NS. (**B-C**) Genes enriched in H3K4me3 or H3K27me3 linked to distinct functional GO terms by ChIP-seq analysis. Bar graphs representing GO terms containing genes having enrichment of H3K4me3 (red) or H3K27me3 (green) at their promoter regions in mIDH1 NS. GO terms’ significance determined by FDR (<0.05); enrichment expressed as OR. (**D**) Genes enriched in H3K4me3 or H3K27ac linked to distinct functional GO terms by CUT&Run. (**E**) H3K4me3 and H3K27ac occupancy in specific genomic regions of *ATG9b*. Y-axis represents estimated number of immunoprecipitated fragments normalized to total number of reads in each dataset. Reference sequence (RefSeq) gene annotations are shown. (**F**) Bru-Seq traces showing differential transcriptional rates (< 1.5-fold; *P* < 0.05) of *ATG9b* in mIDH1 NS (blue) versus WT-IDH1 NS (red). Arrow indicates sequence strand reading direction; positive y-axis represents positive strand signal of transcription moving left to right; negative y-axis represents negative strand signal of transcription moving right to left. Vertical red mark indicates *ATG9b* position within chromosome. Genes and chromosome locations indicated on maps. Data expressed in reads per kilobase per million mapped reads (RPKM). Gene maps generated from RefSeq. (**G**) WB showing ATG9b protein expression in WT-IDH1 NS, mIDH1 NS human glioma SJGBM2 WT-IDH1 and mIDH1, and mIDH1 human glioma cells, SF10602, ±IDH1^R132H^ inhibitor AGI-5198 (5µM). (**H**) ImageJ densitometric quantification of the western blots for ATG9b and β-actin shown in (G). Errors bars represent SEM from independent biological replicates (n = 3). \*\*\*\**p*<0.0001; unpaired t test. (**I**) Diagram of initial steps of autophagy pathways, including phagophore formation with participation of ATG9b protein in the nucleation process.

### Epigenetic reprogramming in human mIDH1 glioma cells impacts autophagy related genes

In addition to *ATG9b*, the formation of phagophores and autophagosomes requires upregulation of several other genes including *UVRAG*, *STK26* (MST4), *ATG7* and *MAP1LC3B* (Figure 3A). To further validate the role of mIDH1 in the epigenetic regulation of these genes, we performed ChIP-seq analysis on human mIDH1 SF10602 cells, in the presence or absence of the mIDH1 inhibitor (Figure 3B). SF10602 cells have a mixed population of both mIDH1 and WT-IDH1 alleles (Supplemental Figure 10). Our results showed that the specific inhibition of mIDH1 alters peak enrichment for H3K36me3, H3K4me3, or H3K27ac marks (Figure 3C-I) around promoter and enhancer regions of *UVRAG* (Figure 3C), *ATG7* (Figure 3D), *MAP1LC3B* (Figures 3E-F), *STK26* (MST4) (Figure 3G), and *ATG9b* (Figure 3H-I). To ensure that these enrichment marks are due to mIDH1 glioma biology, we examined the gene regions within nearby autophagy related genes, that are not differentially regulated, in mIDH1 SF10602 cells versus SF10602 cells treated with mIDH1 inhibitor (Supplemental Figure 11). We found that *VGLL4* (next to *ATG7*), *ZCCHC14* (next to *UVRAG*), and *WNT11* (next to *MAP1LC3B*) are not differentially regulated in mIDH1 SF10602 cells versus SF10602 cells treated with mIDH1 inhibitor (Supplemental Figure 11).

**Figure 3:**
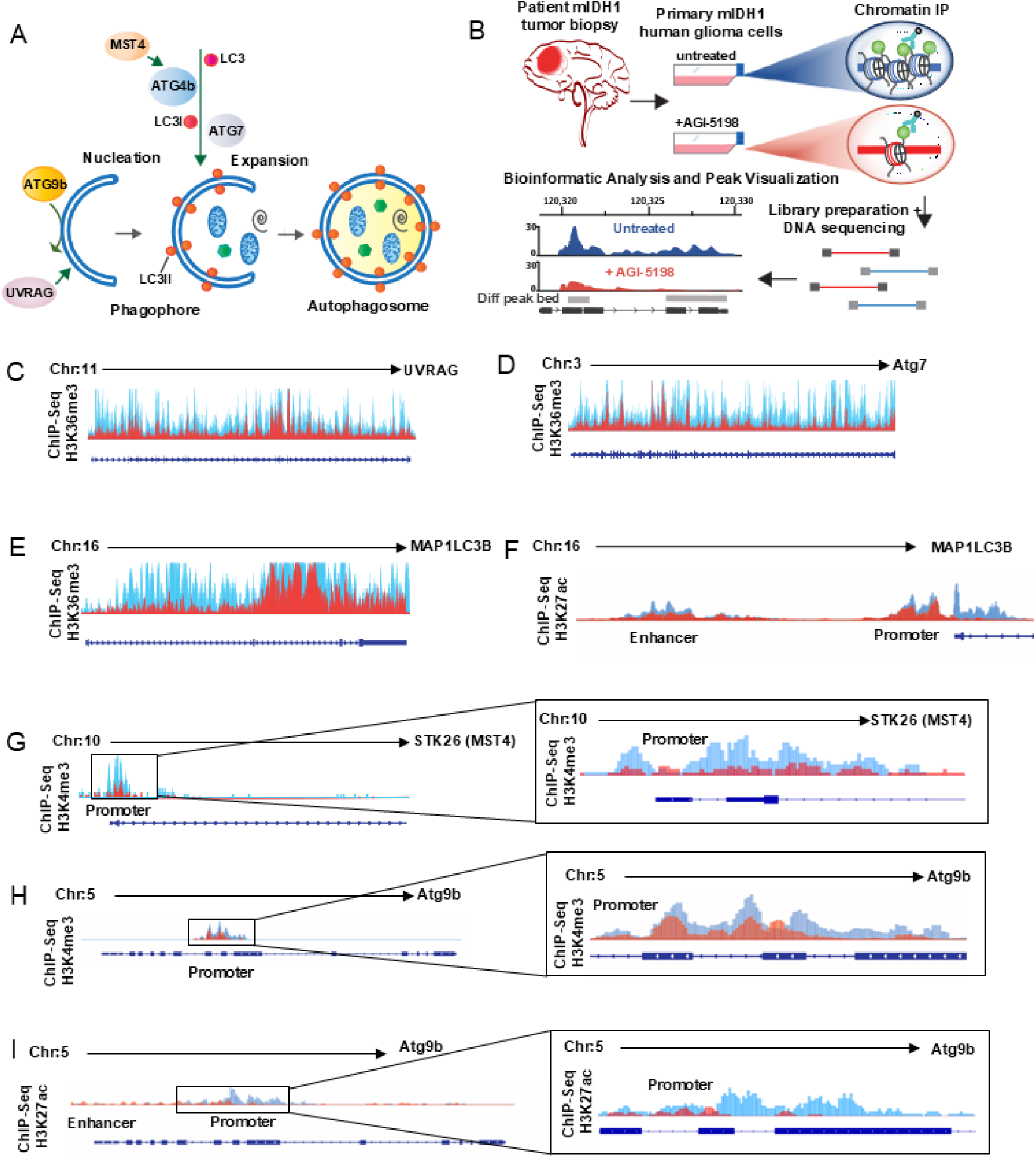
Epigenetic regulation of autophagy related genes in mIDH1 human glioma cells. (**A**) Diagram of autophagosome formation in the autophagic pathway involving autophagy related genes: UVRAG, ATG7, LC3 (*MAP1LC3B*), and MST4 (*STK26*). (**B**) Diagram of experimental design of ChIP-seq analysis performed in primary mIDH1 human glioma cells, SF10602, ±IDH1^R132H^ inhibitor AGI-5198 (5 µM), prior to ChIP. (**C-I**) (C-F) H3K36me3 and (G-I) H3K4me3, H3K27ac occupancy in specific genomic regions of autophagy related genes *UVRAG*, *ATG7*, *MAP1LC3B* (LC3), *STK26* (MST4), and *ATG9b*. The y-axis of each profile represents the estimated number of immunoprecipitated fragments at each position normalized to the total number of reads in each dataset. RefSeq gene annotations are shown. Differential peaks (FDR < 0.05) in untreated SF10602 mIDH1 glioma cells are represented in blue compared to AGI-5198 treated cells in red (n = 3 biological replicates per group).

### Single Cell RNA-seq shows an enhancement of autophagy in human mIDH1 glioma cells

To assess the impact of epigenetic reprogramming mediated by mIDH1 on autophagy, scRNA-seq analysis was conducted on SF10602 cells, under both the presence and absence of the mIDH1 inhibitor AGI-5198 (Figure 4A). The examination included an evaluation of enriched gene ontology (GO) terms associated with autophagy. The results demonstrated that the inhibition of mIDH1 led to decreased positive regulation of autophagy (Figure 4B-C), increased negative regulation of autophagy (Figure 4D; Supplemental Figure 12A), decreased cellular components of autophagosome (Figure 4E; Supplemental Figure 12B), and decreased processes utilizing autophagic mechanisms (Figure 4F-G). The observed changes in gene ontology terms associated with autophagy highlight a significant influence of mIDH1 on the regulation of autophagy-related processes. The specific alterations identified, such as the modulation of positive and negative regulation of autophagy, as well as changes in cellular components and processes related to autophagosome function, provide valuable insights into the intricate relationship between IDH1 mutation and the dysregulation of autophagy pathways in glioma. Taken together, these findings suggest a link between IDH1 mutation and the upregulation of autophagy-related pathways in glioma.

**Figure 4:**
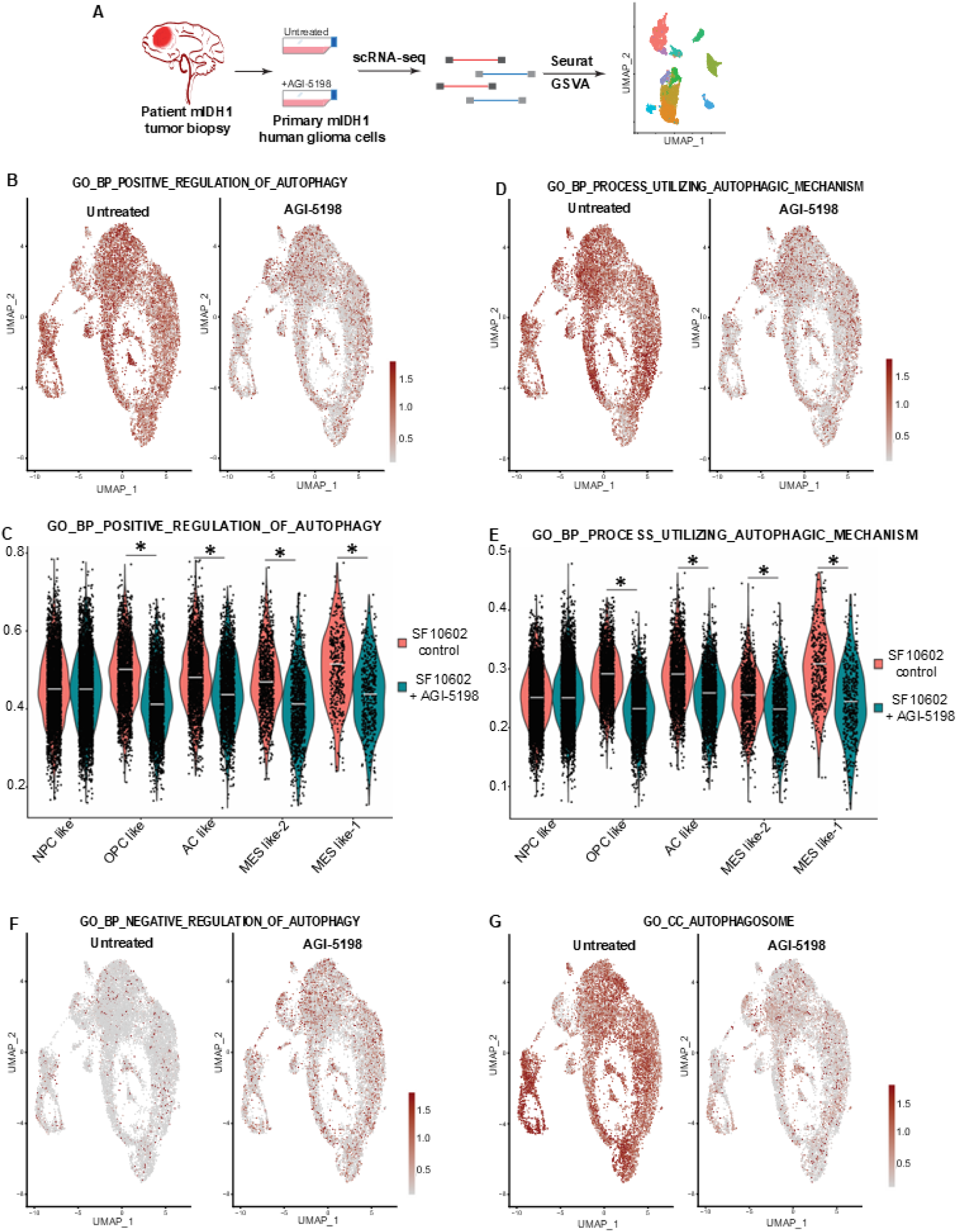
Single Cell RNA-seq shows an enhancement of autophagy in human mIDH1 glioma cells. (**A**) Diagram of experimental design of scRNA-seq analysis performed in primary mIDH1 human glioma cells, SF10602, ±IDH1^R132H^ inhibitor AGI-5198, prior to sequencing. (**B**) GO term enrichment score for GO biological process of positive regulation of autophagy. (**C**) Violin plot of scRNA-seq in human mIDH1 glioma cells related to GO enrichment score for GO biological process of positive regulation of autophagy. (**D**) GO term enrichment score for GO biological process of utilizing autophagy mechanism (**E**) Violin plot of scRNA-seq in human mIDH1 glioma cells related to GO term enrichment score for GO biological process of utilizing autophagy mechanism. (**F-G**) GO term enrichment score for GO cellular component of (F) negative regulation of autophagy and (G) autophagosome. Significance of the violin plots was measured via the Wilcoxon Rank Sum Test from the Seurat/Presto package. \**p* < 0.05.

### Mutant IDH1 NS exhibit altered mitochondrial activity and slower proliferative rate

To gain further insight into the metabolic activity of mIDH1 glioma cells, we used the Seahorse extracellular flux analyzer to compare the bioenergetic profiles of primary NS isolated from WT-IDH1 and mIDH1 tumors. Given our transcriptomics analysis indicating dysregulated mitochondrial activity, and the accumulation of the TCA cycle metabolites, we performed the Mito Stress Test to assess differences in mitochondrial function and metabolic profile between WT-IDH1 and three different mIDH1 NS clones. When compared to WT-IDH1, mIDH1 NS exhibited lower oxygen consumption rate (OCR) at baseline, suggesting mIDH1 glioma cells utilize mitochondrial respiration to a lesser extent compared to that of WT-IDH1 cells (Figure 5A). In line with this finding, the maximal rate of respiration in mIDH1 NS was lower compared to that of WT-IDH1 cells, as seen by the reduction in OCR following treatment with the mitochondrial uncoupler 4-(trifluoromethoxy) phenylhydrazone (FCCP) (Figure 5A and Supplemental Figure 13A). Additionally, examination of glycolytic activity, measured by the extracellular acidification rate (ECAR), revealed that mIDH1 NS exhibit a significant reduction (∼50%) in ECAR compared to WT-IDH1 counterparts at baseline (Figure 5B and Supplemental Figure 13B). Together, these findings demonstrate that mIDH1 NS exhibit less mitochondrial respiration and glycolytic activity indicative of slower proliferative rate when compared to the WT-IDH1 NS (Figure 5C).

**Figure 5:**
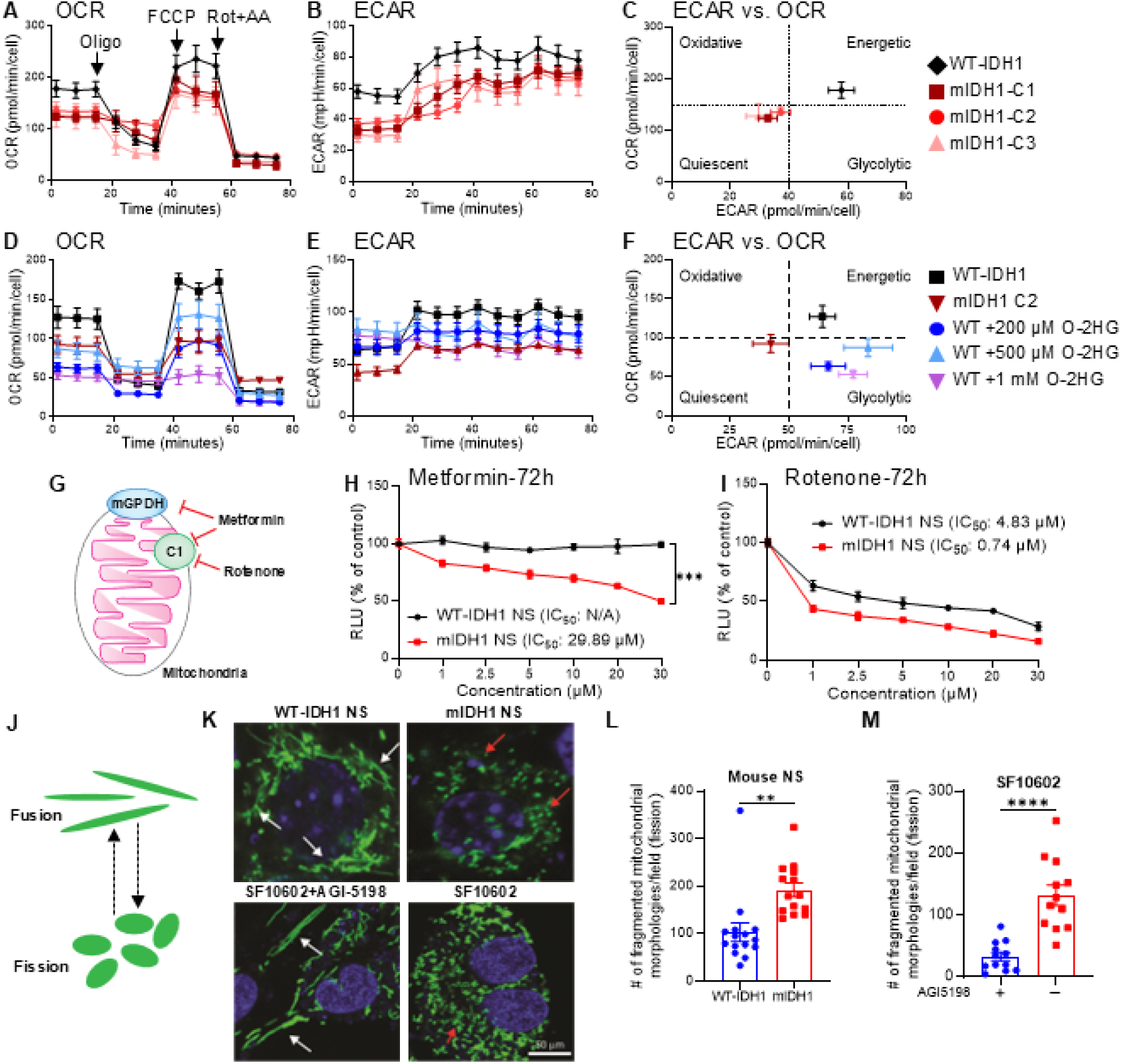
mIDH1 glioma cells exhibit a slower proliferative state. (**A**) Mitochondrial respiration of glioma NS derived from WT-IDH1 or mIDH1 tumors (Clones 1-3) examined through Mito Stress Test. Error bars depict SD of technical replicates from representative plot of 3 independent experiments. Mitochondrial respiration determined by oxygen consumption rates (OCR). (**B**) Glycolytic activity determined by extracellular acidification rates (ECAR) in mIDH1 NS and WT-IDH1 NS. (**C**) Metabolic profile of WT-IDH1 NS and mIDH1 NS comparing ECAR versus OCR. (**D**) Mito Stress Test performed on WT-IDH1 NS, mIDH1 NS, and WT-IDH1 NS cultured with cell-permeable (R)-2HG (WT-IDH1 NS + O-2HG) for 4h at indicated concentrations. Error bars depict SD of technical replicates from representative plot of 3 independent experiments. (**E**) ECAR of WT-IDH1 NS, mIDH1 NS, and WT-IDH1 NS + O-2HG. (**F**) Metabolic profile of WT-IDH1 NS, mIDH1 NS, and WT-IDH1 NS + O-2HG, comparing ECAR versus OCR. Significance determined by one-way ANOVA. \**P* < 0.05, \*\**P* < 0.01, \*\*\**P* < 0.001, \*\*\*\**P* < 0.0001. (**G**) Diagram representing mitochondrial complex 1 (C1) inhibition by metformin and rotenone. (**H-I**) Cell viability assay of WT-IDH1 NS and mIDH1 NS treated with (H) metformin and (I) rotenone for 72h at increasing concentrations. Cell viability expressed in relative LUC units in percent of control (100%). Linear regression model and generalized additive models (GAM) for flexible regression modeling were used to model dose-response relationship. \*\*\**P* < 0.001 (**J**) Diagram representing mitochondrial morphology: fusion (elongation) and fission (division). (**K**) Mitochondrial morphology observed with confocal microscopy in WT-IDH1 NS, mIDH1 NS, and SF10602±AGI-5198, labeled with MitoTracker Green FM. White arrows indicate mitochondrial fusion; red arrows indicate mitochondrial fission. The scale bar represents 50µm. (**L-M**) (L) Quantification of fragmented mitochondrial morphologies in mouse WT-IDH1 and mIDH1 NS using Image J2 software. (M) Quantification of fragmented mitochondrial morphologies in endogenous mIDH1 expressing human patient-derived secondary recurrent GBM cells (SF10602 and SF10602 with AGI5198) using Image J2 software. Standard settings were applied for the Mitochondrial Morphology Macro, the MitoLoc plugin and the MiNa plugin. For the Particle Analyzer method, auto thresholding was applied on images before applying the Analyze particles function. Confocal images were deconvoluted using the Iterative Deconvolve 3D plugin, without a wiener filter gamma and with a low pass filter of 1, a maximum number of iterations of 10 and a termination of iterations at 0.010. Data were collected from three independent experiments.

Mutant IDH1 glioma cells promote the conversion of α-ketoglutarate to (R)-2-hydroxyglutarate (2HG), an inhibitor of α-ketoglutarate-dependent epigenetic enzymes. This can modify the transcriptome of tumor cells, and therefore indirectly influence metabolism, in a process that typically requires multiple days and cell divisions (48). In contrast, 2HG can also directly impact metabolism by inhibiting metabolic enzymes that use α-ketoglutarate as a substrate, including but not limited to transaminases. A recent study demonstrated that 2HG can inhibit BCATs, a mechanistic understanding of which identified novel metabolic dependencies in IDH mutant brain tumors (49). To determine whether 2HG influences the metabolic profile in our models of mIDH1 tumor cells, WT-IDH1 NS were cultured with cell-permeable (R)-2HG (O-2HG) for 4h, after which we performed Seahorse extracellular flux analysis. WT-IDH1 NS supplemented with O-2HG appeared more similar in basal and maximal respiration to mIDH1 (Figure 5D), whereas glycolytic activity was impacted to a lesser degree (Figure 5E). This data illustrates that 2HG supplementation can directly alter the energetic state of glioma NS (Figure 5F), which likely contributes in part to the observed metabolic alterations in mIDH1 cells.

Based on the metabolic defects observed with exogenous 2HG, we then evaluated if mIDH1 glioma cells are more susceptible to mitochondrial inhibitors than those expressing WT-IDH1. To evaluate the sensitivity of mIDH1 to mitochondrial inhibition, we treated mouse WT-IDH1 and mIDH1 cells with different concentrations of the mitochondrial complex 1 (C1) inhibitors, metformin and rotenone (Figure 5G), after which we evaluated cell viability (Figure 5H-I and Supplemental Figure 14). Results showed that mIDH1 cells were more sensitive to rotenone and metformin treatment than WT-IDH1. Also, inhibition of mitochondrial complex I, renders the mIDH1-mouse cells more susceptible to radiation (Supplemental Figure 14). We found a significant difference in cell viability when the mIDH1 cells are treated with either rotenone or metformin in combination with IR compared to radiation alone (Supplemental Figure 14). In addition, we evaluated the sensitivity of two human mIDH1 cell cultures to treatment with mitochondrial complex 1 inhibitors, metformin and rotenone, measuring cell viability (Supplemental Figures 15-16). Our results showed that mIDH1 SJGBM2 cells and SF10602 cells are more sensitive to metformin alone and metformin in combination with IR (Supplemental Figure 15), and to rotenone alone and rotenone in combination with IR (Supplemental Figure 16). To confirm that the sensitivity to mitochondrial inhibition was due to mIDH1 biology and not to slower proliferation rates, we treated both WT-IDH1 and mIDH1 human glioma cells with metformin ± IR in low growth factor containing media (Supplemental Figure 17). Our results indicate that mIDH1 SJGBM2 cells and SF10602 cells are more sensitive to metformin both alone and when combined with IR, when grown in growth factor deprived conditions. We have also performed this experiment treating cells with rotenone, and these results also indicate that mIDH1 SJGBM2 cells and SF10602 cells are more sensitive to rotenone both alone and when combined with IR, when grown in growth factor deprived conditions (Supplemental Figure 18).

### IDH1 mutation correlates with increases in fission state mitochondrial profile

Mitochondria undergo fusion (elongation) and fission (fragmentation) (Figure 5J), which are critical processes for mitochondrial homeostasis and energy adaptation in response to metabolic changes (50). Fusion homogenizes the contents of mitochondria resulting in mitochondrial elongation. Fission consists of mitochondrial fragmentation promoting clearance of damaged mitochondria, closely related with autophagy/mitophagy (50, 51). By performing live cell labeling, we analyzed the mitochondrial morphology in human and mouse glioma cells expressing WT-IDH1, mIDH1 treated with vehicle, or the mIDH1 inhibitor. WT-IDH1 NS and mIDH1 human glioma cells treated with mIDH1 inhibitor AGI-5198 (SF10602+AGI5198) showed a typical mitochondrial fusion morphology, whereas mIDH1 mouse (mIDH1 NS) and untreated human glioma cells (SF10602) presented a mitochondrial morphology corresponding to fission state (Figure 5K). To further validate these findings, we quantified the number of mitochondria in the fission state in all experimental groups. We found that mIDH1 mouse glioma NS had significantly higher quantities of mitochondria in the fission state (mean ± SEM = 192 ± 20 mitochondria in fission state) when compared to WT-IDH1 mouse glioma NS (mean ± SEM = 103 ± 15 mitochondria in fission state; *P* < 0.005; Figure 5L). Furthermore, untreated mIDH1 SF10602 cells had significantly higher quantities of mitochondria in the fission state (mean ± SEM = 132 ± 17 mitochondria in fission state) when compared to SF10602 cells treated with mIDH1 inhibitor (mean ± SEM = 33 ± 7 mitochondria in fission state; p < 0.0001; Figure 5M).

In addition, we analyzed mitochondrial morphology in both WT-IDH1 and mIDH1 genetically engineered mouse cells under reduced growth conditions (Supplemental Figure 19A). We found that even under reduced growth conditions, the number of mitochondrial displaying fission morphology was significantly increased in mIDH1 mouse glioma cells (mean = 270 mitochondria in fission state) when compared to WT-IDH1 mouse glioma cells (mean = 69 mitochondria in fission state; p < 0.001; Supplemental Figure 19B). These results are consistent with the decreased mitochondrial activity observed in mIDH1 glioma cells, suggesting that mIDH1 glioma cells undergo mitochondrial changes to achieve energetic homeostasis, which can involve the autophagy/mitophagy process.

### IDH1^R132H^ is associated with increased autophagy activation in glioma cells

Our results described above indicate that mIDH1 glioma cells have a disrupted mitochondrial activity with a consequently lower energetic charge when compared with WT-IDH1 cells (Figure 5). This is in line with previous studies indicating that metabolite changes associated with IDH1 mutation alter the energic state of glioma cells (23, 25, 52). However, mIDH1 glioma cells can grow and develop tumors in the brains of animals, indicating that they could use alternative mechanisms to generate energy and maintain cell survival. According to our RNA-seq, CHIP-seq, and CUT&RUN data, autophagy is upregulated in mIDH1 glioma cells in correlation with a functional disruption of mitochondrial activity. We then evaluated autophagy activity by protein expression and phosphorylation state of key regulators in the autophagic pathway, including pULK1 (S555) and pULK1 (S757), which are associated with autophagy activation and inhibition, respectively. We also evaluated expression of UVRAG, MST4, and pATG4 that participate in the nucleation and expansion process, involved in autophagosome formation in both mIDH1 and WT-IDH1 NS (Figure 6A). Western blot (WB) analysis revealed that in both mouse and human glioma cells, IDH1 mutations results in increased expression of pULK1-(S555) but decreased expression of pULK1-(S757), when compared with WT-IDH1 glioma cells (Figures 6B-C; see quantification in Supplemental Figures 20-22). Moreover, mIDH1 glioma cells showed upregulation of UVRAG, MST4, and pATG4b (53) (Figures 6B-C; Supplemental Figures 20-22). Interestingly, inhibition of mIDH1 using AG-881 in mIDH1 SF10602 glioma cells reverted the expression pattern of all the key autophagy regulators evaluated (Figure 6D; Supplemental Figure 22). We also measured the protein levels of several key autophagy regulators in an NRAS independent genetically engineered mIDH1 mouse glioma model (RPA/RPAI) and found that mIDH1 cells had higher levels of proteins associated with autophagy activation when compared to WT-IDH1 cells (Supplemental Figure 23). Furthermore, when we treated two different mIDH1 mouse glioma cell models with α-ketoglutarate (α-KG), we found that the mIDH1 cells reverted the expression patterns showing significant reduction in several key autophagy regulators including pATG4b, UVRAG, ATG7, and ATG9b (Supplemental Figure 24).

**Figure 6:**
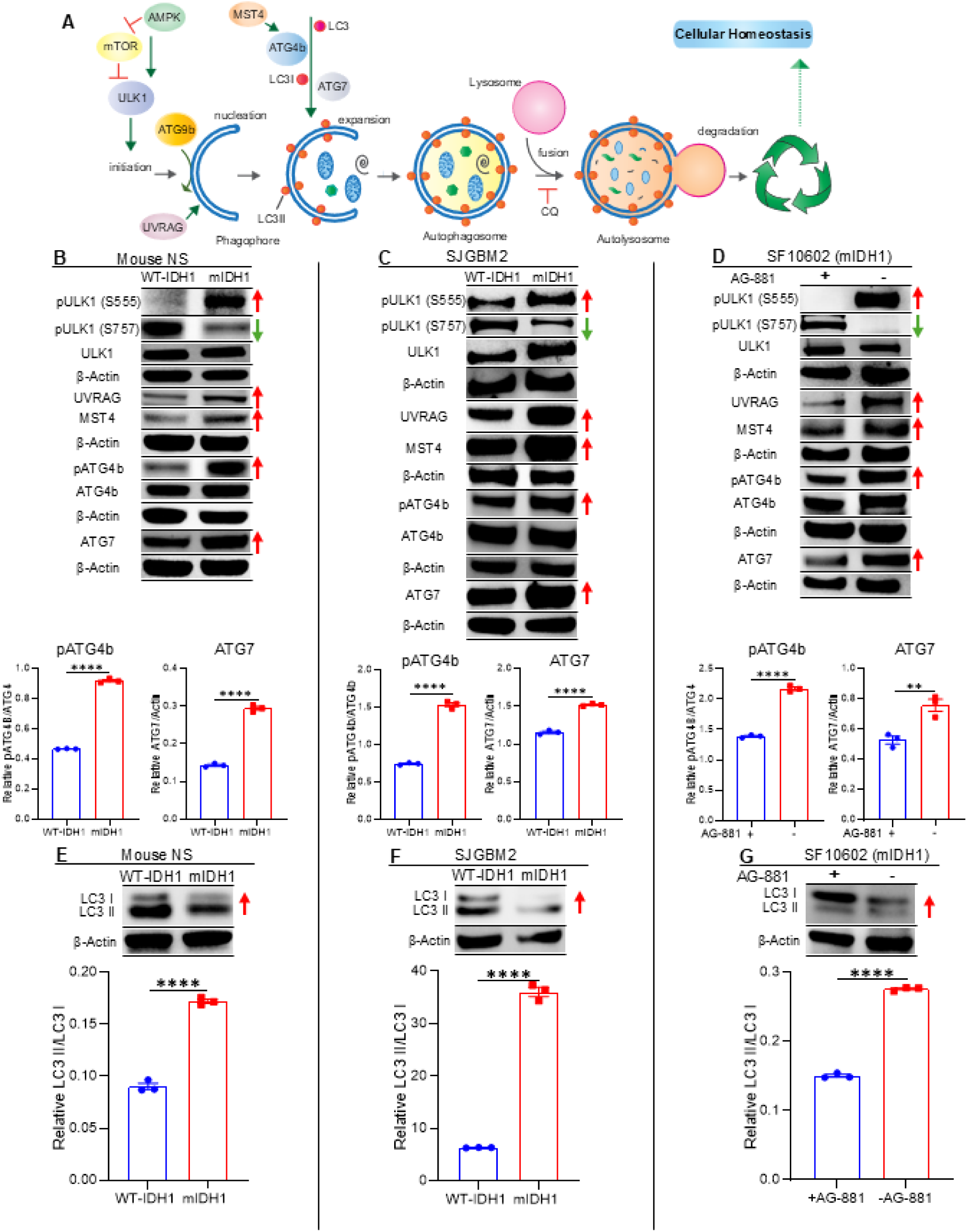
Enhanced autophagy activity in mIDH1 glioma cells. (**A**) The autophagy pathway is responsible for the recycling of cellular residues and damaged organelles thus maintaining hemostasis. This process includes initiation, expansion and degradation steps, where several proteins participate in the formation and transformation of the different cellular structures involved (phagophores, autophagosomes, lysosomes). (**B-D**) WB assay showing the expression of proteins involved in the autophagy pathways and the activated form of them, evaluated in (B) mIDH1 and WT-IDH1 mouse NS, (C) human glioma SJGBM2, and in (D) mIDH1 SF10602 mIDH1 human gliomas cells ± IDH1^R132H^ inhibitor AG-881. The red arrows indicate the proteins and phosphorylated (p) status related with autophagy activation (pULK1 (S555), UVRAG, MST4, pATG4B (S383), ATG7) whereas the green arrows indicate the phosphorylated (p) status related with autophagy inhibition (pULK1 (S757); β-actin, loading control. Bar graph represents densitometric analysis of the ratio of pATG4b/β-actin and ATG7/β-actin WB assay. \*\**P* < 0.005; \*\*\*\**P* < 0.0001; T test. Bars represent means ± SEM (n = 3 technical replicates). (**E-G**) WB assay showing the conversion of LC3I to LC3II as indicator of autophagy activation (A). The increased ratio of LC3II/LC3I (red arrows) indicate autophagy activation. β-actin is a loading control. Bar graph represents densitometric analysis of the ratio of LC3II/LC3I WB assay. \*\*\*\**P* < 0.0001; T test. Bars represent means ± SEM (n = 3 technical replicates).

Another important marker of autophagy activation is the conversion of LC3-I to LC3-II (Figure 6A). A cytosolic form of LC3 (LC3I) is cleaved by ATG4b then conjugated to phosphatidylethanolamine to form LC3-phosphatidylethanolamine conjugate (LC3II), which is recruited to autophagosomal membranes (54) during the expansion step of autophagy (Figure 6A). Thus, the presence and increased intracellular ratio of LC3II over LC3I is a signal of autophagy activation. Results showed that in mIDH1 glioma cells, there was more than 4-fold increase of LC3II/LC3I in both human and mouse glioma cells (Figures 6E-F). As expected, blocking mIDH1 results in 2-fold decrease in the LC3II/LC3I ratio (Figure 6G). We also measured the protein levels of LC3I/II in an NRAS independent genetically engineered mouse model and found that there was a 3-fold increase of LC3II/LC3I ratio in mIDH1 cells (RPAI) versus WT-IDH1 cells (RPA; *P* < 0.0001; Supplemental Figure 25). These results indicate that IDH1 mutation is associated with autophagy activation in gliomas.

### Autophagy flux is enhanced in mIDH1 glioma cells

Autophagy is a continuous process which ends in a recycling of cellular materials (Figure 6A). During autophagic flux, autophagosomes fuse with lysosomes forming autolysosomes which degrade autophagosome contents (55). Using a reporter system where LC3 is fused to both GFP and mCherry (Supplemental Figure 26) (56), we found that mIDH1 glioma cells had increased number of both autophagosomes and autolysosomes when compared with WT-IDH1 cells (autophagosomes > 10-fold increase, *P* < 0.001; autolysosome > 1.4-fold increase in mIDH1 cells, *P* < 0.001). The increase in autophagosome and autolysosomes could reflect increased autophagy or stalled autophagy. To discern between these two fates, we blocked autophagy with chloroquine (CQ), and we observed an increase in the accumulation of LC3-GFP in mIDH1 glioma cells (autophagosomes > 1.6-fold increase in CQ treated mIDH1 cells, *P* < 0.01; no significance observed in autolysosomes in CQ treated mIDH1 cells vs. CQ treated WT-IDH1 cells) (Figures 7A-J). In this assay, CQ inhibits autolysosome formation allowing autophagosomes to form and accumulate within the cell. Together, these results indicate that glioma cells harboring IDH1^R132H^ have an augmented autophagy flux when compared with WT-IDH1 glioma cells.

**Figure 7:**
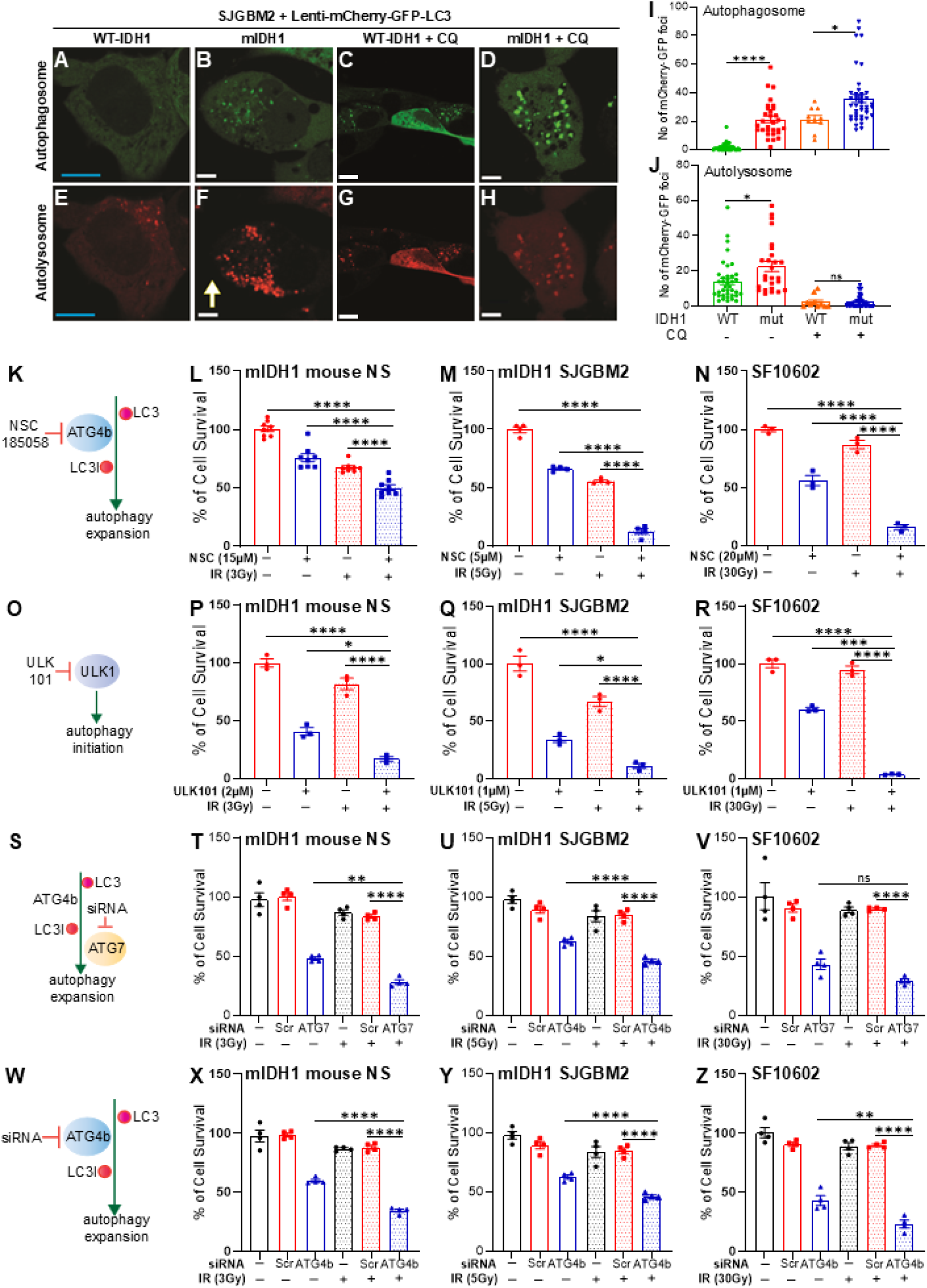
mIDH1 increases autophagy flux and in vitro inhibition of autophagy restores radiosensitivity in mIDH1 glioma cells. (**A-H**) Representative images of GFP and mCherry fluorescence in WT and mIDH1 human gliomas cells, SJGBM2, in basal condition or treated with 10 µM chloroquine (CQ). Scale Bar: 10 µm. (**I-J**) Quantification of number of (I) mCherry-GFP foci (indicating autophagosome formation) or (J) mCherry only foci (indicating autolysosome formation) in WT and mIDH1 SJGBM2 cells, in basal condition or treated with 10 µM chloroquine (CQ). Bar graphs represent the average of total foci per cell means ± SEM (n ≤ 10 replicates). \**P* < 0.05; \*\**P* < 0.01 \*\*\**P* < 0.001; \*\*\*\**P* < 0.0001; ns = not significant; t test. (**K**) NSC-185058 compound inhibits ATG4b, which is involved in LC3 processing during autophagosome expansion. (**L-N**) Cell viability assay performed in (L) mIDH1 mouse and (M) human glioma cells, SJGBM2 and (N) SF10602 treated with indicated doses of NSC-185058 and ionizing radiation (IR). (**O**) ULK101 compound inhibits ULK1 during autophagy initiation. (**P-R**) Cell viability assay performed in (P) mIDH1 mouse and (Q) human glioma cells, SJGBM2 and (R) SF10602 treated with indicated doses of ULK101 and IR. Results are expressed in percent of RLU in relation with the untreated control (100 %). \**P* < 0.05; \*\**P* < 0.01 \*\*\**P* < 0.001; \*\*\*\**P* < 0.0001; One-way ANOVA followed by Tukey’s test (n=4 technical replicates). (**S**) ATG7 inhibition by siRNA (siATG7) during autophagy initiation. (**T-V**) Cell viability assay performed in (T) mIDH1 mouse and (U) human glioma cells, SJGBM2 and (V) SF10602 treated with siATG7, and IR. Results are expressed in percent of cell survival in relation with the untreated control (100 %). (**W**) ATG4 inhibition by siRNA (siATG4) during autophagy initiation. (**X-Z**) Cell viability assay performed in (X) mIDH1 mouse and (Y) human glioma cells, SJGBM2 and (Z) SF10602 treated with siATG7, and IR. Results are expressed in percent of cell survival in relation with the untreated control (100 %). \**P* < 0.05; \*\**P* < 0.01 \*\*\**P* < 0.001; \*\*\*\**P* < 0.0001; One-way ANOVA followed by Tukey’s test (n=4 technical replicates).

### Inhibition of autophagy sensitizes mIDH1 glioma cells to ionizing radiation

Autophagy has been associated with DNA-damage response activation and resistance to ionizing radiation (IR) in tumor cells (57). We have previously reported that mIDH1 gliomas are resistant to radiotherapy. We hypothesized that the inhibition of autophagy will impair a proper response against a DNA damage insult in mIDH1 glioma cells, making them more sensitive to IR treatment. To test this hypothesis, we performed a cell viability assay in glioma cells harboring mIDH1, treated with multiple autophagy inhibitors in presence or absence of irradiation (Figures 7K-R). We first tested the effect of NSC185058, which targets ATG4b (Figure 7K) (53). Treatment with NSC185058 or irradiation alone decreased IDH1 glioma cells viability compared to vehicle (Figures 7L-N). However, combination treatment of NSC185058 with IR dramatically decreased mIDH1 glioma cells viability when compared to control or monotherapy (mIDH1 mouse NS: *P* < 0.0001, Figure 7L; SJGBM2: *P* < 0.001, Figure 7M; SF10602: *P* < 0.0001, Figure 7N). We then tested whether upstream inhibition of the autophagy pathway would be more potent in mIDH1 as a monotherapy. We used ULK101 which inhibits the initial steps of autophagy by targeting ULK1 (Figure 7O). Monotherapy with ULK101 significantly decreased cell viability in mIDH1 glioma cells (Figures 7P-R) when compared with control and IR alone (*P* < 0.0001 in all cells). However, combination therapy of IR and ULK101 resulted in a more dramatic decrease in mIDH1 glioma cells viability when compared to control and monotherapy (mIDH1 mouse NS: *P* < 0.05; SJGBM2: *P* < 0.05; SF10602: *P* < 0.001) or IR alone (mIDH1 mouse NS: *P* < 0.0001; SJGBM2: *P* < 0.001; SF10602: *P* < 0.0001) (Figures 7P-R). In addition, we evaluated the impact of the molecular inhibition of autophagy on glioma cells’ radio-responses through siRNAs (Supplemental Table 3) against autophagy-related genes ATG7 and ATG4b (Figures 7S-Z) and through the knockdown of ATG7 in mouse glioma cells. Using siRNAs (Supplemental Table 3) against autophagy-related genes ATG7 (Figure 7S), ATG4b (Figure 7W), we assessed the viability of both mouse and human glioma cells treated with radiation. We examined the impact of siRNA-ATG7 (Figures 7T-V) and siRNA-ATG4b (Figures 7X-Z) in the presence or absence of IR. Our observations revealed a significant reduction in viability across all tested cells when treated with a combination of siRNAs targeting ATG7 or ATG4b (mIDH1 mouse NS: *P* < 0.0001; SJGBM2: *P* < 0.0001; SF10602: *P* < 0.0001).

To validate effective silencing of ATG4b, ATG7, and ATG9b following siRNA treatment, mIDH1 mouse NS cells were transfected with the respective siRNAs at a concentration of 100nM (Supplemental Figure 27). Western blot analysis revealed significant reduction in protein expression-2.65-fold for ATG4b (*P* < 0.0001), 2.59-fold for ATG7 (*P* < 0.0001), and 2.06-fold for ATG9b (*P* < 0.0001) following 48hrs of siRNA treatment (Supplemental Figure 27).

To validate the results obtained using NPA/NPAI NS, using NRAS-independent genetically engineered mouse models, we proceeded to evaluate the impact of the molecular inhibition of autophagy on glioma cells’ radio-responses (Supplemental Figure 28-29). Using siRNAs ATG4b (Supplemental Figure 28A-B) and ATG7 (Supplemental Figure 28C-D), we assessed the viability of two mIDH1 murine glioma NS (RPAI/CPAI) treated in combination with radiation. Our results showed a significant reduction in viability across all cells when treated with IR in combination with siRNAs targeting ATG7 or ATG4b (RPAI mIDH1 mouse NS: P < 0.0001; CPAI mIDH1 NS: P < 0.0001). We next evaluated the impact of inhibiting the autophagy related gene, ATG9b, by using siRNAs (Supplemental Figure 29). Cell viability was assessed in both human (SJ-GBM2 mIDH1) and murine glioma (mIDH1 NPAI, mIDH1 CPAI, and mIDH1 RPAI) cells treated with ATG9b siRNAs, both in the presence and absence of IR. Silencing of ATG9b in mIDH1 human glioma cells (Supplemental Figure 29A) and mIDH1 mouse NS models (Supplemental Figure 29B–D) significantly reduced cell viability when combined with IR, compared to siRNA treatment alone (SJ-GBM2 mIDH1: *P* < 0.0001; CPAI mIDH1 mouse NS: *P* < 0.0001; RPAI mIDH1 NS: *P* < 0.0001; NPAI mIDH1 NS: *P*< 0.0001). Collectively, these findings indicate that molecular inhibition of autophagy enhances the radiosensitivity of mIDH1 murine and human glioma cells. In addition, we used a shRNA against *ATG7* to generate an autophagy deficient mouse glioma cell (Supplemental Figure 30). Like transient autophagy inhibition using siRNA (against ATG7, ATG4b, or ATG9b), mIDH1 autophagy deficient cells (mIDH1-ATG7KD) showed significant radiosensitization when compared with mIDH1 cells in which the autophagy pathway was not inhibited (P < 0.0001) (Supplemental Figure 31). The impact of concurrently inhibiting autophagy and applying radiation was assessed in both mouse (Supplemental Figure 32) and human glioma cells (Supplemental Figure 33) harboring mIDH1. Our observations revealed a notable shift in cytokine profiling, with a significant increase in cytokine concentration observed in mIDH1 glioma cells (Supplemental Figures 32-33; *P* < 0.01) upon combining autophagy inhibition with radiation. These findings suggest the potential of the combined treatment to elicit immune responses in mIDH1 glioma cells. In summary, our results collectively propose that autophagy inhibition not only heightens the sensitivity of mIDH1 glioma cells to radiotherapy but also modulates immune responses.

### Molecular inhibition of autophagy prolongs the survival of mIDH1 glioma-bearing mice

To study the impact of autophagy inhibition on mIDH1 tumor in vivo, we targeted the autophagy pathway via two approaches: i) a genetically engineered sleeping beauty model that expresses shRNA against *ATG7* (sh*ATG7*; Figure 8A and Supplemental Figures 30, 34); and ii) pharmacological inhibition of autophagy pathway using Synthetic protein nanoparticles (SPNPs) loaded with a siRNA targeting ATG7 (ATG7i-SPNP; Figure 8A and Supplemental Figures 35-36). To ensure ATG7i-SPNPs would impact glioma cell survival when loaded into SPNPs, we measured cell viability in both mouse and human mIDH1 glioma cells (Supplemental Figures 37-38). We also evaluated the impact of autophagy inhibition on the radiosensitivity of glioma cells using ATG7i-SPNPs (Supplemental Figure 37). Cell viability was assessed in mIDH1 glioma murine cell models NPAI, CPAI, and RPAI treated with ATG7i-SPNPs, in the presence or absence of IR. Silencing of ATG7 using SPNPs significantly reduced cell viability when combined with IR in all cells tested versus ATG7i-SPNPs treatment alone (RPAI mIDH1: *P* < 0.0001; NPAI mIDH1: *P* < 0.001; CPAI mIDH1: *P* < 0.0001; Supplemental Figure 37). These findings suggest that using ATG7i-SPNPs to inhibit autophagy enhances the radiosensitivity of mIDH1 glioma cells. We also evaluated the effect of autophagy inhibition on the radiosensitivity of human glioma and patient-derived endogenous mIDH1 expressing SF10602 cells using ATG7i-SPNPs (Supplemental Figure 38). Silencing of ATG7 significantly reduced cell viability when combined with IR versus ATG7i-SPNP treatment alone (*P* < 0.0001). These results indicate that inhibiting autophagy through ATG7i-SPNPs enhances the sensitivity of both mouse and human mIDH1 glioma cells to IR.

**Figure 8:**
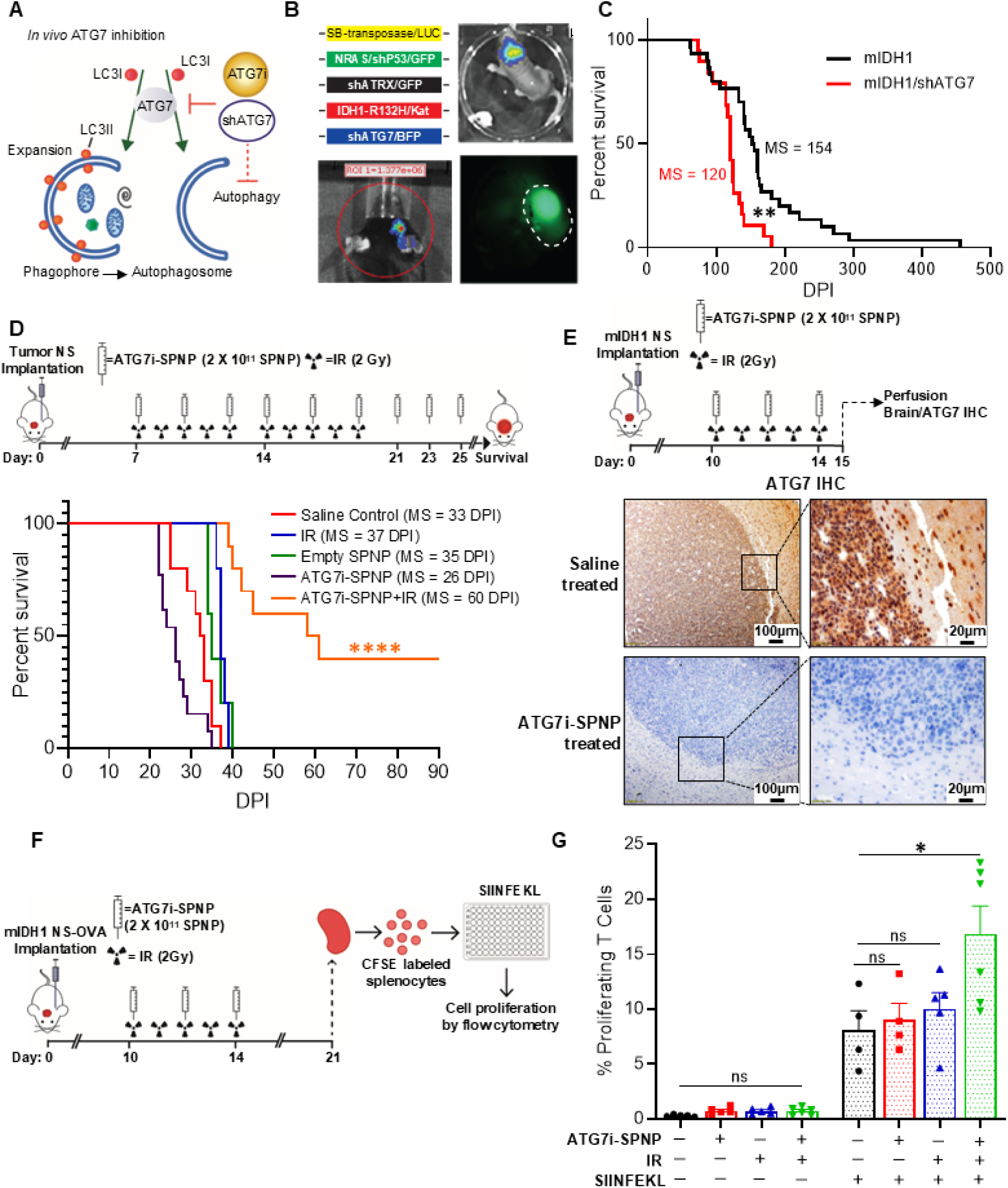
In vivo inhibition of autophagy radiosensitizes mIDH1 gliomas and enhances survival. (**A**) SB plasmid encoding shRNA targeting ATG7 (shATG7) and iRGD SPNPs delivering siRNA (ATG7i) developed for ATG7 inhibition in vivo. (**B**) Autophagy deficient mIDH1 mouse gliomas generated using SB transposon system with plasmids encoding SB-transposase-LUC, NRAS/shP53/GFP, shATRX-GFP, IDH1^R132H^/Katushka, and shATG7/CFP. Transfection efficiency monitored by luminescence. Symptomatic mice brain tumors expressing fluorescent proteins identified macroscopically. (**C**) Kaplan-Meier survival analysis for mice bearing mIDH1 (n=30) or mIDH1 + shATG7 (n=19) gliomas (*\*\*\*\*P*<0.0001, Mantel-Cox test). MS=median survival. (**D**) Autophagy inhibition effect using ATG7i-SPNP on radio-response in mIDH1 glioma model. Adult mice implanted with 50,000 mIDH1 NS (Day 0). At 7 DPI, animals split into groups: (i) untreated saline control; (ii) 9 doses of 2×10^11^ empty SPNPs every other day; (iii) 2 Gy/day; (iv) 9 doses of 2×10^11^ ATG7i-SPNP every other day; (v) 9 doses of 2×10^11^ ATG7i-SPNP every other day and 2 Gy/day (days 7-16). Kaplan-Meier survival analysis performed for each experimental group (*\*\*\*P*<0.001, Mantel-Cox test). (**E**) mIDH1 glioma NS implanted mice treated with ATG7i-SPNP were subsequently perfused at 14 DPI and analyzed for IHC. Representative Immunohistochemical (IHC) staining of 5 μm paraffin-embedded brain sections from saline and ATG7i–SPNP treated groups, stained for ATG7. Low magnification (10×) panels (black scale bar = 100 μm), High magnification (40×) panels (black scale bar = 20 μm) indicate positive staining for the areas in the low-magnification panels. (**F**) Experimental figure where mice were implanted with mIDH1-OVA NS and either received (i) no treatment, (ii) 2 Gy radiation for 5 days, (iii) ATG7i-SPNP for 3 doses, or (iv) both radiation and ATG7i-SPNP combination. Splenocytes were harvested on day 21, CFSE labelled, and cultured with SIINFEKL for 24hrs before proliferation analysis. (**G**) CD8 T cell proliferation, as a measured by CFSE dilution, of splenocytes harvested from the treated mIDH1-OVA NS implanted mice. Bars (mean ± SEM) represent proliferating CD8^+^ T cell percentage within treatment groups (n = 4-6 replicates; one-way ANOVA followed by Tukey’s test. \**P*<0.01; ns=non-statistically significant).

The genetically engineered autophagy deficient mouse mIDH1 glioma model was developed using the Sleeping Beauty (SB) Transposon System through a combination of SB-transposase/LUC; *NRAS*/sh*P53*/GFP; sh*ATRX*/GFP; *IDH1^R132H^*/Kat and sh*ATG7*/CFP plasmids (Figure 8B). We found that mice with *ATG7* KD had a significantly lower median survival (MS) when compared with the control (154 vs. 120 days post-plasmid injection (DPI); *P* < 0.001) (Figure 8C). We then proceeded to assess the impact of autophagy inhibition in combination with radiation treatment using our cell implantable mouse mIDH1 glioma model (15). We intracranially implanted mIDH1 glioma NS in mice, and 7 days later the animals were split in five groups: i) saline control; ii) treated with empty-SPNPs; iii) treated with IR; iv) ATG7i-SPNP; and v) treated with IR + ATG7i-SPNP at indicated doses and schedule (Figure 8D and Supplemental Table 4). In the genetically engineered mouse mIDH1 glioma model, autophagy inhibition alone significantly decreased MS (*P* < 0.05) when compared with the control group (Figure 8D and Supplemental Table 4). However, IR + ATG7i-SPNP led to increased MS (MS = 60 DPI; *P* < 0.001) when compared with all other experimental groups (Figure 8D and Supplemental Table 4). We observed that the combined treatment prolonged the survival of mice long-term (> 90 DPI in 40% of the treated animals). To ensure that ATG7 was suppressed in vivo in response to ATG7i-SPNP treatment, we performed immunohistochemistry on mIDH1 glioma bearing mice treated with ATG7i-SPNP (Figure 8E, Supplemental Figure 39). We found that mIDH1 glioma-bearing mice treated with ATG7i-SPNPs (Figure 8E, lower panel) exhibited no ATG7 expression when compared to saline-treated mIDH1 glioma bearing mice (Figure 8E, upper panel).

Long-term survivors from the IR + ATG7i-SPNP treatment groups were rechallenged with mIDH1 mouse NS in the contralateral hemisphere (Supplemental Figure 40). These animals remained tumor free without further treatment, whereas control mice implanted with glioma cells succumbed due to tumor burden (MS = 33 DPI; P ≤ 0.0001) (Supplemental Figure 40). The animals were sacrificed at 60 DPI after tumor rechallenge, mice showed no evidence of microscopic intracranial tumor. Together, these results suggest the development of anti-glioma immunological memory in mIDH1 glioma rechallenged animals previously treated with IR + ATG7i-SPNP. None of these treatments produced hepatic alterations compared with the control group (Supplemental Figures 41-42).

We then aimed to study the anti–mIDH1 glioma-specific immune response elicited by IR + ATG7i-SPNP therapy, using a T cell proliferation assay. Mice bearing mIDH1 tumors harboring a surrogate tumor antigen, OVA, were treated with saline, IR, ATG7i-SPNP, or IR + ATG7i-SPNP (Figure 8F). Mice were euthanized 14 days after the last treatment dose (30 DPI) and the spleen were removed and processed for flow cytometry analysis. Splenocytes were fluorescently labeled with 5- and 6-CFSE and stimulated for 4 days with 100 nM SIINFEKL (Figure 8F), where T cell proliferation was measured as the reduction of CFSE staining in the CD45+/CD3+/CD8+ population. After SIINFEKL treatment, we observed that T cell proliferation was significantly increased (*P* < 0.05) in mIDH1 glioma-bearing mice treated with IR + ATG7i-SPNP (Figure 8G and Supplemental Figure 43) when compared with the control group. This indicates that the combined treatment was able to induce a systemic anti–mIDH1 glioma-immune response associated with an enhancement in T cell proliferation. In our investigation, we conducted a comprehensive assessment of tumor growth using histological analysis via H&E staining, along with immunohistochemical markers CD3 (reflecting infiltrating T cells) and CD68 (indicating an inflammatory response or the presence of tumor-associated macrophages) in mice bearing mIDH1 tumors subjected to treatment with ATG7i-SPNP + IR. The results revealed a noteworthy reduction in tumor size concomitant with a decrease in the presence of CD3- and CD68-positive cells following the combined treatment, as illustrated in Supplemental Figures 44-45. Taken together, these findings indicate a potent anti-tumor effect, suggesting a multifaceted impact on both T cell infiltration and the inflammatory microenvironment.

## Discussion

Metabolic reprogramming is a hallmark alteration observed in glioma patients harboring IDH1^R132H^ mutation (58), however the biological consequences and its therapeutic implications remain to be fully elucidated. In this study, we show that in mIDH1 gliomas, in the context of *ATRX* and *TP53* inactivating mutations, autophagy is upregulated in conjunction with functional disruption of mitochondrial activity. Our RNA-seq data displays an upregulation of several autophagy related genes and a downregulation of genes related with mitochondrial metabolism. The downregulation of mitochondrial activity and oxidative phosphorylation is evident in mIDH1 glioma cells, as revealed by the signaling pathway enrichment analysis data. Consistently, mass spectrometry-based metabolomics analysis revealed a marked alteration in the mIDH1 glioma metabolic profile and an alteration of the TCA cycle, with an accumulation of TCA cycle metabolites, suggesting impaired utilization of this pathway. In line with our findings, previous studies suggested that IDH1 mutation modulates TCA cycle, mitochondrial metabolism, and oxidative stress pathways (16, 20, 23, 59, 60). The presence of 2HG in glioma cultures reduces both oxidative phosphorylation and ATP levels in glioma cells (61). This phenomenon of metabolic alterations is also observed in our genetically engineered mIDH1 glioma model, which shows an upregulation of DNA-damage responses (DDR) (15). In addition, the enhancement of DDR has been related with metabolic alterations present in mIDH1 gliomas (16). This supports our results showing that the metabolic alterations in mIDH1 glioma cells are involved in the TCA cycle and mitochondrial dysfunctions.

Autophagy is activated in response to mitochondrial dysfunction, serving as a mechanism for cell protection and survival (62–64). Our ChIP-seq data confirms that epigenetic reprograming of mIDH1 gliomas impacts the expression of several *ATG* genes as observed in both mouse and human mIDH1 glioma cells. Our data, obtained by examination of OCR and ECAR, revealed that mitochondrial and glycolytic activity are reduced in mIDH1 glioma cells. These results functionally support our molecular data, indicating a mitochondrial dysfunction along with an energetic state as encountered in slow proliferative cells. It has been suggested that autophagy may provide nutrients necessary to meet bioenergetic demands during transition from slower proliferative state to an energy dependent cell state (65). Furthermore, it has been demonstrated that mitophagy, the mitochondrial autophagy, can be induced by redox mitochondrial agents in cancer cells (63). Our results demonstrate that human and mouse mIDH1 glioma cells present morphological characteristics indicative of the mitochondrial fission process, which is tightly related with autophagy/mitophagy (50). Using an autophagy flux assay and WB to detect activation of *ATG* genes, the conversion of LC3-I to LC3-II, which is one of the distinctive hallmarks of autophagy (66), we demonstrated that autophagy activity is functionally increased in mouse and human mIDH1 glioma cells. We show that the inhibition of IDH1^R132H^ reverts this phenotype in patient-derived mIDH1 glioma cells, suggesting that the enhancement of autophagy is 2HG dependent. Our study describes epigenetic mechanisms involved in this phenotype, where histone methylation plays a key role in metabolic reprogramming and activation of autophagy.

According to our results, autophagy could serve as a cell protection and survival mechanism in mIDH1 gliomas. It has been described that autophagy can have an impact on cancer therapeutic responses (67–72). We evaluated the impact of autophagy inhibition in response to radiation in vitro and in vivo. Our results indicate that autophagy inhibition sensitizes mIDH1 glioma cells to IR treatment, decreasing cell viability of mouse glioma NS and human glioma cells. This is supported by the fact that autophagy can induce DNA-damage response activation, thus inducing resistance to IR in tumor cells (57). We generated a genetically engineered autophagy deficient mIDH1 mouse glioma model, which displayed decreased median survival when compared with the control mIDH1 glioma mouse model.

These results are consistent with known roles autophagy has played in contributing to a tumorigenic environment, such as the protective role of autophagy in the presence of DNA damage (36) and its role in maintaining chromosomal stability (73–75). Our results show an upregulation of *UVRAG*, an *ATG* gene involved in chromosomal stability, which is in line with an increased DDR reported in mIDH1 gliomas (15, 16). Conversely, this phenomenon could increase cell protection against DNA damage assault and contribute to radio-resistance. Using our cell implantable mIDH1 glioma model (15), we evaluated the therapeutic impact of autophagy inhibition using SPNPs loaded with siRNA targeting ATG7 (ATG7i-SPNP) in combination with IR. This combination increased median survival and elicited 40% of long-term tumor-free survivors in mIDH1 glioma bearing mice, demonstrating that autophagy inhibition sensitizes mIDH1 tumors to radiotherapy. Also, we did not observe overt toxicities as the treated animals were examined twice daily by expert veterinary staff at the University of Michigan’s Institutional Animal Care and Use Committee. We evaluated the effects of the ATG7i-SPNP in combination with IR treatment in long-term survivors via histopathological analysis of brains and livers. Our results show that this treatment did not cause any alterations in the brain or liver architecture. Normal liver and kidney function in the treated animals was evidenced by the mouse serum biochemistry analysis (Supplemental Figures 41-42, 44-45).

In addition, our re-challenge experiment displayed a 100% survival rate of mIDH1 animals that were previously treated with IR in combination with ATG7i-SPNP. These results suggest that the combined therapeutic approach induces anti-tumor immunological memory (22). This was further confirmed as we observed increased surrogate tumor antigen (Ovalbumin) specific T cell proliferation in mIDH1 glioma animals treated with IR and ATG7i-SPNP. Although, Ovalbumin does not naturally occur in humans or other mammals, when it is expressed in tumor cells, it creates a “surrogate” tumor antigen. This enabled the detection of tumor-antigen specific T cells in the spleen from the treated animals. Using SIINFEKL (cognate ovalbumin peptide), we are able to demonstrate that the ATG7i-SPNP+IR treatment led to effector T cells that can recognize and proliferate when stimulated by the surrogate tumor antigen, “SIINFEKL”.

Our work demonstrates that autophagy activation leads to radioresistance in patient-derived mIDH1 low grade glioma (LGG) cell cultures. Furthermore, we show that when radiation was delivered in combination with autophagy inhibition (ATG7i-SPNP), the mIDH1 gliomas become radiosensitive which leads to long-term survival and anti-tumor immunological memory (Figure 8). However, evaluating these responses in patient-derived glioma models is limited because patient-derived cells do not form tumors in immunodeficient mice. Our past experiences with genetically engineered animal models, which recapitulate critical features observed in glioma patients and molecular databases (12, 15, 22, 76), provide confidence that our results are relevant to a better understanding of the biological significance of metabolic reprograming observed in mIDH1 gliomas and the involvement of autophagy in mediating therapeutic responses. Our results apply to a particular mIDH1 molecular subtype, which harbors concomitant inactivating mutations in *TP53* and *ATRX*, recently classified by the WHO as mIDH1 astrocytomas (77). This genotype was included in all in vitro, ex vivo, and in vivo experiments presented. Although *NRAS* is not a common mutation found in gliomas, our *RAS* independent genetically engineered mouse models as well as human mIDH1 glioma cells, obtained from surgical biopsies, have successfully replicated the results obtained from our *NRAS-G12V* driven mIDH1 genetically engineered mouse model.

The molecular, functional, and pre-clinical data of mIDH1 gliomas indicate that the epigenetic reprogramming on mitochondrial metabolic activity, which is downregulated when autophagy is enhanced, influences therapeutic responses. Through this study, we have shown that inhibiting autophagy not only radiosensitizes in vivo mIDH1 glioma tumors but also leads to increased survival, long-term survivors, and anti-tumor immunity (Figure 8); thus, providing a promising and novel therapeutic combination treatment that could be further developed and implemented in mIDH1 glioma patients.

### Study Design

To study the impact of autophagy in the context of IDH1^R132H^ mutant gliomas, with *TP53* and *ATRX* inactivating mutations, we previously generated a genetically engineered animal model injecting SB plasmids encoding NRAS G12V, shp53, and shATRX, and with or without IDH1^R132H^ into the lateral ventricle of neonatal mice (15). Sample size and any data inclusion/exclusion were defined individually for each experiment. We also used an animal model previously generated by intracranial implantation of glioma NS (WT-IDH1 and mIDH1) derived from our genetically engineered animal model to test therapeutic responses (15). Furthermore, we used human glioma cells derived from patients harboring IDH1^R132H^, in the context of *TP53* and *ATRX* inactivating mutations, to confirm the results obtained from our animal models. The number of replicates are reported in the figure legends. Our studies were not randomized. We performed blinding for quantitative IHC scoring. All RNA-seq and ChIP-seq data were deposited in public databases as is indicated in the respective sections. Materials and Methods are detailed in the Supplemental Materials.

### Genetically engineered mutant IDH1 glioma model

All animal studies were conducted according to guidelines approved by the IACUC at the University of Michigan. All animals were housed in an AAALAC accredited animal facility; and they were monitored daily. Studies did not discriminate sex, and both male and females were used. The strains of mice used in the study were C57BL/6 (The Jackson Laboratories, 000664). Please see additional details in the Supplemental Materials and Methods.

### Generation of iRGD Synthetic Protein Nanoparticle (SPNP) with siRNA against *ATG7*

The ATG7 Mouse siRNA Oligo Duplex (Locus ID 74244) (Origene, SR427399) was used to generate iRGD SPNP. Please see additional details including SPNP formulation, fabrication, collection and processing, and characterization in the Supplemental Materials and Methods.

### Statistical analysis

All quantitative data are presented as the mean ± SEM from at least three independent samples. ANOVA and two-sample *t* tests were used to compare continuous outcomes between groups. Survival curves were analyzed using the Kaplan-Meier method and compared using Mantel-Cox tests; the effect size is expressed as median survival (MS). Differences were considered significant if *P* < 0.05. All analyses were conducted using GraphPad Prism software (version 6.01), SAS (version 9.4, SAS Institute), or R (version 3.1.3). The statistical tests used are indicated in each figure legend.

### Study Approval

All animal studies were conducted according to guidelines approved by the IACUC of the University of Michigan (protocols PRO00011290 and PRO00011168). All animals were housed in an AAALAC-accredited animal facility and were monitored daily. Studies did not discriminate by sex; both male and female mice were used. The strains of mice used in the study were C57BL/6 (the Jackson Laboratory, strain no. 000664) and CDKN2A-KO mice (Frederick National Library for Cancer Research strain no. 01XB1).

## Data and materials availability

There are no limitations on data availability. The ChIP-Seq dataset generated in this study can be acquired through NCBI’s Gene Expression Omnibus (GEO) with identifier GSE99806. The RNA-Seq dataset generated in this study can be acquired through NCBI’s Gene Expression Omnibus (GEO) with identifiers GSE94902, GSE94974, and GSE94975. The scRNA-Seq dataset generated in this study can be acquired through NCBI’s Gene Expression Omnibus (GEO) with identifier GSE261042. All other data generated within this study are available via the supplemental data files; cells and plasmids will be freely distributed upon request from the corresponding author, Maria G Castro. All individual data points can be found in the Excel file “Supporting Data Values”.

## Author contributions

Conceptualization: MGC, FJN. Methodology: All: FJN, KB, AAM, AM, CET, ZZ, SR, MRS, JAPA, JZ, AT, PK, SVC, MBGF, AC, MSA, BLM, SMF, ZCN, HSH, PS, TQ, MAS, ML, JDW, SYC, PRL, JL, CAL, MGC. Investigation: All: FJN, KB, AAM, AM, CET, ZZ, SR, MRS, JAPA, JZ, AT, PK, SVC, MBGF, AC, MSA, BLM, SMF, ZCN, HSH, PS, TQ, MAS, ML, JDW, SYC, PRL, JL, CAL, MGC. Visualization: All: FJN, KB, AAM, AM, CET, ZZ, SR, MRS, JAPA, JZ, AT, PK, SVC, MBGF, AC, MSA, BLM, SMF, ZCN, HSH, PS, TQ, MAS, ML, JDW, SYC, PRL, JL, CAL, MGC. Funding acquisition: MGC, PRL, CAL, JL. Project administration: MGC. Supervision: MGC, PRL, CAL, JL. Writing – original draft: All: FJN, KB, AAM, AM, CET, ZZ, SR, MRS, JAPA, JZ, AT, PK, SVC, MBGF, AC, MSA, BLM, SMF, ZCN, HSH, PS, TQ, MAS, ML, JDW, SYC, PRL, JL, CAL, MGC. Writing – review & editing: All: FJN, KB, AAM, AM, CET, ZZ, SR, MRS, JAPA, JZ, AT, PK, SVC, MBGF, AC, MSA, BLM, SMF, ZCN, HSH, PS, TQ, MAS, ML, JDW, SYC, PRL, JL, CAL, MGC. Co-First Authors: FJN is listed first because he was part of the conceptualization, developed the genetically engineered mouse model, and he also performed the initial experiments shown in this manuscript. All other co-first authors (KB, AAM, AM, CET, ZZ) were placed in alphabetical order and contributed equally to the experiments presented, writing and editing, data analysis, and preparation of new figures for publication.

## Acknowledgments

We thank Dr. J. Costello (UCSF) for providing the mIDH1 human glioma cells SF10602, which were a gift from the Dabbiere Family and funded by NIH/NCI R01CA244838; the COG Repository at the Health Science Center for providing the human glioma cells SJGBM2; and Dr. J. Ohlfest (University of Minnesota, deceased) for providing the SB model plasmids. This work was supported by the NIH/National Institute of Neurological Disorder & Stroke (NINDS) Grants R37-NS094804, R01-NS122165, and R21-NS123879-01, the National Cancer Institute (NCI) Cancer Center Support Grant 2P30CA46592, and the Rogel Cancer Center Faculty Scholar Award (to MGC); NIH/NINDS Grant R01-NS124167 (to MGC and JL); NIH/NINDS Grants R01NS122234 and R01-NS122378 (to PRL); The Pediatric Brain Tumor Foundation, Leah’s Happy Hearts Foundation, Ian’s Friends Foundation, Chad Tough Foundation, and Smiles for Sophie Forever Foundation (to MGC and PRL); NCI Grants R01CA248160 and R01CA244931 (to CAL); and the National Science Foundation Graduate Research Fellowship Grant DGE 1256260 (to AM).

