## Supplementary material for "Epigenetic Reprogramming of Autophagy Leads to Uncovering a Novel Therapeutic Target for Mutant IDH1 Astrocytomas": Supp

### Supplemental Materials and Methods

#### Genetically engineered mutant IDH1 glioma model

The wild type (WT)-IDH1 and mIDH1 glioma models used in this study were generated previously for our group using SB Transposon System (15). The plasmid used to generate those models are: (i) SB transposase and LUC (pT2C-LucPGK-SB100X, henceforth referred to as SB/Luc), (ii) a constitutively active mutant of NRAS, NRAS-G12V (pT2CAG-NRASV12, henceforth referred to as NRAS), (iii) a short hairpin against p53 (pT2-shp53-GFP4, henceforth referred to as shp53), (iv) a short hairpin against ATRX (pT2-shATRX53-GFP4, henceforth referred to as shATRX), and (v) mutant IDH1$^{R132H}$ (pKT-IDH1$^{R132H}$-IRES-Katushka, henceforth referred to as mIDH1). Neonatal P01 C57BL/6 mice were used in all experiments. The genotype of SB- generated mice involved these combinations: (i) NRAS, shP53, and shATRX (WT-IDH1) and (ii) NRAS, shp53, shATRX, and IDH1$^{R132H}$ (mIDH1). Mice were injected according to a previously described protocol (78). Plasmids were mixed in mass ratios of 1:2:2:2 or 1:2:2:2:2 (20 μg plasmid in a total of 40 μL plasmid mixture) with in vivo-jetPEI (Polyplus Transfection, 201-50G) (2.8 μL per 40 μL plasmid mixture) and dextrose (5% total) and maintained at room temperature for at least 15 min prior to injection. The lateral ventricle (1.5 mm AP, 0.7 mm lateral, and 1.5 mm deep from lambda) of neonatal mice (P01) was injected with 0.75 μL plasmid mixture (0.5 μL/min) that included: (1) SB/Luc, (2) NRAS, (3) shp53, (4) with or without shATRX, and (5) with or without IDH1$^{R132H}$. To monitor plasmid uptake in neonatal pups, 30 μL of luciferin (30 mg/mL) was injected s.c. into each pup 24-48h after plasmid injection. In vivo bioluminescence was measured on an IVIS Spectrum (Perkin Elmer, 124262) imaging system. For the IVIS spectrum, the following settings were used: automatic exposure, large binning, and aperture f=1. For in vivo imaging of tumor formation and progression in adult mice, 100 μL of luciferin solution was injected i.p. and mice were then anesthetized with oxygen/isoflurane (1.5-2.5% isoflurane). To score luminescence, Living

Image Software Version 4.3.1 (Caliper Life Sciences) was used. A region of interest (ROI) was defined as a circle over the head, and luminescence intensity was measured using the calibrated unit's photons/s/cm2/sr. Multiple images were taken over a 25min period following injection, and maximal intensity was reported.

For survival studies, animals were monitored daily for signs of morbidity, including ataxia, impaired mobility, hunched posture, seizures, and scruffy fur. Animals displaying symptoms of morbidity were intracardially perfused using Tyrode's solution, followed by fixation with 4% paraformaldehyde (PFA) in PBS.

**Alternative genetically engineered mouse glioma models**

In CPA/CPAI models, the genetic lesions were incorporated into neonatal CDKN2A homozygous knocked-down (CDKN2A$^{-/-}$) mice (12). CDKN2A$^{-/-}$ mice were obtained from the Frederick National Library for Cancer Research as frozen embryos. Embryos were implanted into receptive adult B6 females by the Transgenic Animal Core of the University of Michigan and the obtained transgenic mice, which were CDKN2A heterozygous, were mated with B6 mice. The CDKN2A KO heterozygous mice of second generation were mated and the progeny obtained was genotyped to confirm the homozygous CDKN2A deletion. From this point, the CDKN2A$^{-/-}$ colony was maintained by mating CDKN2A$^{-/-}$ mice and monitored regularly by CDKN2A genotypification. The genotype of the genetically engineered mice involved these combinations: (i) CDKN2A deletion, TP53 and ATRX short hairpin knockdown, with or without IDH1-R132H. In the RPA/RPAI models, glioma was driven by platelet derived growth factor receptor alpha mutation (PDGFRA$^{D842V}$) to promote constituent activation of the MAPK pathway (18). These mouse glioma cells include ATRX and TP53 short hairpin knockdown, with or without endogenous express IDH1-R132H, to simulate the molecular genetic lesions that define the molecular features of astrocytoma.

**Primary glioma neurosphere cultures (NS)**

Mouse glioma neurosphere (NS) cultures were generated as previously described (15, 79). Briefly, brain tumors in mice were harvested at the time of euthanasia by intracardial perfusion with Tyrode's solution only. The tumor mass was dissociated using non-enzymatic cell dissociation buffer, filtered through a 70 µm strainer and maintained in neural stem cell medium DMEM/F12 with L-Glutamine (Gibco, 11320-033), B-27 supplement without Vitamin A (1X) (Gibco, 12587-010), N-2 supplement (1X) (Gibco, 17502-048), Penicillin-Streptomycin (100X) (10,000 IU Penicillin) (10,000 µg/mL Streptomycin) (Corning, Cellgro, 30-001-CI), and Normocin (1X) (Invivogen, ant-nr-1) at 37°C, 5% $CO_2$. Human-EGF (Peprotech, 100-15) and human-FGF (Peprotech, 100-18) were added twice weekly at 1 µL (20 ng/µL each stock) per 1 mL medium for a final concentration of 20 ng/mL. For reduced growth conditions, human-EGF and human-FGF were added twice weekly at 1 µL (20 ng/µL) per 3 mL medium for a final concentration of 6.6 ng/mL.

**RNA-seq and bioinformatics analysis**

RNA-seq was performed in collaboration with the University of Michigan sequencing core. All the RNA-seq datasets used to identify differential enrichments related with metabolism and autophagy were previously generated in our lab as detailed in (15) with public access. Briefly, RNA was isolated from tumor NS and 100 ng samples of purified RNA were sent for RNA Seq analysis. After passing all quality controls, Illumina HiSeq 2500/4000 fastq files were processed using the Tuxedo Suite for alignment and differential expression analysis. Genes and transcripts were identified as differentially expressed based on three criteria: test status = "OK", FDR $\leq$ 0.05, and fold change $\geq \pm$ 1.5 Gene Ontology (GO) enrichment analysis, which was performed using iPathwayGuide (http://www.advaitabio.com/ipathwayguide) separately for upregulated and downregulated genes. GO Biological Processes, selected for relevance to phenotype, were plotted in a horizontal bar graph using R. Gene set analysis (GSA) was performed using the R package GSA (http://statweb.stanford.edu/~tibs/GSA/), which implements a version of Gene Set Enrichment Analysis (GSEA) (80) with improved power statistics (81). GSA takes an expression matrix and a gene set collection as input. The Gene Set Knowledgebase (GSKB) data package (version 1.3.0) was downloaded from Bioconductor. GSKB is specific to mice and includes annotation from over 40 sources. GSA outputs significant gene sets (FDR $\leq$ 0.05), both negatively and positively correlated to phenotype, as well as normalized enrichment scores, and the lists of genes contributing to the significance of each gene set (FDR $\leq$ 0.05). For each list of genes found in a GSKB functional term we tested for enrichment in a small set of Gene Ontology terms, selected for relevance to the phenotype, using the hypergeometric test in R. After RNA-seq analysis, enrichment maps were generated using the Cytoscape platform with enrichment map; the designation of node color was amended to complement our RNA-seq expression color scheme of positively (red) and negatively (green) regulated pathways. The stringency of FDR was reduced to truly encapsulate the underlying biological ramifications of ATRX and IDH1 mutations in our transposon-mediated tumor formation model. Heatmaps were plotted with Heatmap2 package in R. RNA-seq datasets have been deposited in NCBI's Gene Expression Omnibus with identifiers GSE94902, GSE94974, and GSE94975.

**Bru-seq and bioinformatics analysis**

The Bru-seq dataset used in this work to compare transcriptional levels of genes related with metabolism and autophagy was previously generated as described in (15). The protocol used was described in (82). Nascent RNA labeling was performed for 30 min at 37°C with 2 mM 5-bromouridine in conditioned medium. The cells were then incubated for 6h at 37°C. At the completion of labeling, the cells were lysed in TRIzol, and the Bru-containing RNA was isolated using anti-BrdU antibodies conjugated to magnetic beads. The isolated RNA was converted into cDNA libraries and prepared for sequencing using the Illumina TruSeq RNA Library Preparation Kit v2 followed by deep sequencing to around 50 million single-end 50 nucleotide reads. Bioinformatics and data analysis pipeline was implemented using the q pipeline manager (http://sourceforge.net/projects/qppln-mngr/). The bioinformatics programs used, read mapping, genome annotation, and expression scoring were previously described (82), section 2.4. RPKM (reads per kilobase per million mapped reads) values were calculated for individual genes that were at least 300 bp long. For genes with lengths of 30 kb and less, RPKM values were calculated using read counts from the entire gene. For genes longer than 30 kb, an RPKM value was calculated using read counts from the first 30 kb downstream of the TSS. The R package DESeq35 was used to test differential expression of genes where the mean RPKM between samples was greater than 0.5. Significant changes in transcription initiation were defined as follows: adjusted p-values < 0.05; fold change < 1.5.

**ChIP-Sequencing and bioinformatics analysis**

For each immunoprecipitation (IP) performed, $1x10^6$ NS were collected, washed in HBSS (Gibco), and pelleted in low binding Corning Costar microtubes. A 95 µL aliquot of digestion buffer (50 mM Tris-HCl (Sigma Aldrich), 1 mM $CaCl_2$ (Sigma Aldrich), 0.2% Triton X-100 (Sigma Aldrich), pH 8.0) supplemented with 1:100 protease inhibitors (Sigma Aldrich) was added per $1x10^6$ NS and immediately pipetted gently up and down to lyse. Chromatin digestion was initiated by the addition of 5 µL of Micrococcal Nuclease (ThermoFisher Scientific at $3.83 \times 10^{-3}$ U/µL final concentration, for 12 minutes at 37°C, and stopped by the addition of 10X stopping buffer (110 mM Tris-HCl, 55 mM EDTA (Sigma Aldrich), pH 8). The lysate was then diluted with an equal volume of X2 RIPA buffer (280 mM NaCl (Sigma Aldrich), 5 mM EGTA (ThermoFisher Scientific), protease inhibitor diluted 1:100). At this stage 10% of the lysate volume was withdrawn as input. The remaining lysate was then precleared with 10 µL of Dynabeads (1:1 mixture of Protin A (ThermoFisher Scientific) and protein G (ThermoFisher Scientific)) and washed in TIP buffer (10 mM Tris-HCl, 1 mM EDTA, 140 mM NaCl, 1% Triton X-100, 0.1% SDS, 0.1% sodium deoxycholate, pH

8).). Samples were then incubated overnight at 4°C with 1 µg of anti-H3K4me3 (Diagenode, C15410003), anti-H3K27ac antibody (Diagenode), anti-H3K27me3 antibody (Diagenode), anti-H3K36me3 antibody (Diagenode) or anti-total mouse IgG antibody (Santa Cruz Biotech). Antibodies were validated prior to use with Active Mofit's MODified Histone Peptide Array (Active Motif). Immunoprecipitation was performed in the follow day by incubating the samples for 3 hours with 10 µL of Dynabeads (1:1 mixture of Protein A and Protein G (ThermoFisher Scientific)) in 1x RIPA, followed by washing steps with 1x RIPA (5 times, supplemented with 1:1000 dilution of protease inhibitor (Sigma Aldrich)), LiCl buffer (1 time, 250 mM LiCl (Sigma Aldrich, 10 mM Tris-HCl, 1 mM EDTA, 0.5% NP-40, 0.5% sodium deoxycholate, pH 8, supplemented with 1:1000 dilution of protease inhibitor), and finally TE buffer (1 time, protease inhibitor-free (Corning)). Immunoprecipitated chromatin was freed from histones and Dynabeads by incubating the samples for 1 hour, 37°C with proteinase K (Qiagen) at a final concentration of 0.50 mg/mL. Chromatin samples were then purified by the Qiagen QIAquick PCR Purification Kit (Qiagen) and eluted with 50 µL of the EB buffer provided in the kit. Libraries were prepared at the Epigenomics Core using the TruSeq ChIP Library Preparation Kit (Illumina). Input and IP DNA (5-10 ng) was blunt-ended and phosphorylated, followed by the addition of a single adenine nucleotide, prior to ligation with an adaptor duplex with a T overhang. Ligated products were size-selected by agarose gel electrophoresis, purified, and PCR-amplified for the final library preparation. Quality control was performed on each input/IP library generated using loci specific primers to verify that the original enrichment was retained in the final library. Input/IP libraries that pass this quality control were then sequenced on an Illumina NovaSeq 6000 platform.

For ChIP-Seq data analysis, the ChIP-seq raw data quality was assessed using FastQC 0.11.3, and the adaptors were trimmed using TrimGalore 0.4.4. ChIP-seq and input reads were aligned to the mouse reference genome (mm10) using bowtie2 (83) with default options, and the unique map reads were extracted for peak calling. The post-alignment quality control envisulaized were performed by Deeptools 2 3.5.1 (84). PePr (85) was used to identify the differential peaks between WT-IDH1 and mIDH1 groups using FDR < 0.05 and fold-change > 3 as the cutoffs. The identified peaks were annotated to genomic features 1-5 kb upstream of promoter, promoter, intron, exon, UTR, CDS, and intergenic (using Bioconductor *annotatr* package (86). As a comparison, the same number of peaks were randomly generated across the same chromosome and annotated the same way by *annotatr*. Gene set enrichment testing was performed on the identified differential peaks by Bioconductor package *ChIPenrich* (87) to detect enriched Gene Ontology Biological Process Terms.

**Cleavage Under Targets & Release Using Nuclease (CUT&RUN)**

CUT&RUN was performed using CUTANA ChIC/CUT&RUN kit (Epicypher #14-1048) following manufacturer's protocol. Briefly, $1 \times 10^6$ NS cells were coupled with Concanavalin A beads, permeabilized with 0.01% Digitonin and incubated overnight at 4°C with 0.5 μg target antibody in antibody buffer (H3K4me3 Antibody - SNAP-Certified™ for CUT&RUN, Epicypher #13-0041, H3K27ac Antibody, SNAP-Certified™ for CUT&RUN and CUT&Tag, Epicypher #13-0059). The following day, cells were first incubated for 10 min with micrococcal nuclease fused to proteins A and G (pAG-MNase), which was then activated by $CaCl_2$ addition. After 2 hours incubation at 4°C, the reaction was stopped with stop buffer and E. coli DNA was added as spike-in DNA. DNA was then isolated using SPRI beads and quantified with Qubit. Libraries were prepared using CUTANA™ CUT&RUN Library Prep Kit, following manufacturer's recommendations. Fragment size was detected using a Tape Station system and multiplexed libraries were sequenced on AVITI24 sequencer (Element Biosciences) at 150bp paired end reads. Data analysis was performed the same way as described in ChIP-Sequencing section.

**Human Single Cell RNA-seq**

Human glioma scRNA-seq was performed and analyzed as previously described (12). The data has been deposited in NCBI's Gene Expression Omnibus with identifier GSE. Briefly, primary mIDH1 human glioma cells, SF10602, cultured for 7 days, in presence or absence of specific $IDH1^{R132H}$ inhibitor AGI-5198. 3' single cell libraries were generated using the 10X Genomics Chromium Controller following the manufacturer's protocol for 3' V3.1 chemistry with NextGEM Chip G reagents (10X Genomics). Final library quality was assessed using the Tapestation 4200 (Agilent), and libraries were quantified by Kapa qPCR (Roche). Pooled libraries were subjected to 150 bp paired-end sequencing according to the manufacturer's protocol (Illumina NovaSeq 6000). Raw sequencing data files were converted to fastq files and aligned to the human reference genome hg38 using the Cell Ranger Pipeline 7.0.0 (10X Genomics). The data were clustered, and gene expression were analyzed using the Seurat R package. Pathway enrichments were performed by Gene Set Variation Analysis (GSVA) package (88).

**Human glioma cells stably transfected to express $IDH1^{R132H}$**

SJGBM2 pediatric glioma cells are ATRX mutant, and they were cultured in IMDM medium with L-glutamine (0.3 mg/mL) (Gibco, 12440061), 20% FBS (Peak Serum, PS-FB3), and antibiotic-antimycotic (1X) (Gibco, 15240-062) at 37°C, 5% $CO_2$. SJGBM2 cells were seeded in a 6-well plate (1.5 x $10^5$ each per well), and after 24h they were transfected with p-CMV-$IDH1^{R132H}$-Entry plasmid using jetPRIME transfection system (VWR, 89129-922). One day after transfection, the medium was replaced by selection medium containing G418 (Gibco, 10131027) at a concentration of 800 µg/mL. On day 15 of selection, individual cell colonies were taken from the well using autoclaved filter paper squares (2x2 mm approx.) previously embedded in HyQTase Cell Detachment Solution (TroyBiologicals, SV3003001) and placed within wells of 24-well plates with the appropriate cell culture medium. After 24h, the medium was replaced with selection medium. Each well corresponded to an isolated colony that was expanded for in vitro experiments. $IDH1^{R132H}$ protein expression was confirmed by WB assay (15). For reduced growth conditions, FBS was reduced to a final concentration of 5%.

**Metabolomics profiling**

Mutant IDH1 and WT NS (3 X $10^6$ cells) were seeded in a T75 flask in complete media as described previously. A parallel plate for protein estimation and sample normalization was also set up with the same number of cells. After overnight incubation, the culture media was aspirated off and replaced with fresh media. The cells were then cultured for a further 24h. For intracellular metabolites, the media was aspirated off and the samples washed once with 1 mL cold PBS before incubation with 1 mL ice cold 80% methanol on dry ice for 10min. Thereafter, cell lysates were collected from each well and transferred into separate 1.5 mL Eppendorf tubes. The samples were then centrifuged at 12,000xG, and the vol of supernatant to collect for each experimental condition was then determined based on the protein concentration of the parallel plate. The collected supernatants were dried using SpeedVac Concentrator, reconstituted with 50% v/v methanol in water, and analyzed by mass spectrometry and processed as previously described (89). Heatmaps were plotted with Heatmap2 package in R.

**Generation of human glioma NS from patient tumor biopsies**

SF10602 cells were generated in Dr. Joseph Costello's laboratory at UCSF from resected biopsies from patients harboring mIDH1 tumors with TP53 and ATRX mutation (45). SF10602 were cultured and maintained in serum free glioma neural stem (GNS) cell medium comprises of Neurocult NS-A (StemCell Technologies, 05750) supplemented with 2 mM L-glutamine (Gibco, 25030081), N-2 (Gibco, 17502-048), B-27 (without vitamin A, Gibco, 12587-010), Normocin (100 μg/ml) (InvivoGen, ant-nr-1), 0.1 mg/mL sodium pyruvate (Gibco, 11360-070) and Antibiotic-Antimycotic (1X) (Gibco, 15240-062) at 37°C, 5% $CO_2$. Human EGF (animal-free; PeproTech, AF-100-15), human FGF (animal-free; PeproTech, 100-18B) and PDGF-α (animal-free; PeproTech, 100-13A) were added twice weekly using 1μL of

20 ng/μL stock solution for each growth factor, equivalent to 1000X formulation per 1mL medium. For reduced growth conditions, human-EGF, human-FGF, and PDGF-α were added twice weekly at 1 μL (20 ng/μL) per 3 mL medium for a final concentration of 6.6 ng/mL.

**Western blot**

Mouse NS and human glioma cells (1.0 x $10^6$ cells) were harvested for each genotype, and total protein extracts were prepared in a RIPA lysis and extraction buffer (Thermo Fisher Scientific, Pierce, 89900) with 1X of Halt protease and phosphatase inhibitor cocktail (Thermo Fisher Scientific, 78442). 20 μg of protein extract (determined by bicinchoninic acid assay (BCA), Pierce, 23227) were separated by 4-12% SDS-PAGE (Thermo Fisher Scientific, NuPAGE, NP0322BOX) and transferred to nitrocellulose membranes (Bio-Rad, 1620112). The membrane was probed with 1:2000 of a rabbit anti-ATG9b Ab (Novus Biologicals, NBP1-77169), 1:1000 of a rabbit anti-ATG7 Ab (Cell Signaling Technology, 8558S), 1:1000 of a rabbit anti-pULK1 (S555) Ab (Cell Signaling Technology, 5869); 1:1000 of a rabbit anti-pULK1 (S757) Ab (Cell Signaling Technology, 6888); 1:1000 of a rabbit anti-ULK1 (Cell Signaling Technology, 8054); 1:2000 of a rabbit anti-MST4 Ab (Abcam, ab52491); 1:1000 of a rabbit anti-ATG4b Ab (Cell Signaling Technology, 5299); 1:1000 of a rabbit anti-phosphoATG4b Ab (Cell Signaling Technology, 19386);  1:1000 of an anti-UVRAG Ab (Abcepta, AP1850d); 1:1500 of a rabbit anti-LC3 I/II Ab (Novus Biologicals NB100-2220); 1:2000 of an anti-β-actin Ab (Cell Signaling Technology, 3700), then followed by secondary [Dako, Agilent Technologies, goat anti-rabbit 1:4000 (P0448), rabbit anti-mouse 1:4000 (P0260)]. Enhanced chemiluminescence reagents were used to detect the signals following the manufacturer's instructions (SuperSignal West Femto, Thermo Fisher Scientific, 34095). Blots were imaged using a ChemiDoc (Bio-Rad ChemiDoc™ MP System). WB quantification was performed using ImageJ, and the reported data are from three replicates. For the cells treated with α-ketoglutarate (α-KG), mIDH1 mouse NS (5.0 x $10^5$ cells) were treated with 2.5 mM α-KG (Cayman Chemicals, 876150-14-0) for 5 hours prior to protein extraction.

**Western Blot Assay of Histones**

To assess specific histone markers post translational modifications, histone extracts were obtained using Histone Purification Mini Kit (Active Motif, 40026) and WB were performed on 1.5 µg of histone extract. Histone markers' specific antibodies were as follows: 1:1000 of H3K4me3 antibody (Hologic Diagenode, C15410003); 1:1000 of H3K36me3 antibody (Hologic Diagenode, C15410192); 1:2000 of H3K27me3 antibody (Millipore, B07-449) for human glioma cell protein extracts; 1:1000 of H3K27me3 antibody (Hologic Diagenode C1541095); and 1:2000 of total histone H3 antibody (Cell Signaling Technologies, 4499).

**Metabolic Flux Assay**

To assess metabolic activity, oxygen consumption rates (OCR) and extracellular acidification rates (ECAR) were performed using the XF-96 Extracellular Flux Analyzer (Agilent). Cells were washed and resuspended in nonbuffered DMEM, adjusted to pH~7.4. WT-IDH1 and mIDH1 cells (1x$10^5$ cells/well) were seeded on laminin-coated plates and allowed to equilibrate for 30min in a non-$CO_2$, 37°C incubator. OCR and ECAR were measured under basal conditions and in response to the following mitochondrial inhibitors: oligomycin (1 µM), FCCP (1 µM), rotenone (100 nM), and antimycin A (1 µM). After the assay, OCR and ECAR measurements were normalized to cell number using CyQuant NF analysis (Invitrogen). For R-2HG treatment, cells were incubated in 2.5 mM of (2R)-Octyl-α-hydroxyglutarate for 5h before assessing metabolic activity.

**Mitochondria staining**

To assess the mitochondrial morphology, cells (1 x $10^5$ cells) were plated and were left overnight in 37°C incubator with 5% $CO_2$. Following incubation, cells were labeled with

MitoTracker Green FM (50 nM, Thermo Fisher Scientific, M7514) for 45 mins according to the manufacturer's instructions with slight modification. Cells were also stained with Hoechst 33342 dye (5 µM; Invitrogen Fisher Scientific, H3570) for 20 mins to label the nucleus. Cells were subsequently rinsed with HBSS and replaced by fresh growth medium before analyzed through confocal microscopy.

**Autophagy flux assay using mCherry-GFP-LC3 construct**

Lentiviral particles expressing mCherry-GFP-LC3 were generated by the University of Michigan Vector Core using the FUW mCherry-GFP-LC3 plasmid (56), which was gift from Anne Brunet. WT-IDH1 and mIDH1 SJGBM cells were cultured in a T-12.5 flask and infected with Lenti-mCherry-GFP-LC3 particles in a concentration of 1X of lentivirus. After 36h, $5x10^4$ cells were plated in a confocal microscopy chamber cover glass system, in presence or absence of 10 µM of chloroquine and cultured for 12h before analysis of GFP and mCherry expression by confocal microcopy.

**Confocal microscopy and image analysis**

Cells were placed into the incubator chamber of the microscope at 37°C with a 5% $CO_2$ atmosphere. Microscopy confocal images were acquired using a single photon laser scanning inverted confocal microscope LSM 880 AxioObserver (Carl Zeiss, Jena, Germany). Two laser lines were used for simultaneous excitation with a Plan-apochromat 63x/1.4 numerical aperture (NA) oil DIC M27 objective. Cells transfected with mCherry-GFP-LC3 were excited at 488 nm and 561 nm, and cells stained with MitoTracker™ green dye at 405 nm and 488 nm wavelength, respectively. Line-sequential scan mode and 1.4 scan zoom were used. The system was driven by ZEN Black software. Multichannel immunofluorescence images were processed and analyzed using Fiji (NIH). Autophagosome (green) and autolysosome (red) were manually identified and quantified using the plugging Cell Counter.

**In vitro experiments with radiation and autophagy inhibitors**

WT-IDH1 and mIDH1 mouse NS and human glioma cells (i.e., NPA/NPAI, SJ-GBM2/SJ-GBM2 mIDH1, and SF10602/SF10602 with AGI-5198) were plated at a density of $1x10^3$ cells/well in a 96-well plate 24h before treatment. WT-IDH1 and mIDH1 mouse NS (NPA/NPAI) were then incubated with either ATG7 siRNAs (100 nM; Origene, SR427399), ATG4b siRNAs (100 nM; Origene, SR413163), ULK101 (1 µM; SelleckChem, S8793), or NSC185058 (5 µM; Cayman Chemical, 23957) in combination with 3Gy of IR, 2h post-inhibitor treatment. Human glioma cells (SJGBM2/SJGBM2 mIDH1, and SF10602/SF10602 with AGI-5198) were then incubated with either ATG7 siRNAs (150 nM; Origene, SR323157), ATG4b siRNAs (150 nM; Origene, SR323518), ULK101 (3 µM; SelleckChem, S8793), or NSC185058 (15 µM; Cayman Chemical, 23957) in combination with 5Gy (SJGBM2/SJGBM2 mIDH1) or 20Gy (SF10602/SF10602 with AGI-5198) of IR, 2h post-inhibitor treatment. Cell viability was evaluated 72h post-IR treatment using CellTiter-Glo 2.0 (Promega, G9242) luminescence cell viability assay, following the manufacturer's protocol. The resulting luminescence was read with the Enspire Multimodal Plate Reader (PerkinElmer, 2300-0000). Data were represented graphically using GraphPad Prism software (version 8), and statistical significances were determined using one-way ANOVA followed by Tukey's test for multiple comparisons.

**In vitro Dose-Response and evaluation of radio-sensitivity using mitochondrial electron transport chain inhibitors**

To assess the susceptibility of both mutant and WT-IDH1 human (SJ-GBM2 WT-IDH1/ SJ-GBM2 mIDH1 and SF10602/SF10602 with AG-881) and mouse NS (NPA/NPAI, CPA/CPAI, and RPA/RPAI) to mitochondrial electron transport chain inhibitors (mETCi), two Complex I inhibitors were used: Rotenone (SelleckChem, S2348) and Metformin (SelleckChem, S1950) under normal and growth factor starved conditions. Both cell lines were plated at a density of 1000 cells per well in a 96-well plate (Fisher, 12-566-00) 24h prior to treatment where wells per inhibitor dose (1 μM, 2.5 μM, 5 μM, 10 μM, 20 μM, and 30 μM) were evaluated for each cell type. Cells were then incubated for 3 days under normal (EGF: 1:1000, FGF: 1:1000, and PDGFα: 1:1000 dilutions) as well as growth factor starved conditions (EGF: 1:3000, FGF: 1:3000, and PDGFα: 1:3000 dilutions) and viability was assessed using CellTiter Glo 2.0 assay (Promega, G9242) following manufacturer's protocol. To assess radio-sensitivity of each cell line, cells were incubated with either free-Rotenone or Metformin, or in combination with radiation at their respective IC50 doses for 72 hours in triplicate wells per condition. Cells were pre-treated with both the inhibitors 2 hours prior to IR with 3Gy for mouse, 5Gy for SJ-GBM2 and 20Gy for SF10602 of radiation, respectively. Resulting luminescence was read with the Enspire Multimodal Plate Reader (PerkinElmer, 2300-13 0000).

**Release of cytokines in response to autophagy inhibition and radiation**

To assess the release of cytokines (DAMPs and type-I IFNs) into the conditioned media, mIDH1 mouse NS and human glioma cells were seeded at a density of $1x10^6$ cells/well in 6-well plates and allowed to settle overnight before treatment. The following day, mIDH1 mouse NS were treated with either ATG7 (100nM; Origene, SR427399) or ATG4b (100nM; Origene, SR413163) for 2h prior to exposure to 3Gy of IR, and the human glioma cells were treated with either ATG7 (150nM; Origene, SR323157) or ATG4b (150nM; Origene, SR323518) for 2h prior to exposure to 5Gy of ionizing radiation. After 72h, the levels of various cytokines in the cultures' supernatants were measured using ELISA, either following the manufacturer's protocol (Novus Biologicals) or at the Cancer Center Immunology Core, University of Michigan.

**Implantable syngeneic murine glioma models**

Female C57BL/6 mice, aged 6-8wk old, were used for implantation models. Intracranial tumors were established by stereotactic injection of $5x10^4$ WT-IDH1 or mIDH1 mouse tumor NS into the right striatum using a 22-gauge Hamilton syringe (1 μL/min) with the following coordinates: +1.00 mm anterior, 1.8 mm lateral, and 3.5 mm deep. The presence of tumors was verified 5 days post-implantation (DPI) by bioluminescence imaging.

**Generation of autophagy deficient glioma cells using SB transposon system**

Our laboratory has modelled both high grade glioma and mIDH1 low grade glioma by integrating oncogenic plasmid DNA into the neural stem cells present within the developing brain of post-natal mice utilizing the sleeping beauty (SB) transposon system. The high-grade glioma cells endogenously express genetic lesion that drive oncogenesis through NRASG12V and simulate tumor suppressor loss by ATRX and TP53 short hairpin knockdown; thus named NPA (NRASG12V, TP53, and ATRX) hereafter. The mIDH1 glioma cells endogenous express $IDH1^{R132H}$ along with genetic lesions that drive oncogenesis through NRASG12V and simulate tumor suppressor loss by ATRX and TP53 short hairpin knockdown; thus named NPAI (NRASG12V, TP53, ATRX, and $IDH1^{R132H}$) hereafter. Tumor NS were derived from endogenous NPA tumors that were adapted to in vitro culture and could be reimplanted into immunocompetent C57BL6 (The Jackson Laboratory, C57BL/6J Strain# 000664) mice for preclinical experiments in this study.

Sleeping-beauty derived glioma cells NPA and NPAI were transfected using jetPEI (VWR, catalog 89129-960) with the plasmid pT2-shATG7-EBFP and sleeping beauty transposase plasmid (Supplemental Figure 30A-B). After transfection, cells were allowed to grow for 72h then they were subjected to FACS for isolation of BFP cells. These cells were sorted 3 times each before confirming ATG7 knockdown via Western Blot assay (Supplemental Figure 30C-D).

**Generation of autophagy deficient mouse model using SB transposon system**

Using the SB transposon system (78), we generated a glioma model harboring $IDH1^{R132H}$, shATRX, and shP53 as previously described (15), plus a shRNA for ATG7 (shATG7) to disrupt the autophagy pathway into the tumor. The shATG7 candidates were obtained from codex database (http://cancan.cshl.edu/cgi-bin/Codex/Codex.cgi) and cloned into the pT2 plasmid replacing shATRX sequences from our previous PT2-shATRX construction (79), generating the pT2-ATG7-BFP (Supplemental Figure 30). A WB assay was performed to select the best candidate to use for the generation of the animal model. The selected shATG7-A (HP_603911) (CTCGAGTGCTGTTGACAGTGAGCGACCAGAAGAAGTTGAACGAGTATAGTGAAGCCACAGATGTATACTCGTTCAACTTCTTCTGGGTGCCTACTGCCTCGGAGAATTC) was finally used in the combination cocktail of plasmids as described in Supplemental Figure 30A-B.

**Genetically engineered mutant IDH1 glioma model**

All animal studies were conducted according to guidelines approved by the IACUC at the University of Michigan. All animals were housed in an AAALAC accredited animal facility; and they were monitored daily. Studies did not discriminate sex, and both male and females were used. The strains of mice used in the study were C57BL/6 (The Jackson Laboratories, 000664).

The wild type (WT)-IDH1 and mIDH1 glioma models used in this study were generated previously for our group using SB Transposon System (15). The plasmid used to generate those models are: (i) SB transposase and LUC (pT2C-LucPGK-SB100X, henceforth referred to as SB/Luc), (ii) a constitutively active mutant of NRAS, NRAS-G12V (pT2CAG-NRASV12, henceforth referred to as NRAS), (iii) a short hairpin against p53 (pT2-shp53-GFP4, henceforth referred to as shp53), (iv) a short hairpin against ATRX (pT2-shATRX53-GFP4, henceforth referred to as shATRX), and (v) mutant IDH1$^{R132H}$ (pKT-IDH1$^{R132H}$-IRES-Katushka, henceforth referred to as mIDH1). Neonatal P01 C57BL/6 mice were used in all experiments. The genotype of SB- generated mice involved these combinations: (i) NRAS, shP53, and shATRX (WT-IDH1) and (ii) NRAS, shp53, shATRX, and IDH1$^{R132H}$ (mIDH1). Mice were injected according to a previously described protocol (78). Plasmids were mixed in mass ratios of 1:2:2:2 or 1:2:2:2:2 (20 μg plasmid in a total of 40 μL plasmid mixture) with in vivo-jetPEI (Polyplus Transfection, 201-50G) (2.8 μL per 40 μL plasmid mixture) and dextrose (5% total) and maintained at room temperature for at least 15 min prior to injection. The lateral ventricle (1.5 mm AP, 0.7 mm lateral, and 1.5 mm deep from lambda) of neonatal mice (P01) was injected with 0.75 μL plasmid mixture (0.5 μL/min) that included: (1) SB/Luc, (2) NRAS, (3) shp53, (4) with or without shATRX, and (5) with or without IDH1$^{R132H}$. To monitor plasmid uptake in neonatal pups, 30 μL of luciferin (30 mg/mL) was injected s.c. into each pup 24-48h after plasmid injection. In vivo bioluminescence was measured on an IVIS Spectrum (Perkin Elmer, 124262) imaging system. For the IVIS spectrum, the following settings were used: automatic exposure, large binning, and aperture f=1. For in vivo imaging of tumor formation and progression in adult mice, 100 μL of luciferin solution was injected i.p. and mice were then anesthetized with oxygen/isoflurane (1.5-2.5% isoflurane). To score luminescence, Living Image Software Version 4.3.1 (Caliper Life Sciences) was used. A region of interest (ROI) was defined as a circle over the head, and luminescence intensity was measured using the calibrated unit's photons/s/cm2/sr. Multiple images were taken over a 25min period following injection, and maximal intensity was reported.

For survival studies, animals were monitored daily for signs of morbidity, including ataxia, impaired mobility, hunched posture, seizures, and scruffy fur. Animals displaying symptoms of morbidity were intracardially perfused using Tyrode's solution, followed by fixation with 4% paraformaldehyde (PFA) in PBS.

**Generation of iRGD Synthetic Protein Nanoparticle (SPNP) with siRNA against *ATG7***

*SPNP formulation*:

The preparation of the SPNP formulations followed previously reported methods but with modifications (44). In the basic formulation, two solutions were prepared separately then combined. HSA (7.5% w/v) was solubilized in a solvent system comprised of ultrapure deionized water and ethylene glycol (80:20 v/v). The peptide, iRGD was then added along with BSA Alexa Fluor 647 conjugate (0.25% w/w relative to the albumin) to produce fluorescently labeled SPNPs. Then, the siRNA, which was resuspended according to the manufacturer's protocol, was complexed for 30min at room temperature under rotation with 60 kDa branched polyethyleneimine (5% w/v) before also being added to the serum albumin solution. Next, a bi-functional macromer, O,O'-Bis[2-N-Succinimidyl-succinylamino)ethyl]polyethylene glycol, was added at 10% w/w relative to the HSA solution. Once mixed, the two solutions were mixed to form the final formulation. For empty SPNP groups, all the components that were added in the iRGD ATG7i-SPNPs were included except for the ATG7 siRNA.

*SPNP fabrication*

SPNPs were fabricated via electrohydrodynamic jetting (44). The parameters used were the same as previously described. Briefly, the final formulation was loaded into 1 mL syringes equipped with a 1.5" 25-gauge stainless steel blunt needle. It was pumped at a rate of 0.2 mL/h to form droplets at the base of the needle. A voltage source was connected to the needle and grounded at the collection plate, located 6 inches from the base of the pump. The voltage was adjusted typically to a range between 8 kV and 15 kV to achieve a stable Taylor cone whereby rapid evaporation of the solvent occurred, creating solid nanoparticles on the collection plate. The collection plate was replaced with a clean plate every 30min until the solution within the syringe emptied. The collection plates were enclosed and incubated for 7 days at 37°C to form stable crosslinks.

*SPNP collection and processing*

After the 1wk period of incubation, the SPNPs on collection plates were removed. About 3-4mL of 0.01% Tween 20 in DPBS was added to each pan and physically agitated with plastic razor blades to release the SPNPs off the plate. The SPNP suspension was collected into a falcon tube. Each pan was agitated with fresh 0.01% Tween 20 in DPBS three times. The collected SPNPs was tip sonicated at an amplitude of 7 for 30s (1s on and 3s off) in an ice bath to break up aggregates, strained through a 40 µm filter into a new falcon tube then centrifuged at 3220 RCF for 5min. The supernatant was removed and distributed into 2 mL Eppendorf tubes and centrifuged for 1h at 4˚C at 21,500 RCF. The supernatant was discarded, and the resulting pellets were combined into a single 2 mL tube. The particles were washed two times with fresh DPBS without Tween 20.

*SPNP characterization*

Scanning electron Microscopy: Scanning electron microscopy samples were prepared by placing a silicon wafer on top of the collection plate during the jetting process. The samples were then placed on a copper tape covered scanning electron microscopy stub, then gold coated for 40s and visualized through the FEI NOVA 200 SEM/FIB instrument. The dry state SPNP quantification of morphology parameters were conducted through ImageJ analysis as described previously (90).

Dynamic Light Scattering: The hydrodynamic diameter and zeta potential was determined through dynamic light scattering on the Malvern Zetasizer. Samples were prepared by diluting the stock sample in DPBS and measured in folded capillary zeta cells. An average of at least three measurements was used to characterize each sample.

Bicinchoninic acid assay (BCA assay): BCA assay was used to quantify SPNP concentration. A standard curve was prepared for every SPNP concentration quantification measurement.

**In vitro mouse T cell proliferation assay**

To evaluate the effect of the combination of radiation (IR) treatment and autophagy inhibition, we tested the effect of IR (2 Gy) and the administration of iRGD nanoparticles loaded with a siRNA against *ATG7* (ATG7i-SPNP) in vivo. Twenty mice were implanted with mIDH1-OVA mouse tumor NS as stated above, and 7 days post implantation, mice were divided in the following 4 groups (n=5 mice): Control (no treatment); ATG7i-SPNP only; IR only; and IR + ATG7i-SPNP. Mice were treated with 2 Gy IR, for five consecutive days. ATG7i-SPNP ($2.0 \times 10^{11}$ particles) were administered 3 times total, once every other day starting on day 5. The mice were euthanized, and the spleens were collected on day 21 post implantation.

The spleens were collected to analyze the development of anti-tumor immune response. Splenocyte processing was performed as detailed previously (91). The splenocytes were cultured with 100 nM of SIINFEKL for 24 hours in 10% FBS-media with 55 µM 2-ME. Cells were then stained with anti-CD45, anti-CD3 and anti-CD8 antibodies, and proliferation was assessed by CFSE dye dilution. All flow data were acquired on a FACSAria flow cytometer (BD Biosciences) and analyzed using FlowJo version 10 (Treestar).

**IHC of paraffin-embedded brains**

Following perfusion, mouse brains were fixed in 4% paraformaldehyde (PFA) for an additional 48h at 4°C, then transferred to 70% ethanol, and processed and embedded in paraffin at the University of Michigan Microscopy & Image Analysis Core Facility using a Leica ASP 300 paraffin tissue processor/Tissue-Tek paraffin tissue embedding station (Leica). Tissue was sectioned using a rotary microtome (Leica) set to 5 µm in the z-direction. Antigen retrieval and IHC of paraffin-embedded sections were performed using antibodies and dilutions as follows: Tissue sections were blocked with blocking solution (1x PBS with 0.2% Tween-20 with 5% Goat Serum) for 2h. Sections were incubated with the following primary antibodies: anti-CD3ε (Cell Signaling, 99940); anti-CD68 (Abcam, ab125212); anti-MBP (Millipore, MAB386); anti-GFAP (Millipore, AB5541) at 4°C overnight. Tissue sections were then labeled with Secondary Biotinylated Ab (1:1000) in PBS with 0.2% Tween-20 for 15min at RT then 4°C overnight. ABC Avidin-Biotin-COMPLEX Binding was done using VECTASTATIN ABC

Reagent (Vectastain Elite ABC-HRP kit; Vector Laboratories, PK-6100). Sections were incubated for 1h with VECTASTAIN ABC Reagent in the dark and then washed with gentle agitation. Colorimetric Detection of Peroxidase was performed using Betazoid DAB Chromogen kit (BioCare BDB2004) according to the manufacturer's instructions. Afterward, Hematoxylin Counterstain was done, and coverslips were mounted with a xylene-based mounting medium and let dry on a flat surface at room temperature until imaging. Images were obtained using brightfield/epifluorescence (Zeiss Axioplan2, Carl Zeiss MicroImaging) or laser scanning confocal microscopy (Leica DMIRE2, Leica Microsystems) and analyzed using LSM5 software (Carl Zeiss MicroImaging). Immunostaining was performed using the Discovery XT processor (Ventana Medical Systems).

To access ATG7 expression levels post treatment of ATG7i-SPNPs, we implanted mice intracranially with $2x10^4$ NPAI cells on Day 0. At day 10, mice were randomly divided into 2 groups of 3 mice each. One group was treated with saline and the other group with ATG7i-SPNPs in combination with IR. Mice received a total of 10 Gy of radiation in 5 days. Mice were administered three doses of ATG7i-SPNPs ($2x10^{11}$) at day 10, 12, and day 14. At the end of day 14, mice were perfused, and brain and liver tissues were collected for further analysis. Brain sections from these mice were stained for MBP (myelin basic protein) (Sigma, MAB386), GFAP (glial fibrillary acidic protein) (Sigma, AB5541), ATG7 (Invitrogen, PIPA535203), and cleaved Caspase-3 (Cell Signaling, 9661S).

**Supplemental Figure 1**

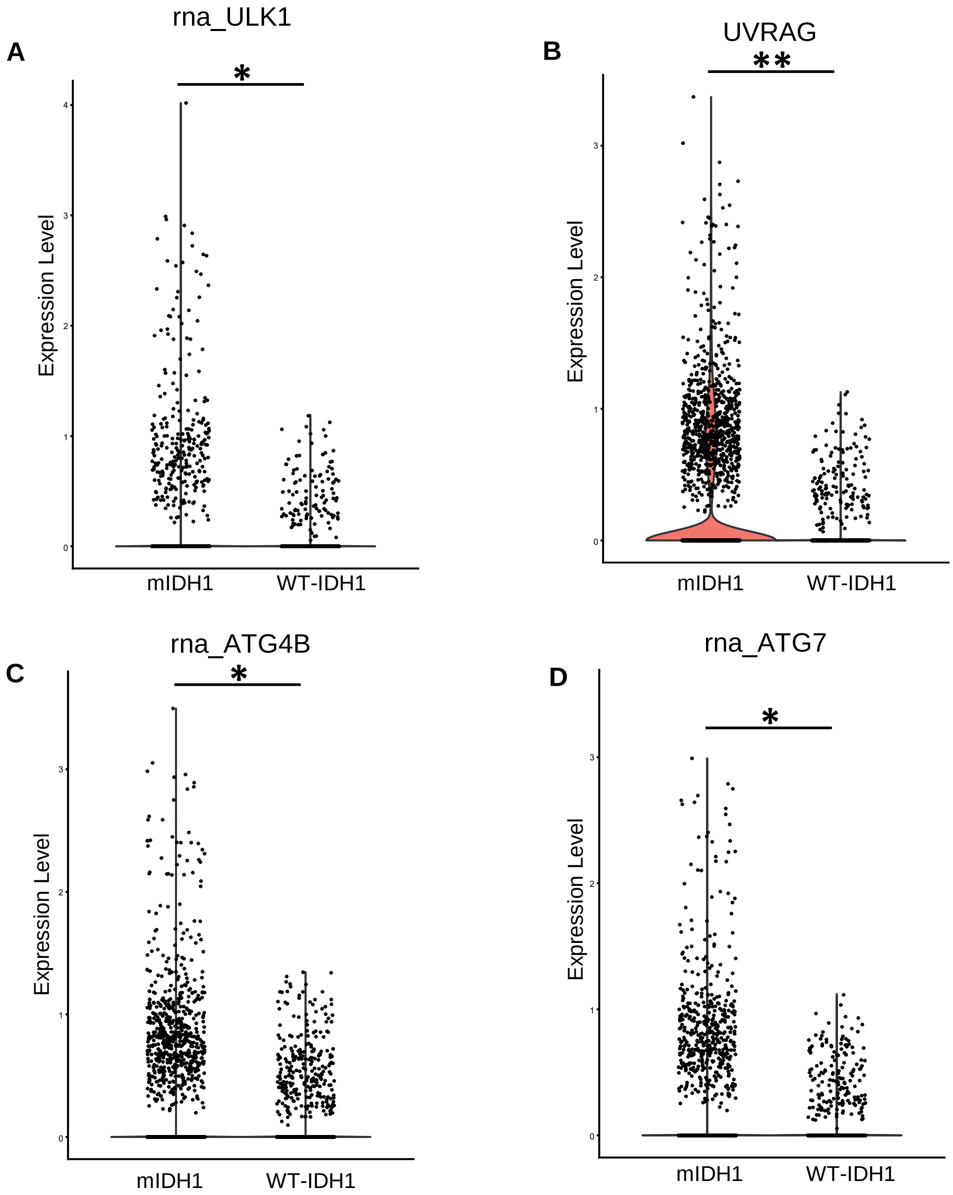


**Supplemental Figure 1:** Autophagy related gene expression of human scRNA-seq from glioma patients. Human glioma scRNA seq was analyzed for the expression of ULK1, UVRAG, ATG4B and ATG7 in WT-IDH1 and mIDH1 glioma tumor cells. Violin plots represent the expression of (**A**) ULK1 (**B**) UVRAG (**C**) ATG4B (**D**) ATG7. **P* < 0.01; ***P* < 0.005; unpaired t-test.

Supplemental Figure 2

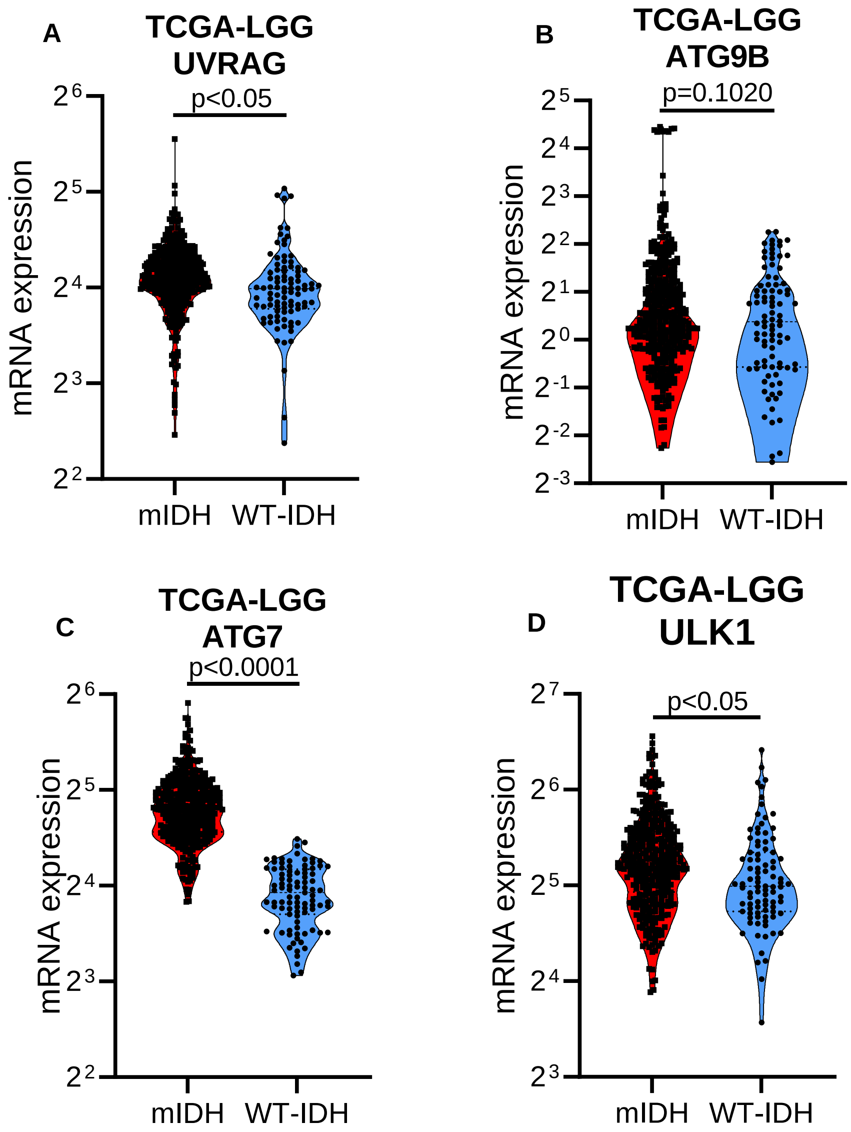


**Supplemental Figure 2:** Autophagy related gene expression from TCGA. (**A**) Analysis of RNA-seq data obtained from TCGA to quantify UVRAG mRNA in LGG with wtIDH1 or mIDH1. (**B**) Analysis of RNA-seq data obtained from TCGA to quantify ATG9B mRNA in LGG with wtIDH1 or mIDH1. (**C**) Analysis of RNA-seq data obtained from TCGA to quantify ATG7 mRNA in LGG with wtIDH1 or mIDH1. (**D**) Analysis of RNA-seq data obtained from TCGA to quantify ULK1 mRNA in LGG with wtIDH1 or mIDH1. Unpaired t test.

Supplemental Figure 3

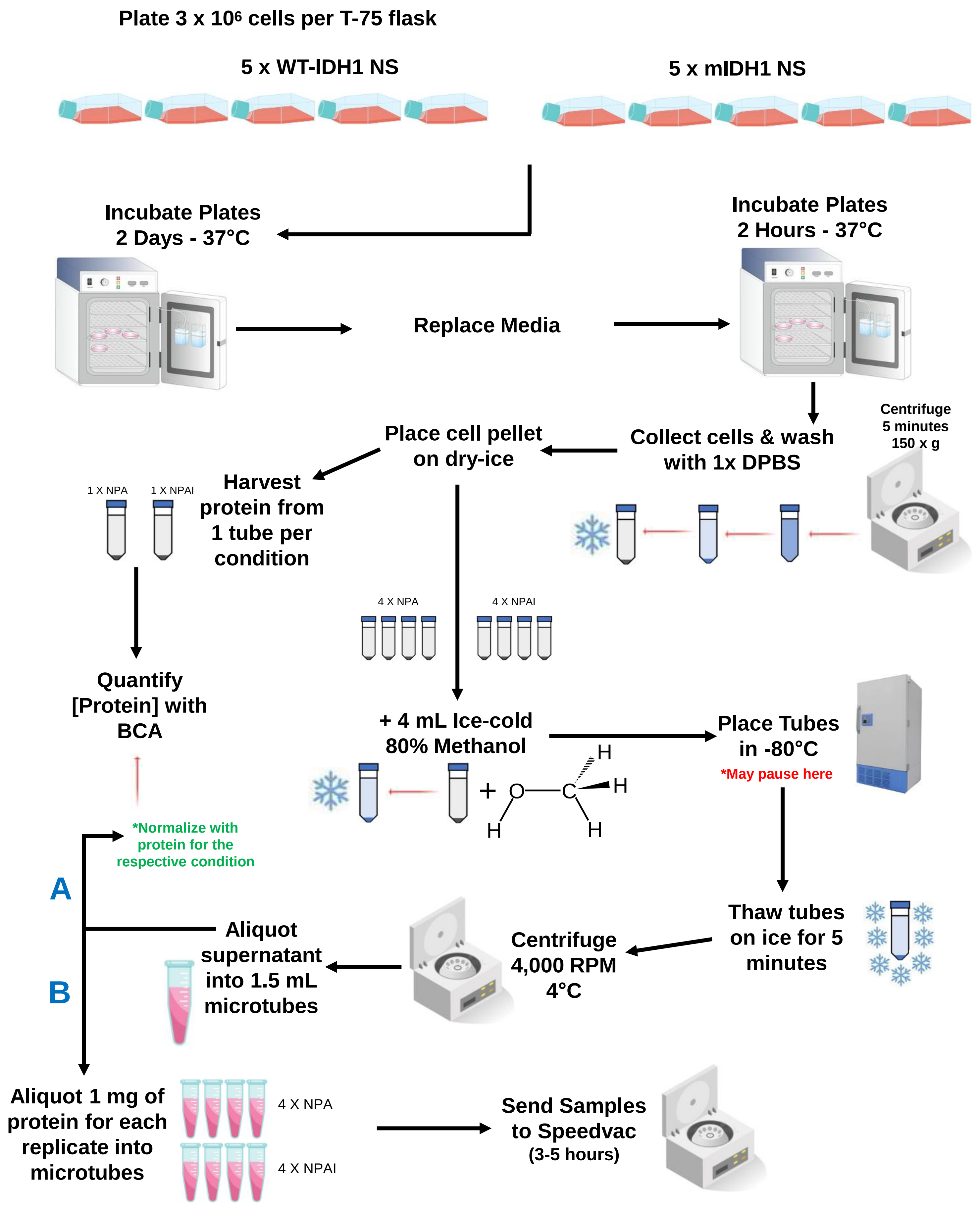


**Supplemental Figure 3:** Sample processing for metabolomic profiling in glioma cells. Diagram showing the method and sample processing for metabolic profiling in glioma cells.

**Supplemental Figure 4**

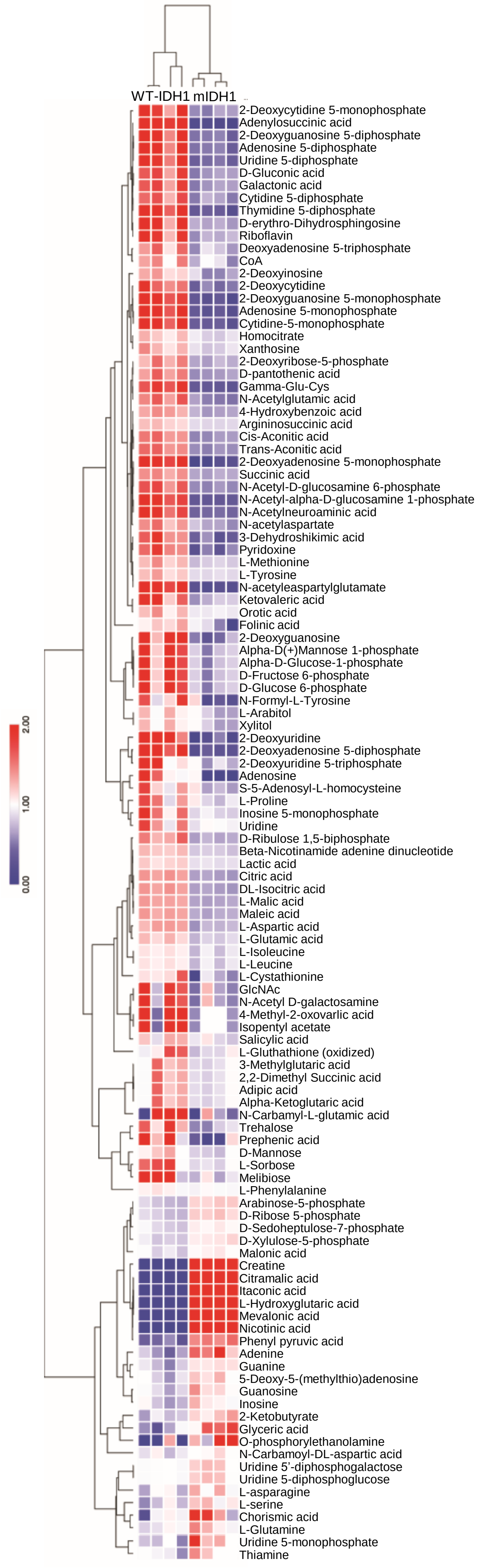


Filtered metabolites P < 0.01

**Supplemental Figure 4:** Heatmap of the metabolites in glioma cells. Heatmap showing the metabolomic profiling in WT-IDH1 and mIDH1 glioma NS filtered by $P < 0.1$.

**Supplemental Figure 5**

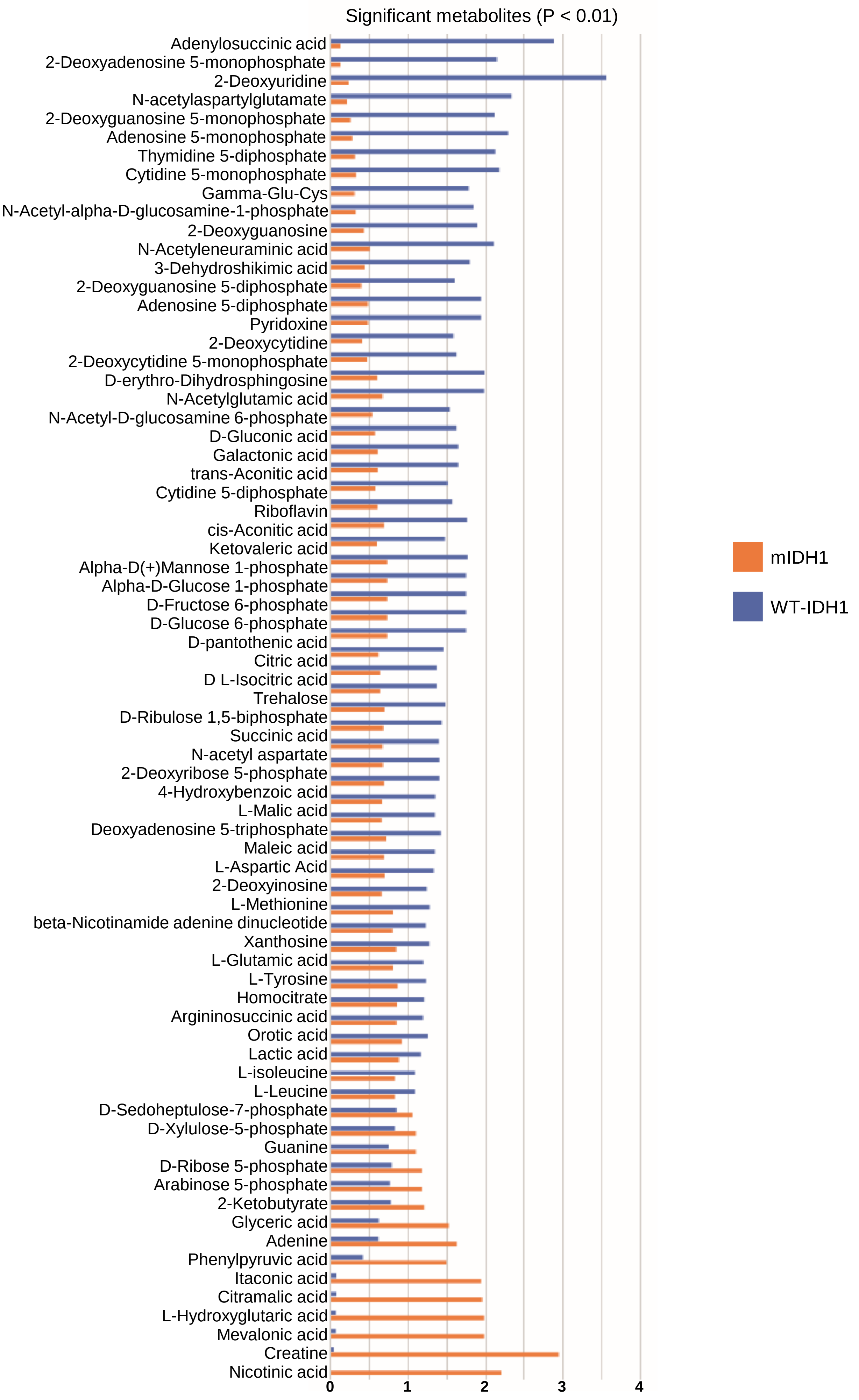

**Supplemental Figure 5:** Significantly different metabolites in glioma cells. Bar graph showing significantly different metabolites ($P < 0.01$) comparing the WT-IDH1 NS (blue bars) with the mIDH1 NS (orange bars) expressed in AU.

**Supplemental Figure 6**

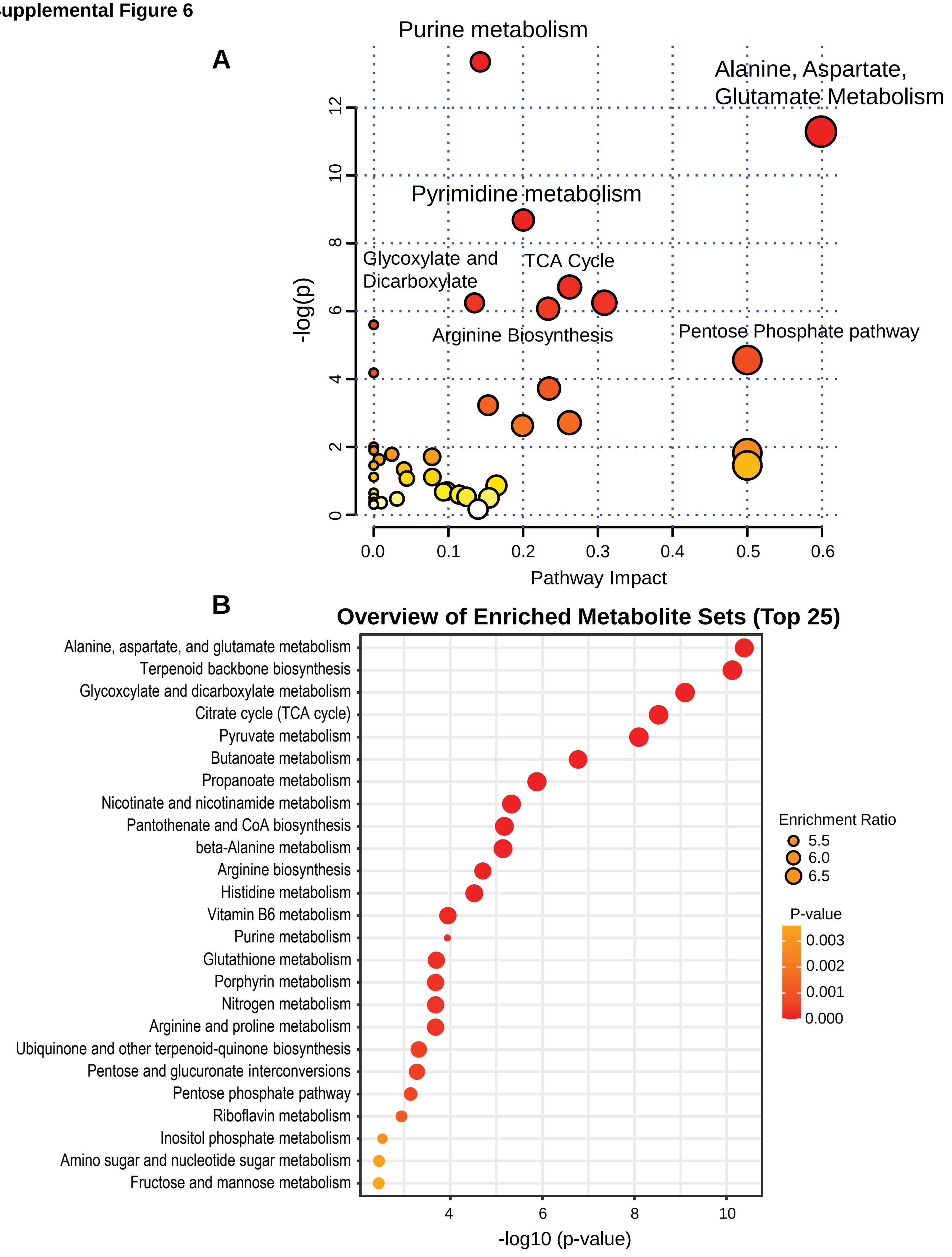


**Supplemental Figure 6:** Metabolic pathway analysis plot created using MetaboAnalyst. (**A**) Plots depict several metabolic pathway alterations induced in mIDH1 NS when compared with WT-IDH1 NS. The x-axis represents the pathway impact value computed from pathway topological analysis, and the y-axis is the negative log of the p-value obtained from pathway enrichment analysis. The pathways that were most significantly changed are characterized by both a high -log(p) value and high impact value (top right region). (**B**) Enrichment analysis on metabolic set alterations induced in mIDH1 NS when compared with WT-IDH1 NS.

**Supplemental Figure 7**

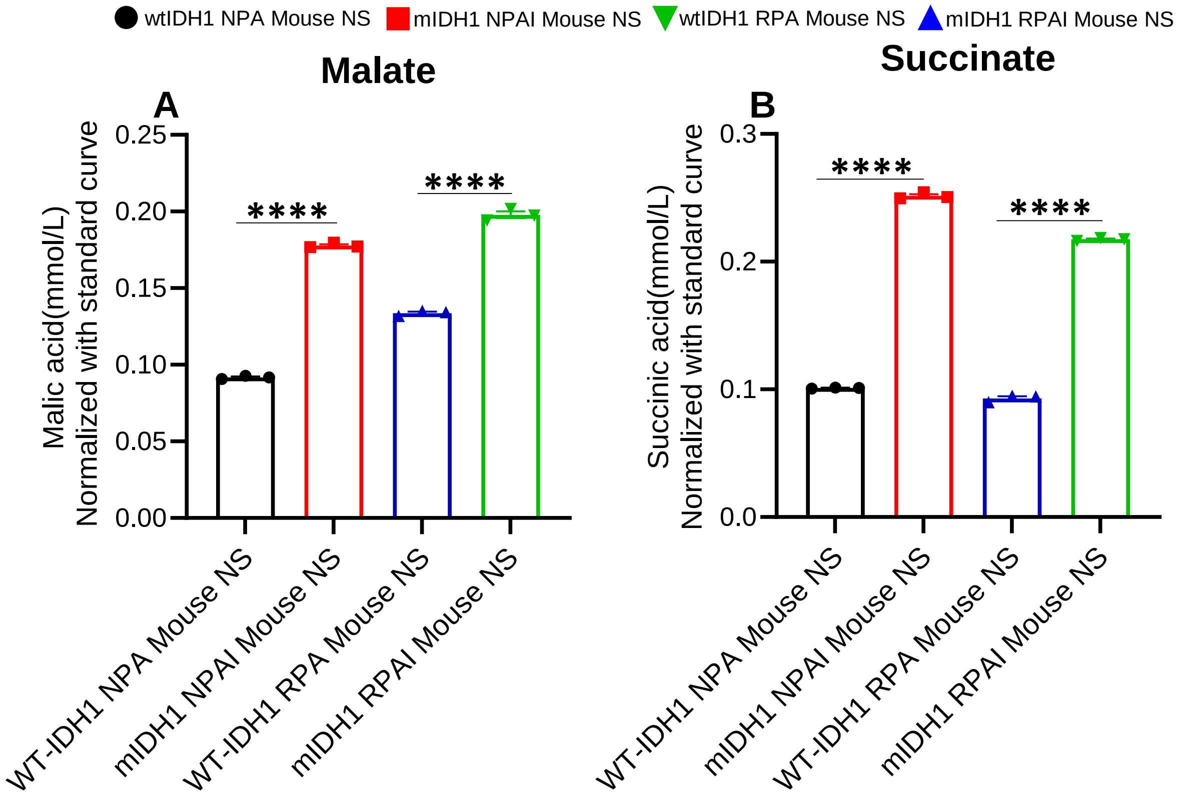


**Supplemental Figure 7:** Measurement of Malate and Succinate in genetically engineered mouse cells under reduced growth conditions. (**A**) Malate and (**B**) Succinate were measured via colorimetric assay in two different genetically engineered mouse models under reduced growth conditions. Black bars represent WT-IDH1 mouse NS. Red bars represent mIDH1 mouse NS. Blue bars represent PDGFRα-D842V driven WT-IDH1 mouse NS. Green bars represent PDGFRα-D842V driven mIDH1 mouse NS. Error bars represent SEM from independent biological replicates (n = 3). ****$P$ < 0.0001; unpaired t test.

**Supplemental Figure 8**

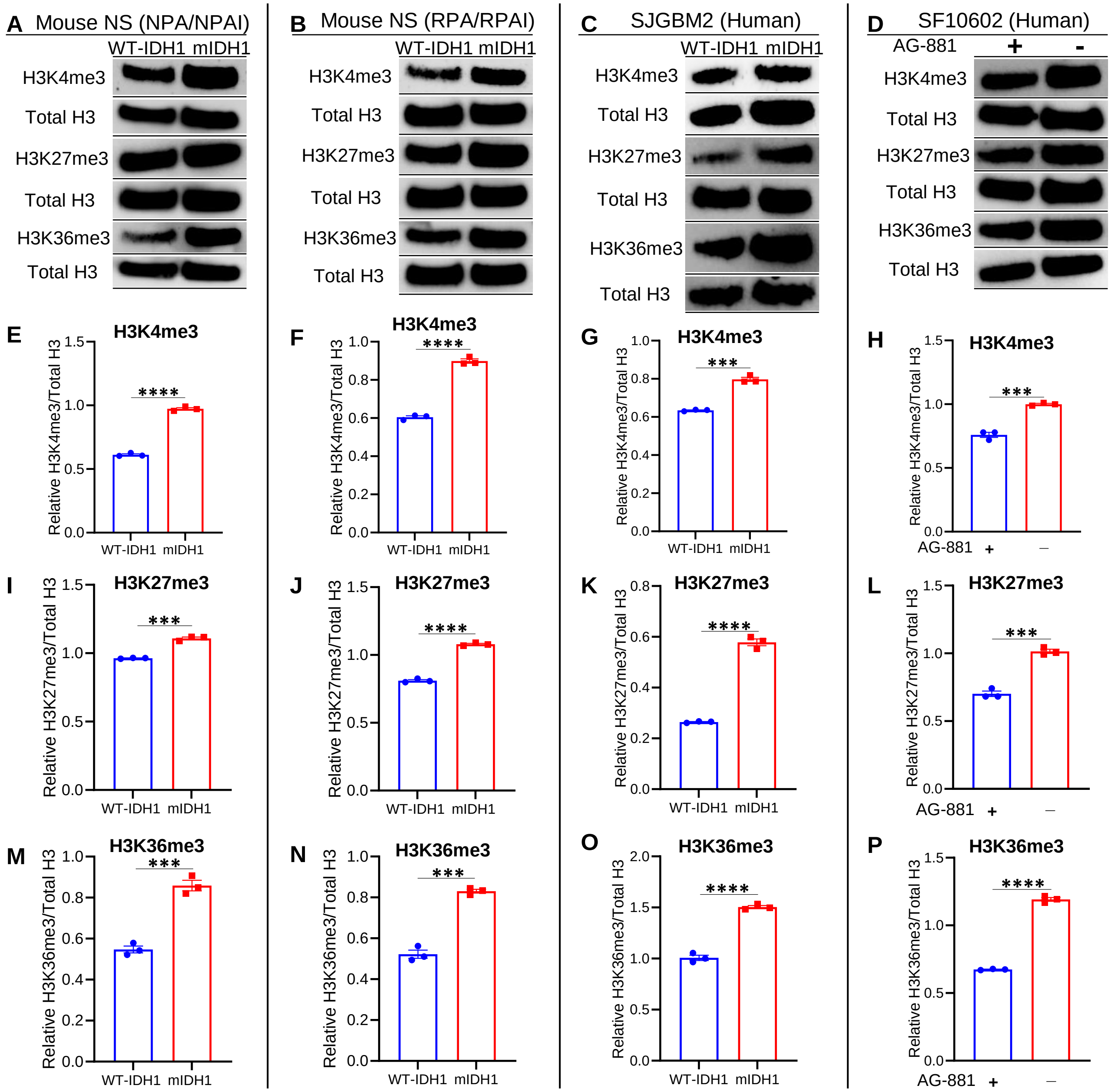


**Supplemental Figure 8:** Western Blot analysis of H3K4me3, H3K27me3, and H3K36me3 in mouse and human cells. (**A-D**) Western blot analysis of H3K4me3, H3K27me3, and H3K36me3 in (A) Mouse NS NPA and NPAI, (B) Mouse NS RPA and RPAI, (C) patient-derived SJGBM2 cells, and (D) patient derived SF10602 cells with and without treatment with mIDH1 inhibitor, AG-881. (**E-H**) ImageJ densitometric quantification of the western blots for H3K4me3 in (E) Mouse NS NPA and NPAI, (F) Mouse NS RPA and RPAI, (G) SJGBM2 cells, and (H) SF10602 cells with and without treatment with AG-881. (I-L) ImageJ densitometric quantification of the western blots for H3K27me3 in (I) Mouse NS NPA and NPAI, (J) Mouse NS RPA and RPAI, (K) SJGBM2, and (L) SF10602 with and without treatment with AG-881. (M-P) ImageJ densitometric quantification of the western blots for H3K36me3 in (M) Mouse NS NPA and NPAI, (N) Mouse NS RPA and RPAI, (O) SJGBM2, and (P) SF10602 with and without treatment with AG-881. Error bars represent SEM from independent technical replicates (n = 3). ***$P < 0.001$; ****$P < 0.0001$; unpaired t test.

**Supplemental Figure 9**

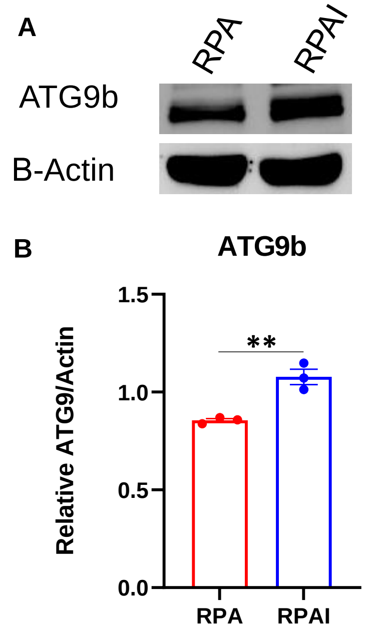


**Supplemental Figure 9:** Western Blot analysis of ATG9b in additional genetically engineered mouse model. (**A**) Western blot analysis of autophagy related gene ATG9b in WT-IDH1 cells (RPA) and mIDH1 cells (RPAI) with β-Actin as a loading control. (**B**) ImageJ densitometric quantification of the western blot for ATG9b and β-Actin. Error bars represent SEM from independent technical replicates (n = 3). ***P* < 0.005; unpaired t test.

**Supplemental Figure 10**

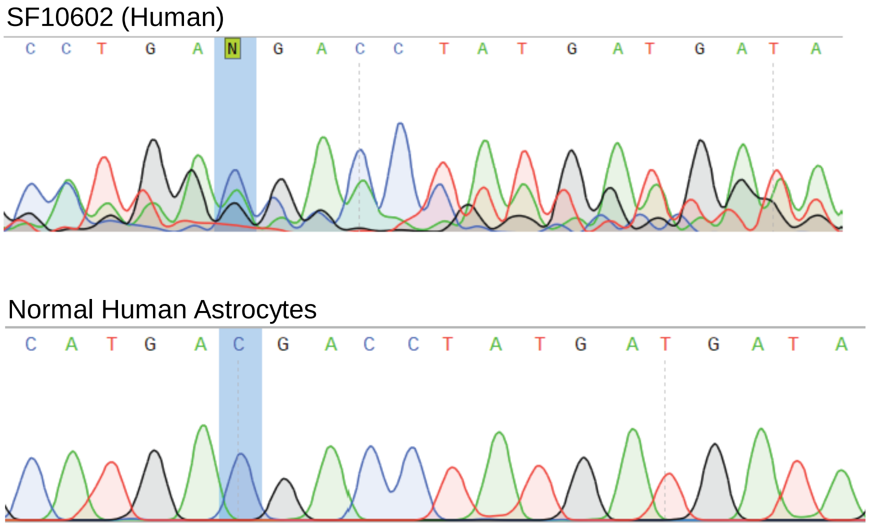


**Supplemental Figure 10:** Primary mIDH1 human glioma cells, SF10602 has both WT-IDH1 and mIDH1 mutations. Specific region of IDH1 mutations were amplified by PCR and sequenced by sanger sequencing.

Supplemental Figure 11

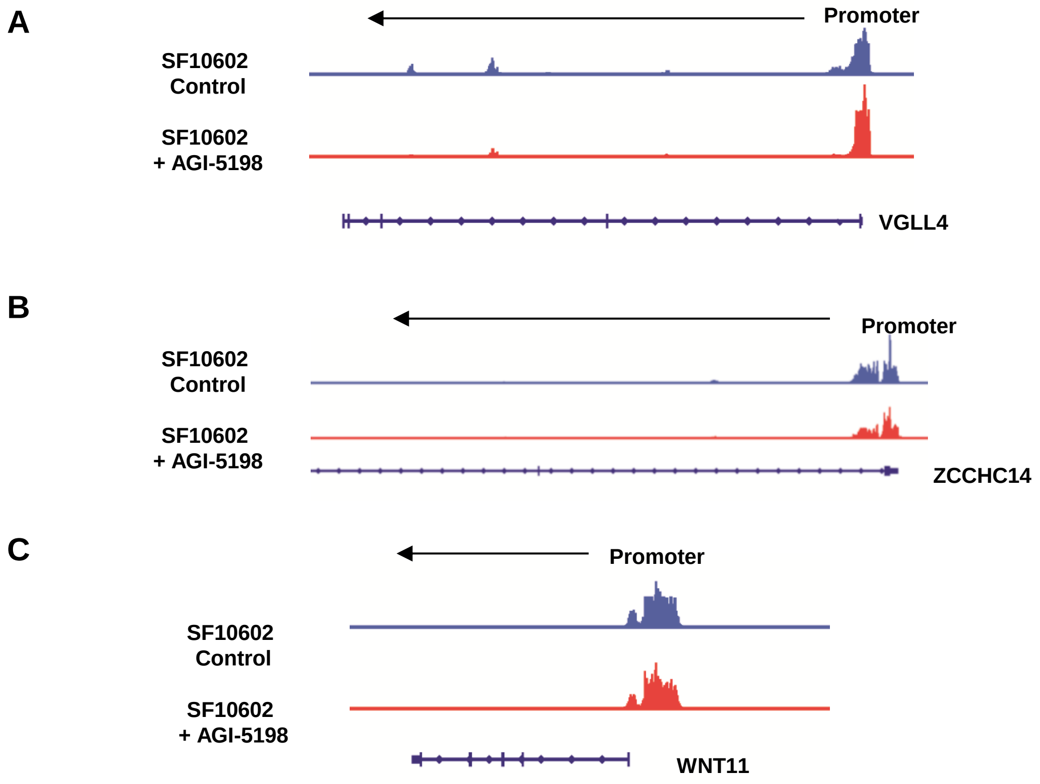


**Supplemental Figure 11:** H3K4me3 occupancy at promoter regions of genes that do not exhibit differential deposition in mIDH1 SF10602 cells vs. SF10602 cells treated with mIDH1 inhibitor AGI-5198 which are located in the vicinity of autophagy related genes, (**A**) *VGLL4* next to *ATG7*; (**B**) *ZCCHC14* next to *UVRAG*; (**C**) *WNT11* next to *MAP1LC3B* (LC3).

**Supplemental Figure 12**

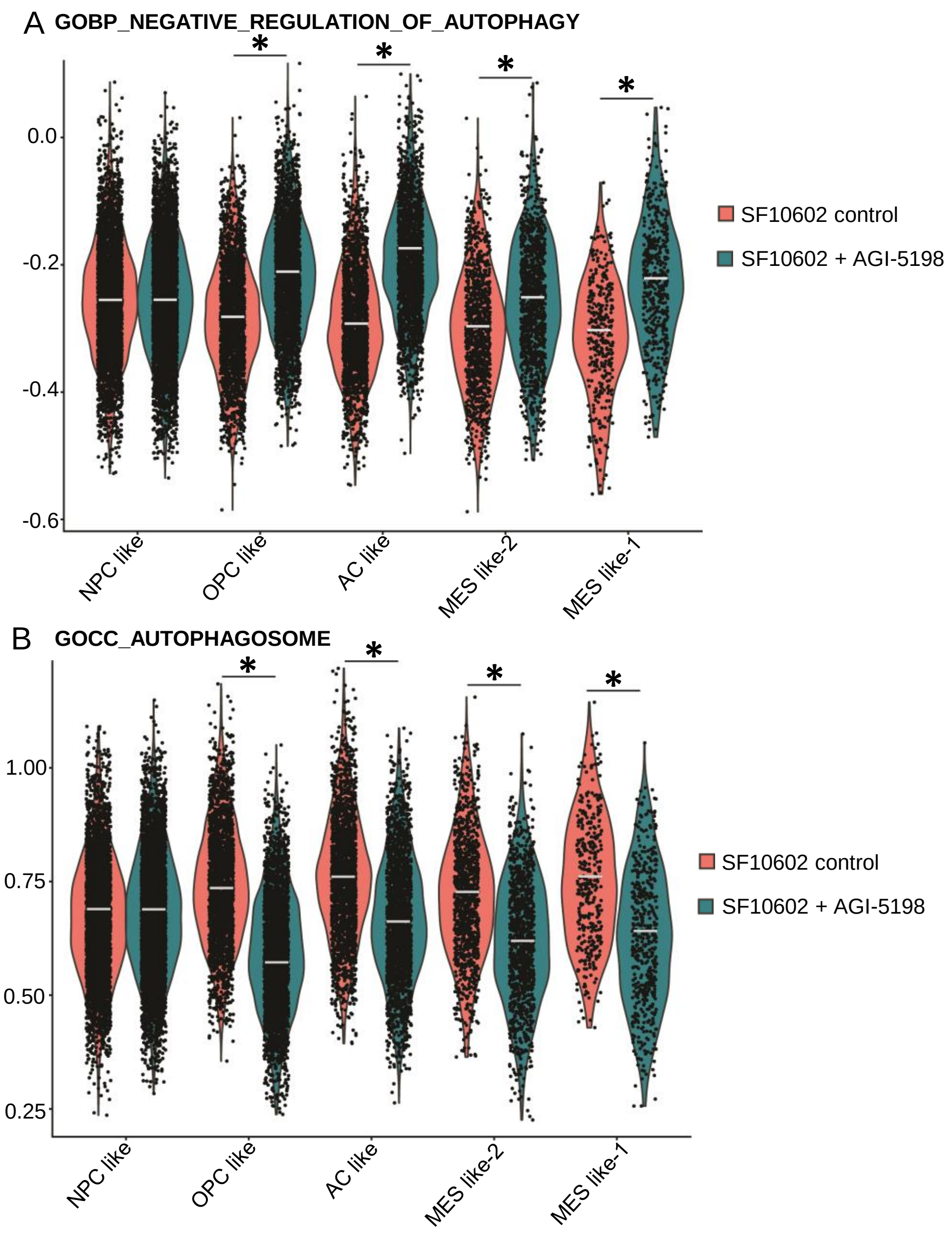


**Supplemental Figure 12:** Violin plots of scRNA-seq in human mIDH1 glioma 1 cells related to Fig. 4 (**A**) GO term enrichment score for GO biological process of negative regulation of autophagy. (**B**) GO term enrichment score for GO cellular component of autophagosome. Significance was measured via the Wilcoxon Rank Sum Test from the Seurat/Presto package. *$P < 0.05$.

**Supplemental Figure 13**

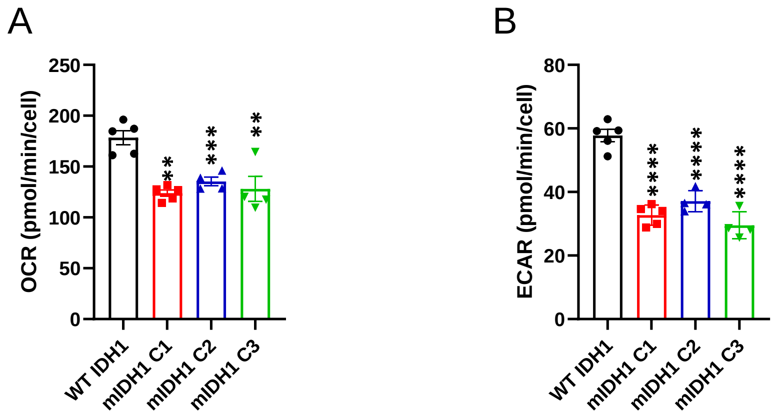


**Supplemental Figure 13:** Maximal OCAR and ECAR in glioma NS. Mitochondrial respiration of NS derived from WT or mIDH1 tumors (Clones 1-3) examined through the Mito Stress Test. (**A**) Basal oxygen consumption rates (OCAR) and (**B**) extracellular acidification rates (ECAR) of mIDH1 and WT glioma NS. Error bars depict SD of technical replicates from a representative plot of 3 independent experiments. $^{**}P < 0.01$, $^{***}P < 0.001$, $^{****}P < 0.0001$; one-way ANOVA.

**Supplemental Figure 14**

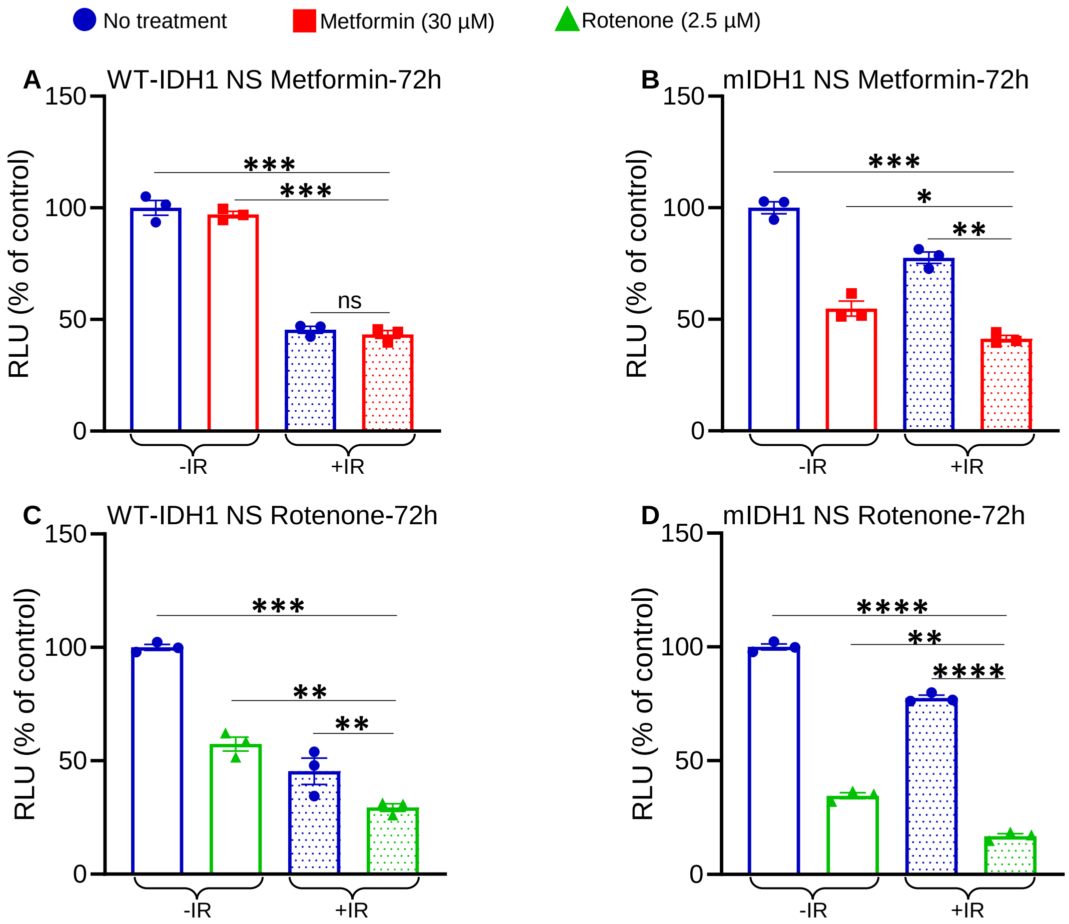


**Supplemental Figure 14:** Impact of mitochondrial complex 1 inhibition on glioma cell viability. In vitro inhibition of Complex I of ETC restores radiosensitivity in mIDH1 NS. (**A-B**) Impact of Metformin on radiosensitivity in NRAS/shP53/shATRX/mIDH1 (NPAI) +/- mouse NS cells. Cell viability assay shows the effect of Metformin at 30 µM doses in combination with radiation (3 Gy) on cell proliferation in (**A**) WT-IDH1 and (**B**) mIDH1 mouse NS. Results are expressed in relative luminescence units (RLU). *$P < 0.05$, **$P < 0.01$, ***$P < 0.001$; two-way ANOVA followed by Tukey's test (n= 3 technical replicates). (**C-D**) Impact of Rotenone on radiosensitivity in NPAI +/- mouse NS cells. Cell viability assay shows the effect of Rotenone at 2.5 µM doses in combination with radiation (3 Gy) on cell proliferation in (**C**) WT-IDH1 and (**D**) mIDH1 mouse NS. Results are expressed in RLU. *$P < 0.05$, **$P < 0.01$, ***$P < 0.001$; ****$P < 0.0001$; two-way ANOVA followed by Tukey's test (n = 3 technical replicates).

**Supplemental Figure 15**

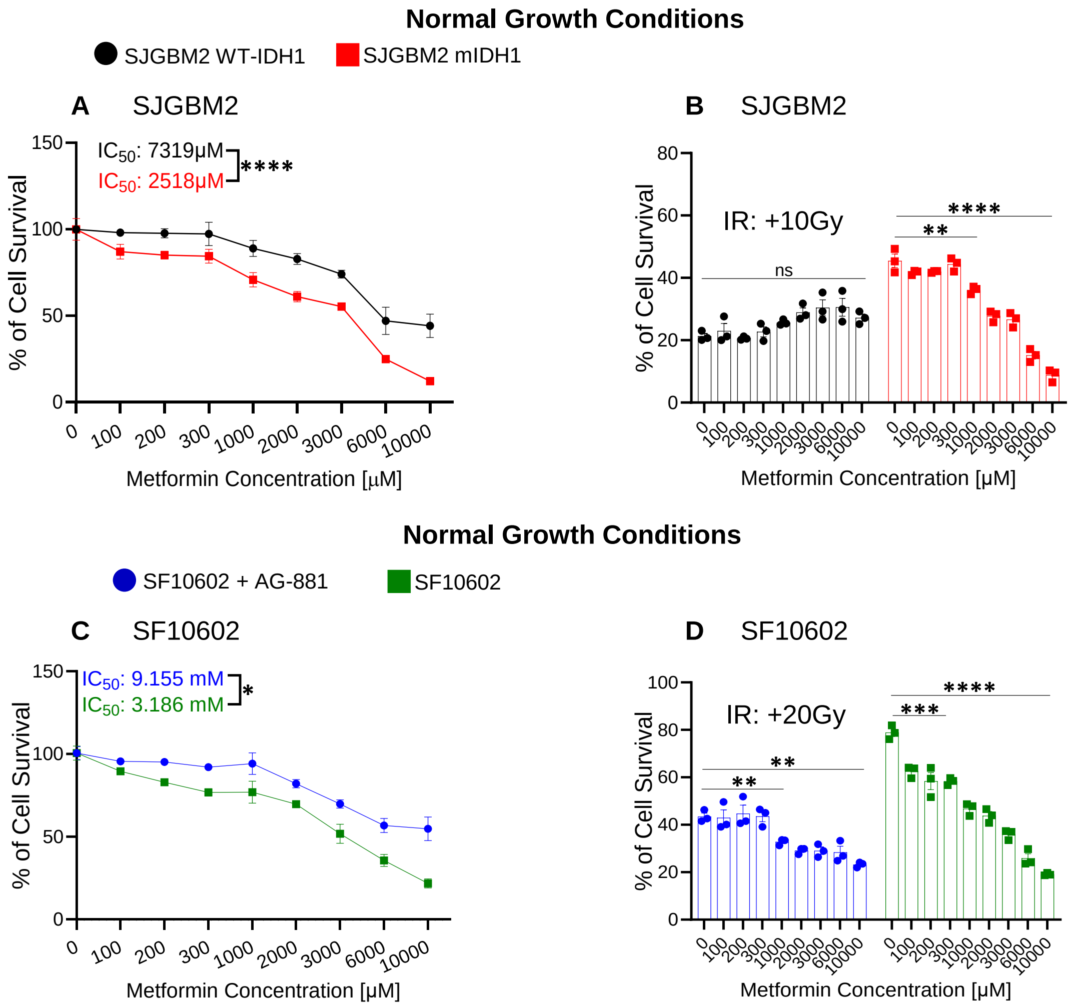


**Supplemental Figure 15:** In Vitro inhibition of Complex I of the ETC induces cell death and restores radiosensitivity under normal growth conditions in two human mIDH1 glioma cells. (**A**) Cell viability assay of WT-IDH1 and mIDH1 genetically engineered human glioma cells, SJGBM2, treated with metformin for 72 hours at increasing concentrations. Cell viability expressed as % of cell survival relative to control populations. A linear regression model and generalized additive models were used to determine the dose-response relationship. (**B**) Cell viability assay showing the effect of metformin at increasing concentrations in combination with radiation (10Gy) on the proliferation of WT-IDH1 and mIDH1 SJGBM2 cells. (**C**) Cell viability assay of endogenous mIDH1 glioma cells, SF10602, treated with and without mIDH1 inhibitor, AG-881, were treated with metformin for 72 hours at increasing concentrations. Cell viability expressed as % of cell survival relative to control populations. A linear regression model and generalized additive models were used to determine the dose-response relationship. (**D**) Cell viability assay showing the effect of metformin at increasing concentrations in combination with radiation (20Gy) on the proliferation of AG-881 treated SF10602 and untreated SF10602 cells. Results are expressed as % of Cell survival relative to control populations. Statistical analyses were conducted using student's t-test and error bars depict SD of technical replicates from 3 independent experiments. *$P$ < 0.1, **$P$ < 0.01, ***$P$ < 0.001, ****$P$ < 0.0001, ns = not significant.

**Supplemental Figure 16**

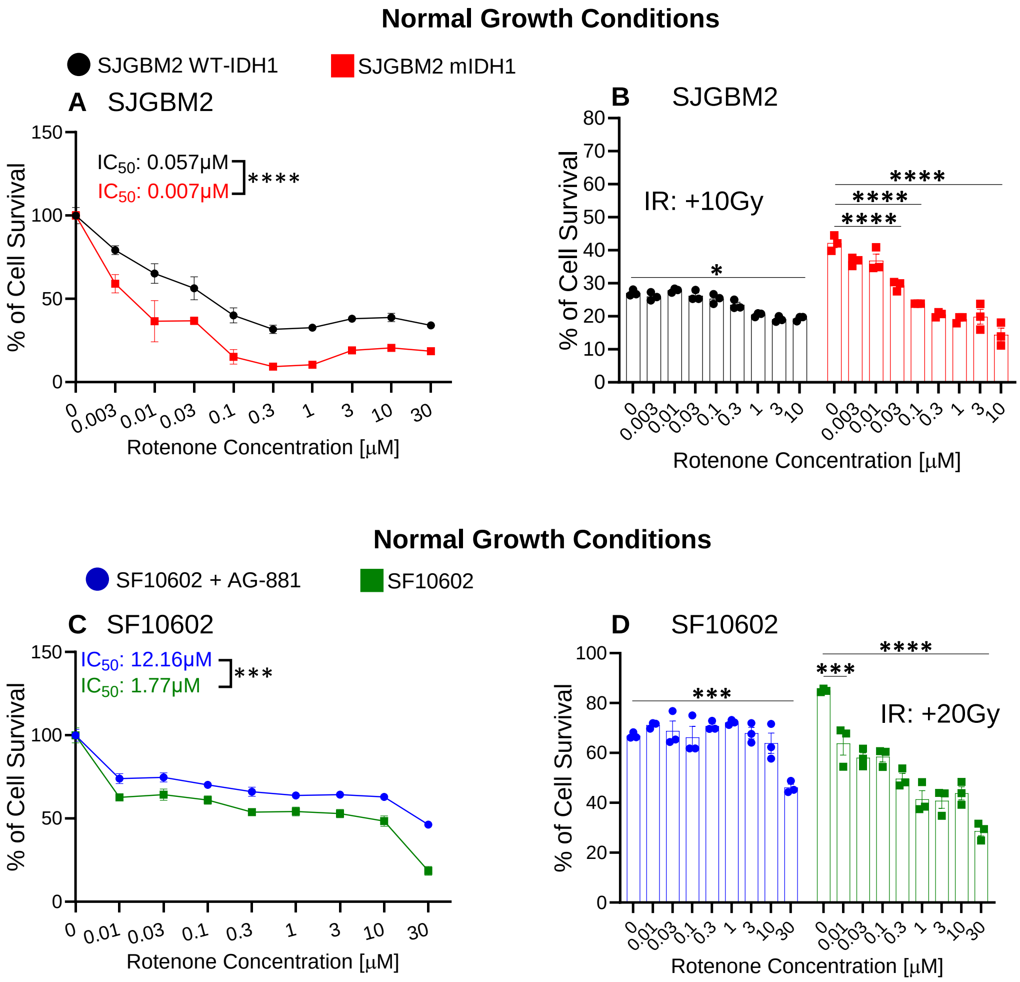


**Supplemental Figure 16:** In Vitro inhibition of Complex I of the ETC induces cell death and restores radiosensitivity under normal growth conditions in two human mIDH1 glioma cells. (**A**) Cell viability assay of WT-IDH1 and mIDH1 genetically engineered human glioma cells, SJGBM2, treated with rotenone for 72 hours at increasing concentrations. Cell viability expressed as % of cell survival relative to control populations. A linear regression model and generalized additive models were used to determine the dose-response relationship. (**B**) Cell viability assay showing the effect of rotenone at increasing concentrations in combination with radiation (10Gy) on the proliferation of WT-IDH1 and mIDH1 SJGBM2 cells. (**C**) Cell viability assay of endogenous mIDH1 glioma cells, SF10602, treated with and without mIDH1 inhibitor, AG-881, were treated with rotenone for 72 hours at increasing concentrations. Cell viability expressed as % of cell survival relative to control populations. A linear regression model and generalized additive models were used to determine the dose-response relationship. (**D**) Cell viability assay showing the effect of rotenone at increasing concentrations in combination with radiation (20Gy) on the proliferation of AG-881 treated SF10602 and untreated SF10602 cells. Results are expressed as % of Cell survival relative to control populations. Statistical analyses were conducted using student's t-test and error bars depict SD of technical replicates from 3 independent experiments. **P* < 0.1, ***P* < 0.01, ****P* < 0.001, *****P* < 0.0001, ns = not significant.

**Supplemental Figure 17**

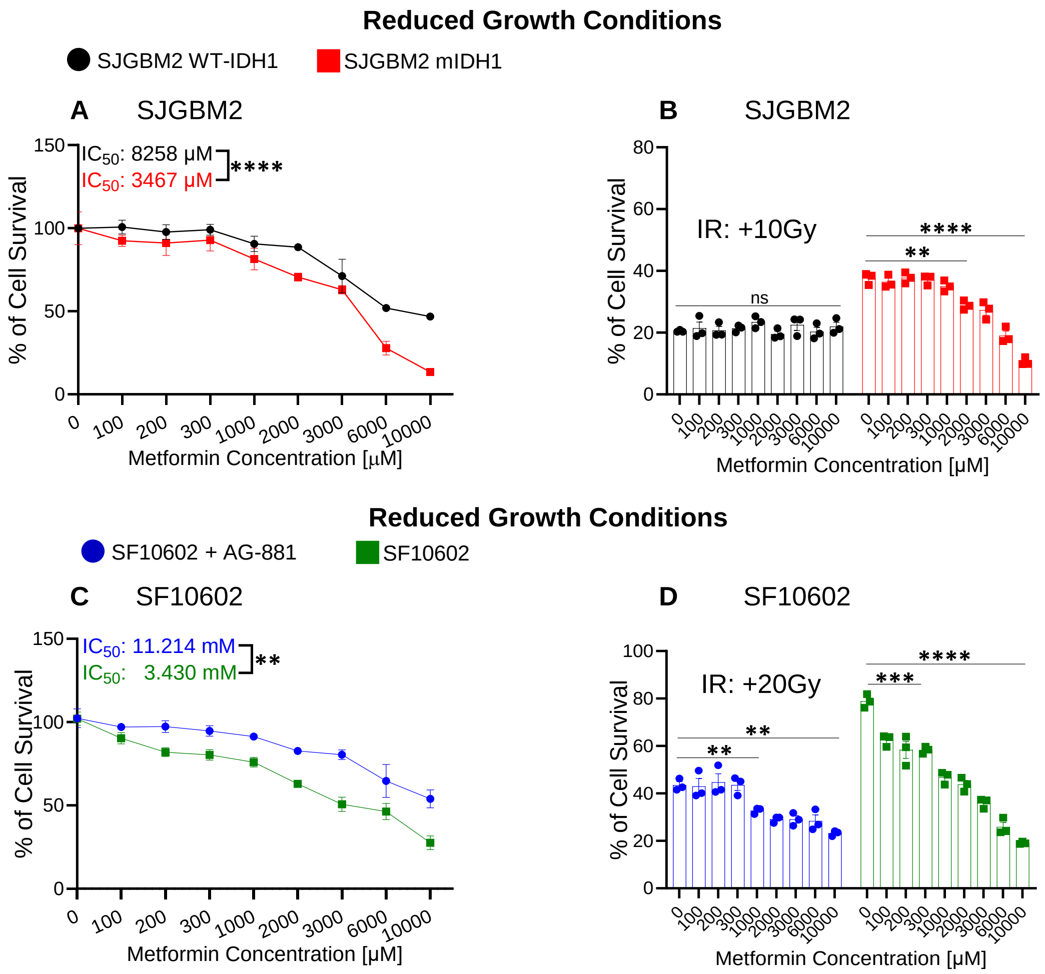


**Supplemental Figure 17:** In Vitro inhibition of Complex I of the ETC induces cell death and restores radiosensitivity under reduced growth conditions in two human mIDH1 glioma cells. (**A**) Cell viability assay of WT-IDH1 and mIDH1 genetically engineered human glioma cells, SJGBM2, under reduced growth conditions, treated with metformin for 72 hours at increasing concentrations. Cell viability expressed as % of cell survival relative to control populations. A linear regression model and generalized additive models were used to determine the dose-response relationship. (**B**) Cell viability assay showing the effect of metformin at increasing concentrations in combination with radiation (10Gy) on the proliferation of WT-IDH1 and mIDH1 SJGBM2 cells under reduced growth conditions. (**C**) Cell viability assay of endogenous mIDH1 glioma cells, SF10602, treated with and without mIDH1 inhibitor, AG-881, under reduced growth conditions, were treated with metformin for 72 hours at increasing concentrations. Cell viability expressed as % of cell survival relative to control populations. A linear regression model and generalized additive models were used to determine the dose-response relationship. (**D**) Cell viability assay showing the effect of metformin at increasing concentrations in combination with radiation (20Gy) on the proliferation of AG-881 treated SF10602 and untreated SF10602 cells under reduced growth conditions. Results are expressed as % of Cell survival relative to control populations. Statistical analyses were conducted using student's t-test and error bars depict SD of technical replicates from 3 independent experiments. *$P$ < 0.1, **$P$ < 0.01, ***$P$ < 0.001, ****$P$ < 0.0001, ns = not significant.

**Supplemental Figure 18**

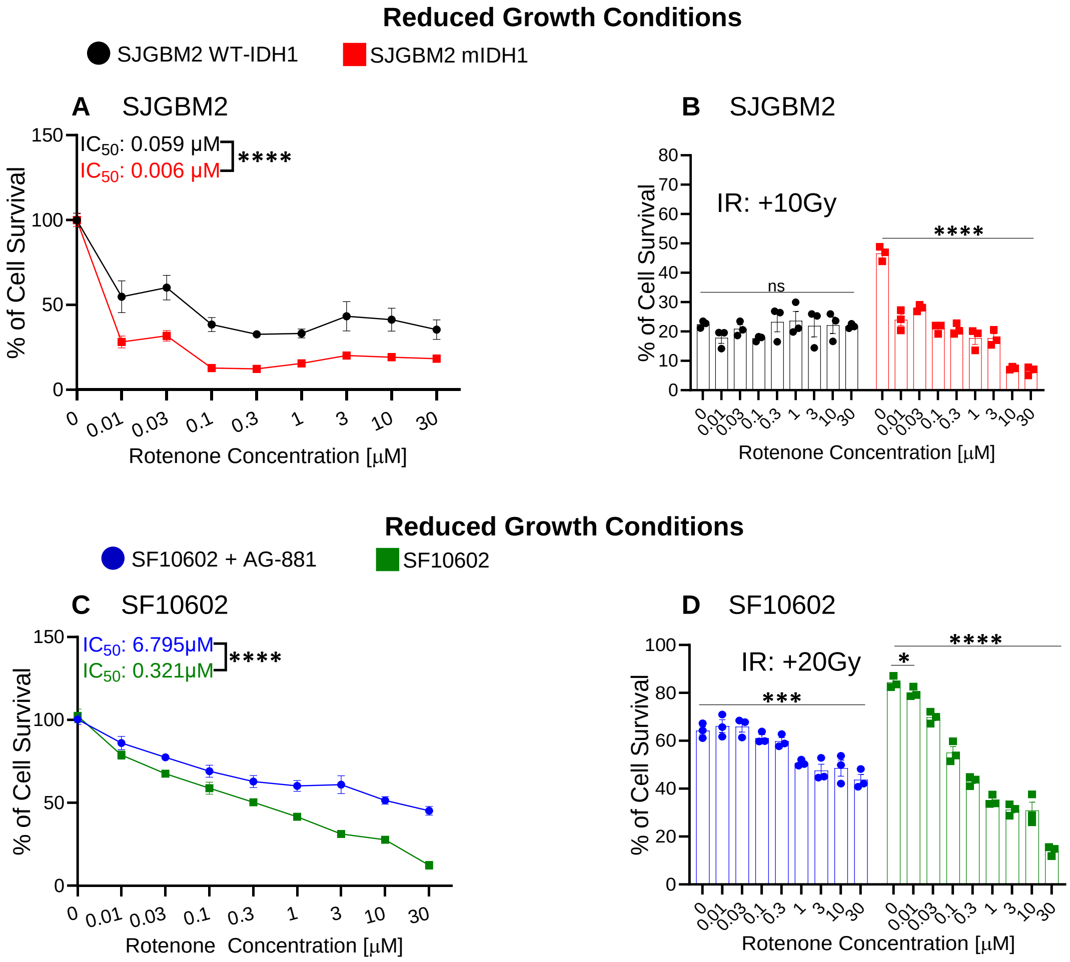


**Supplemental Figure 18:** In Vitro inhibition of Complex I of the ETC induces cell death and restores radiosensitivity under reduced growth conditions in two human mIDH1 glioma cells. (**A**) Cell viability assay of WT-IDH1 and mIDH1 genetically engineered human glioma cells, SJGBM2, under reduced growth conditions, treated with rotenone for 72 hours at increasing concentrations. Cell viability expressed as % of cell survival relative to control populations. A linear regression model and generalized additive models were used to determine the dose-response relationship. (**B**) Cell viability assay showing the effect of rotenone at increasing concentrations in combination with radiation (10Gy) on the proliferation of WT-IDH1 and mIDH1 SJGBM2 cells under reduced growth conditions. (**C**) Cell viability assay of endogenous mIDH1 glioma cells, SF10602, treated with and without mIDH1 inhibitor, AG-881, under reduced growth conditions, were treated with rotenone for 72 hours at increasing concentrations. Cell viability expressed as % of cell survival relative to control populations. A linear regression model and generalized additive models were used to determine the dose-response relationship. (**D**) Cell viability assay showing the effect of rotenone at increasing concentrations in combination with radiation (20Gy) on the proliferation of AG-881 treated SF10602 and untreated SF10602 cells under reduced growth conditions. Results are expressed as % of Cell survival relative to control populations. Statistical analyses were conducted using student's t-test and error bars depict SD of technical replicates from 3 independent experiments. **P* < 0.1, ***P* < 0.01, ****P* < 0.001, *****P* < 0.0001, ns = not significant.

**Supplemental Figure 19**

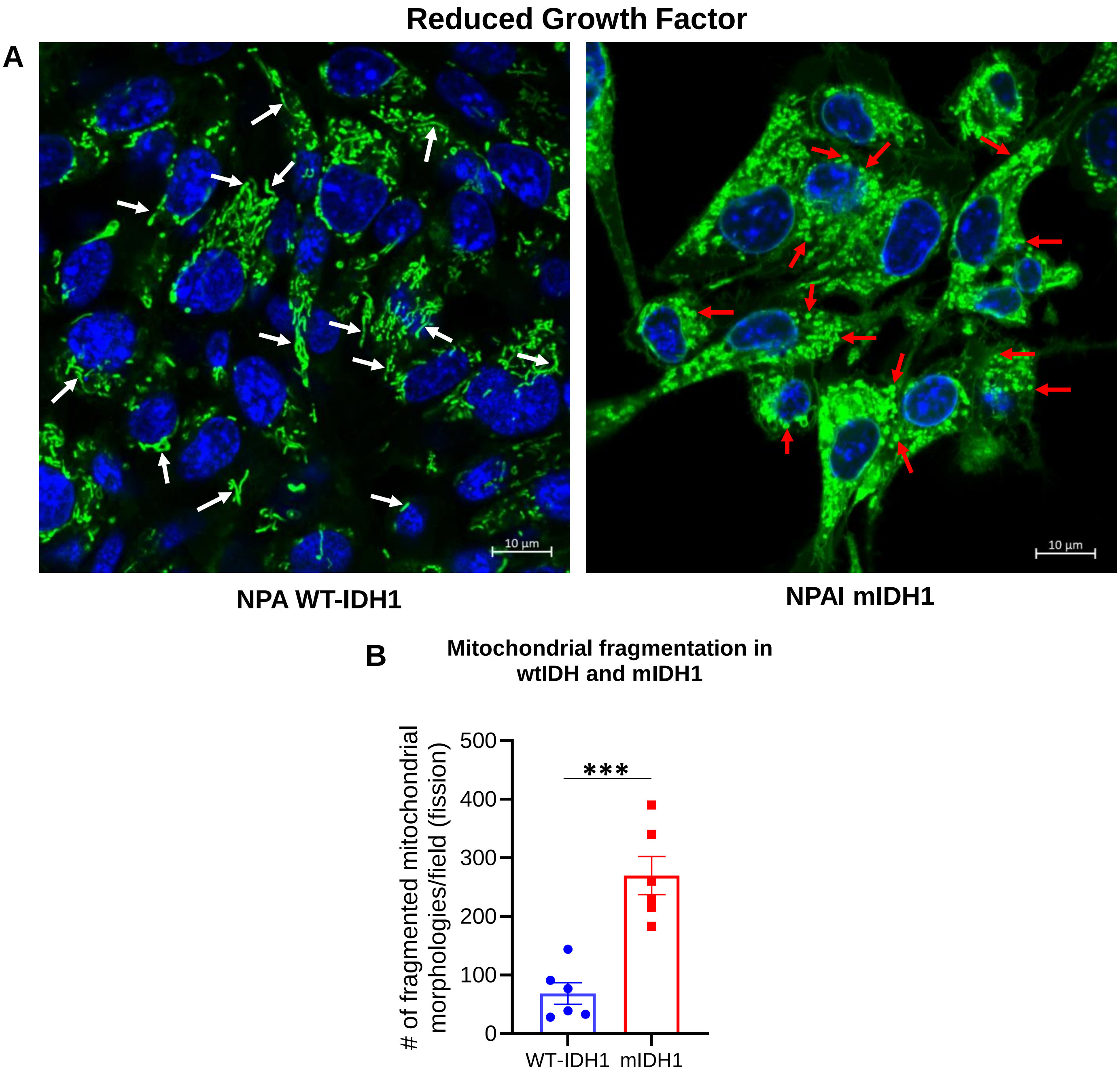


**Supplemental Figure 19:** Mitochondrial morphology of WT-IDH1 NPA NS and mIDH1 NPAI NS under growth factor starved conditions. (**A**) Confocal microscopy of mitochondrial morphology in WT-IDH1 NS, and mIDH1 NS, labeled with MitoTracker Green FM under growth factor starved conditions (final concentration: EGF: 6.5 ng/mL; FGF: 6.5 ng/mL]. White arrows indicate mitochondrial fusion; red arrows indicate mitochondrial fission. Scale bar: 10µm. (**B**) Quantification of fragmented mitochondrial morphologies in mouse WT-IDH1 and mIDH1 NS using Image J2 software. Standard settings were applied for the Mitochondrial Morphology Macro, the MitoLoc plugin and the MiNa plugin. For the Particle Analyzer method, auto thresholding was applied on images before applying the Analyze particles function. Confocal images were deconvoluted using the Iterative Deconvolve 3D plugin, without a wiener filter gamma and with a low pass filter of 1, a maximum number of iterations of 10 and a termination of iterations at 0.010. Data were collected from three independent experiments. ***$P < 0.001$.

**Supplemental Figure 20**

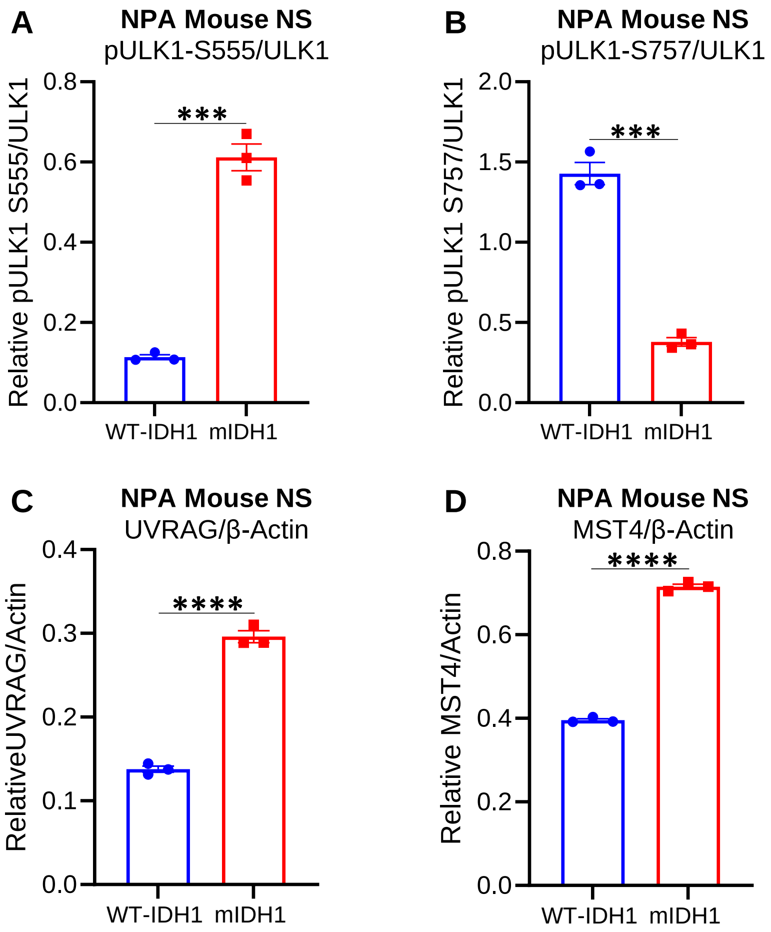


**Supplemental Figure 20:** Densiometric analysis of Mouse NS Western blots. ImageJ densitometric quantification of the western blots of (**A**) pULK1-S555, (**B**) pULK1-S757, (**C**) UVRAG, and (**D**) MST4. Errors bars represent SEM from independent technical replicates (n = 3). ****P* < 0.001; *****P* < 0.0001; unpaired t test.

**Supplemental Figure 21**

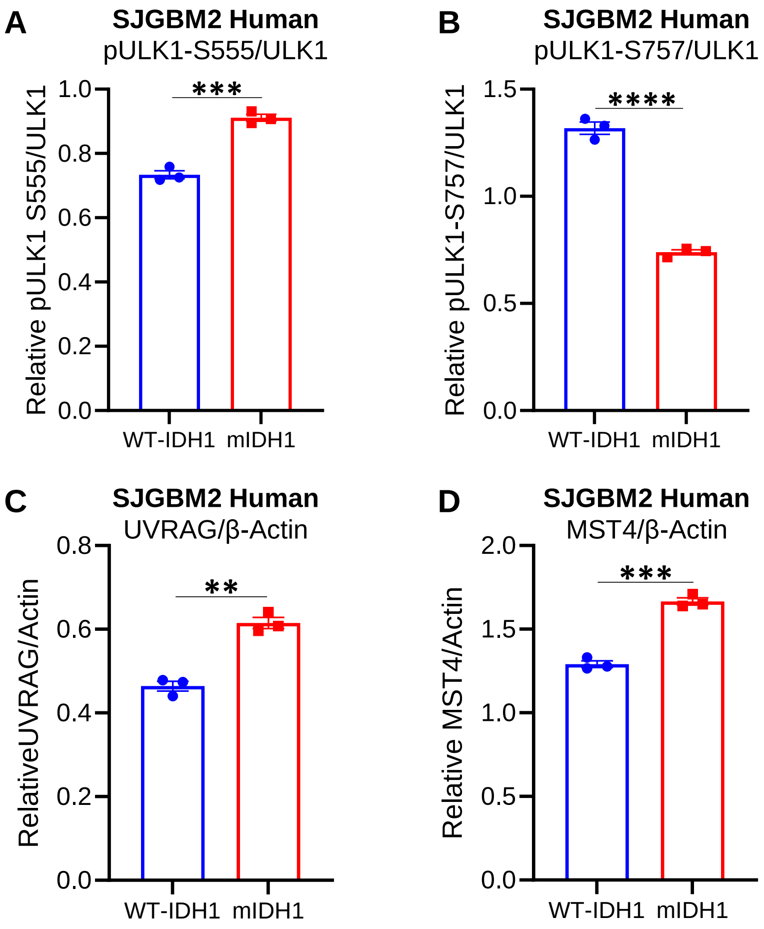


**Supplemental Figure 21:** Densiometric analysis of SJGBM2 Western blots. ImageJ densitometric quantification of the western blots of (**A**) pULK1-S555, (**B**) pULK1-S757, (**C**) UVRAG, and (**D**) MST4. Errors bars represent SEM from independent technical replicates (n = 3). ***P* < 0.005; ****P* < 0.001; unpaired t test.

**Supplemental Figure 22**

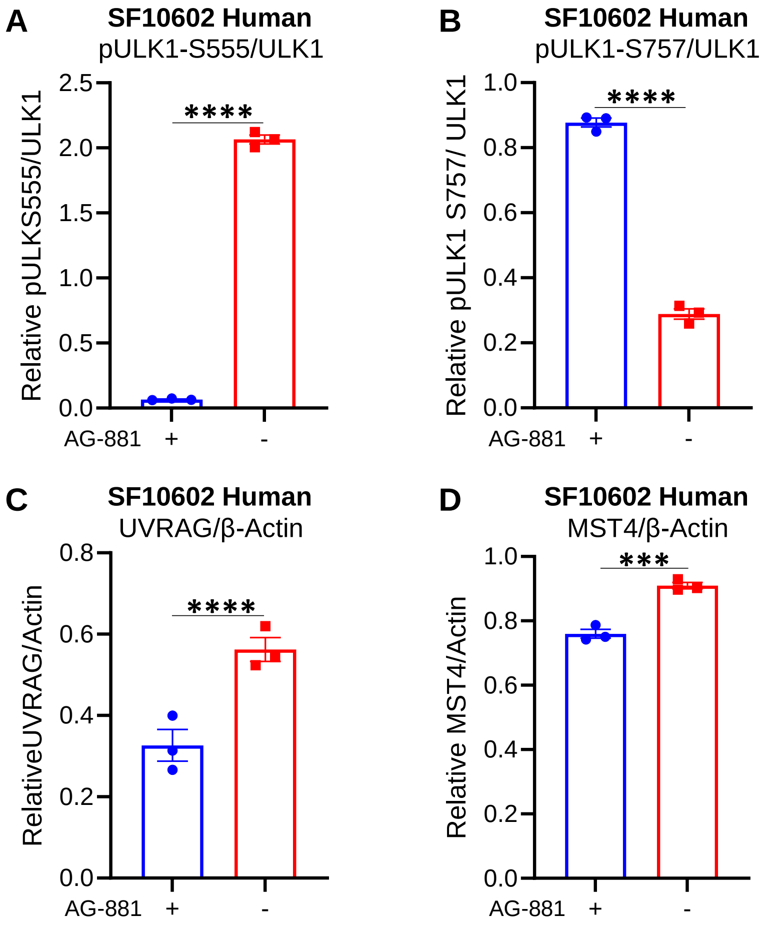


**Supplemental Figure 22:** Densiometric analysis of SF10602 ± AG-881 Western blots. ImageJ densitometric quantification of the western blots of (**A**) pULK1-S555, (**B**) pULK1-S757, (**C**) UVRAG, and (**D**) MST4. Errors bars represent SEM from independent technical replicates (n = 3). ***$P$ < 0.001; ****$P$ < 0.0001; unpaired t test.

**Supplemental Figure 23**

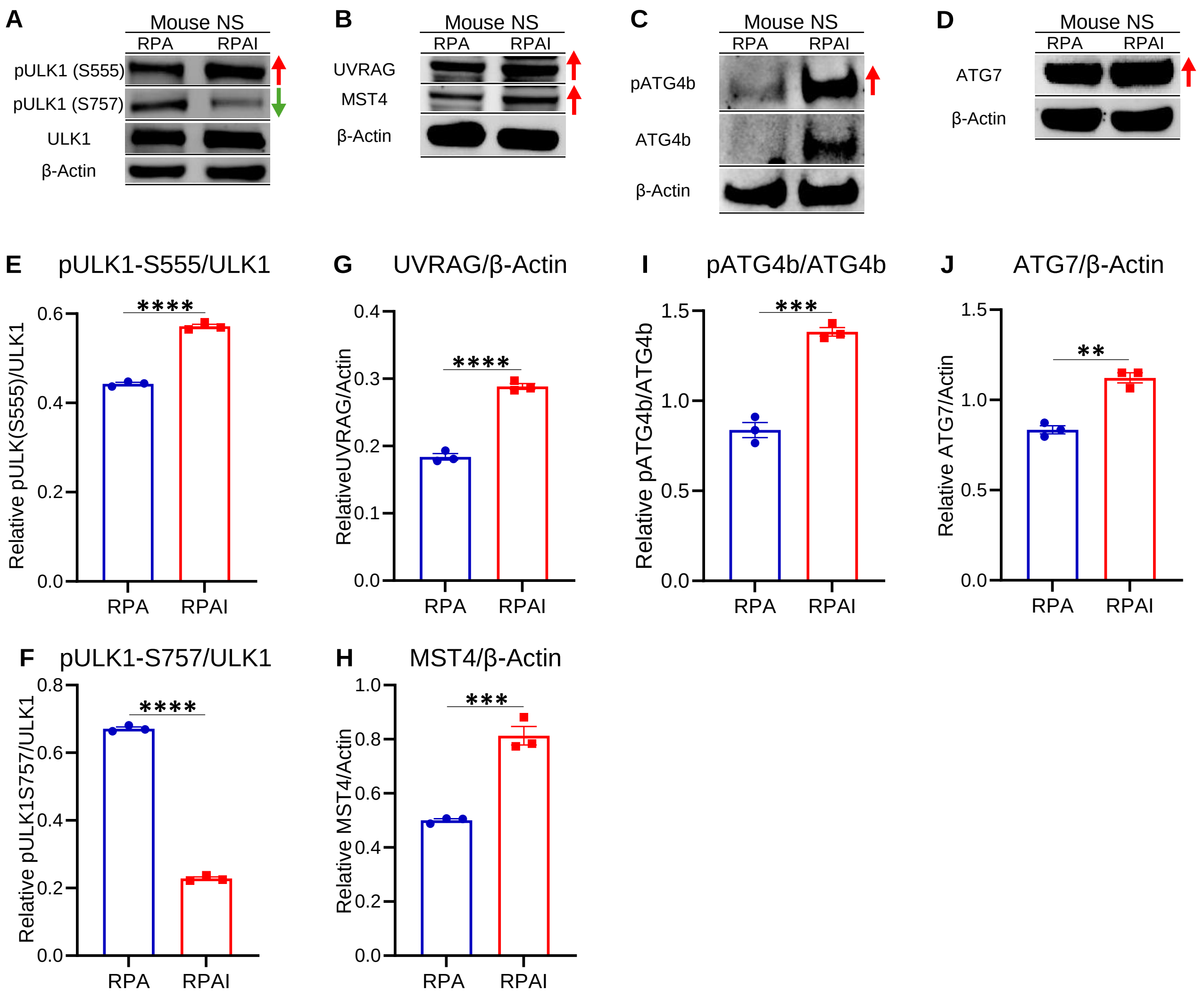


**Supplemental Figure 23:** Western Blot analysis of Autophagy related genes in PDGFRα-D842V driven genetically engineered mouse model. (**A-D**) Western blot analysis of autophagy related genes (A) pULK1-S555, pULK1-S757, ULK1; (B) UVRAG, MST4; (C) pATG4b, ATG4b; (D) ATG7 in genetically engineered mouse models (i) RPA: PDGFRα-D842V, shTP53, and shATRX; and (ii) RPAI: PDGFRα-D842V, shTP53, shATRX, and IDH1-R132H. The red arrows indicate the proteins and phosphorylated (p) status related with autophagy activation (pULK1 (S555), UVRAG, MST4, pATG4B (S383), ATG7) whereas the green arrows indicate the phosphorylated (p) status related with autophagy inhibition (pULK1 (S757); β-actin, loading control. (**E-J**) ImageJ densitometric quantification of the western blots for (E) relative pULK-S555/ULK1, (F) relative pULK1-S757/ULK1, (G) relative UVRAG/β-Actin, (H) relative MST4/β-Actin, (I) relative pATG4b/ATG4b, and (J) relative ATG7/β-Actin. Error bars represent SEM from independent technical replicates (n = 3). $^{**}P < 0.005$; $^{***}P < 0.001$; $^{****}P < 0.0001$; unpaired t test.

**Supplemental Figure 24**

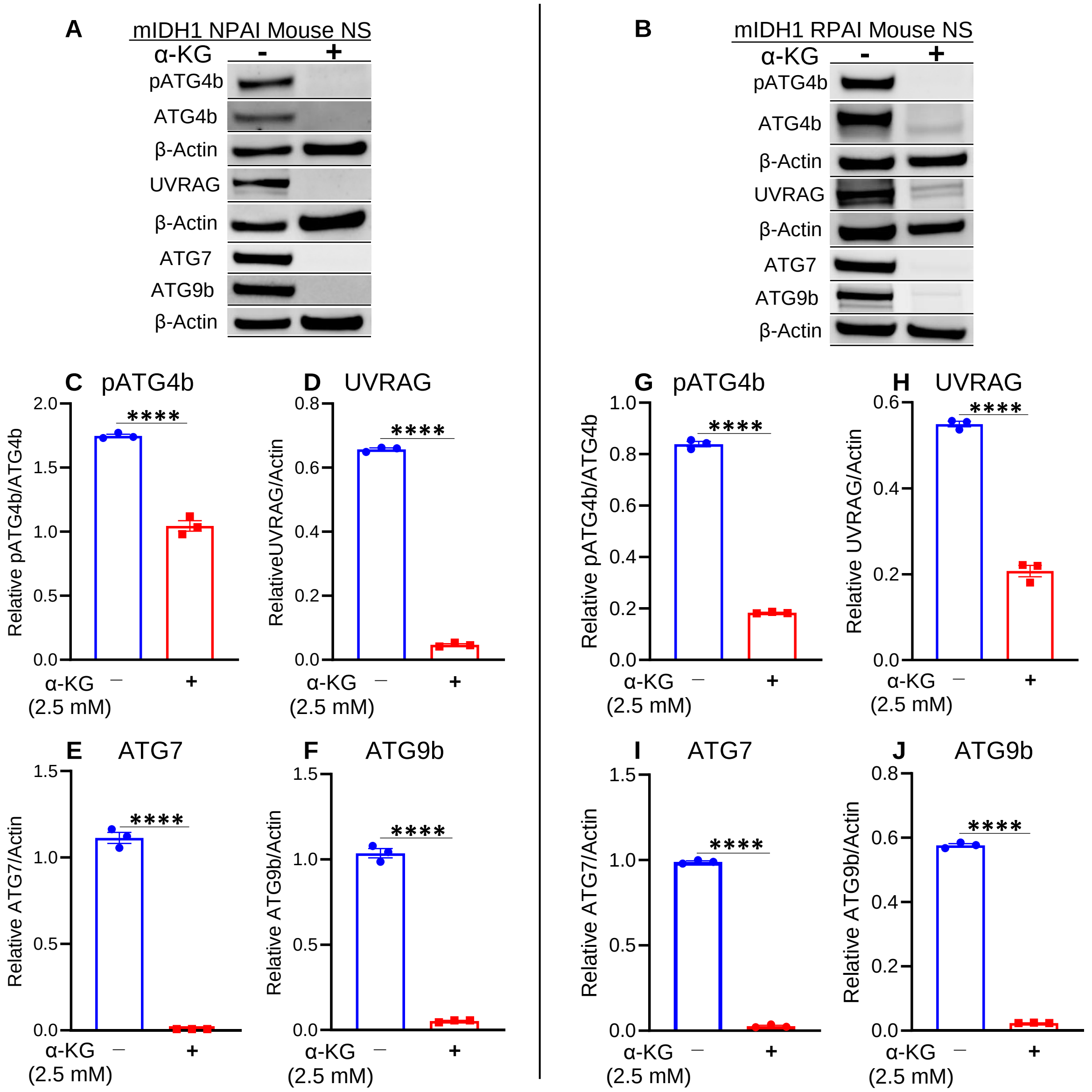


**Supplemental Figure 24:** Western blot measurement of proteins involved in autophagy pathway of either NPAI or RPAI mIDH1 mouse NS treated with α-ketoglutarate. (**A-B**) WB assay showing the expression of proteins involved in the autophagy pathways and the activated form of them, evaluated in (A) mIDH1 NPAI mouse NS ± treatment with α-ketoglutarate (α-KG) or (B) mIDH1 RPAI mouse NS ± treatment with α-KG. β-actin, loading control. (**C-F**) ImageJ densitometric quantification of western blot assays of (C) pATG4b, (D) UVRAG, (E) ATG9b, and (F) ATG7 in the mIDH1 NPAI mouse NS treated with α-KG. (**G-J**) ImageJ densitometric quantification of western blot assays of (G) pATG4b, (H) UVRAG, (I) ATG9b, and (J) ATG7 in the mIDH1 RPAI mouse NS treated with α-KG . Errors bars represent SEM from independent technical replicates (n = 3). ****$P$ < 0.0001; unpaired t test.

**Supplemental Figure 25**

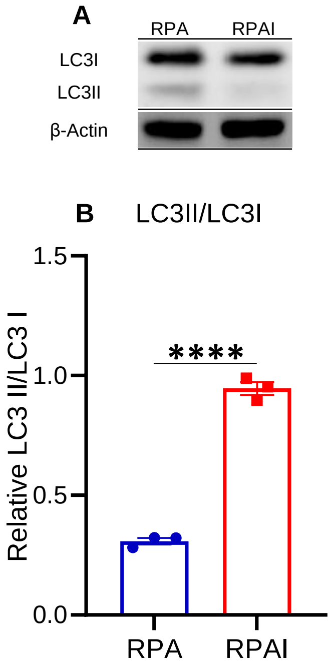


**Supplemental Figure 25:** Western Blot analysis of Autophagy related genes in PDGFRα-D842V driven genetically engineered mouse model. (**A-D**) Western blot analysis of autophagy related genes (A) pULK1-S555, pULK1-S757, ULK1; (B) UVRAG, MST4; (C) pATG4b, ATG4b; (D) ATG7, and LC3I/II in genetically engineered mouse models (i) RPA: PDGFRα-D842V, shTP53, and shATRX; and (ii) RPAI: PDGFRα-D842V, shTP53, shATRX, and IDH1-R132H. The red arrows indicate the proteins and phosphorylated (p) status related with autophagy activation (pULK1 (S555), UVRAG, MST4, pATG4B (S383), ATG7) whereas the green arrows indicate the phosphorylated (p) status related with autophagy inhibition (pULK1 (S757); β-actin, loading control. (**E-K**) ImageJ densitometric quantification of the western blots for (E) relative pULK-S555/ULK1, (F) relative pULK1-S757/ULK1, (G) relative UVRAG/β-Actin, (H) relative MST4/β-Actin, (I) relative pATG4b/ATG4b, (J) relative ATG7/β-Actin, and (K) relative LC3II/LC3I. Error bars represent SEM from independent technical replicates (n = 3). ****$P < 0.0001$; unpaired t test.

**Supplemental Figure 26**

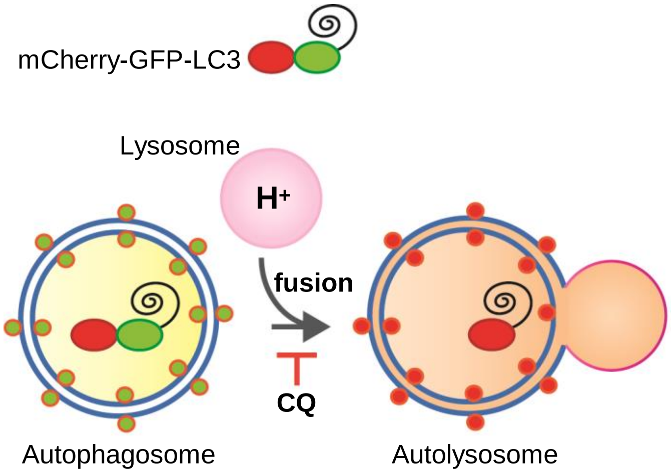


**Supplemental Figure 26:** Autophagy flux assay using a lentivirus to express LC3 protein associated with GFP and mCherry reporter protein. In cells expressing mCherry-GFP-LC3, autophagosomes display both GFP and mCherry fluorescence, whereas autolysosomes display only mCherry fluorescence (red) because GFP is denatured by the acidity of the lysosome. Chloroquine (CQ) inhibits the fusion step in autophagy flux, disrupting autolysosome formation.

**Supplemental Figure 27**

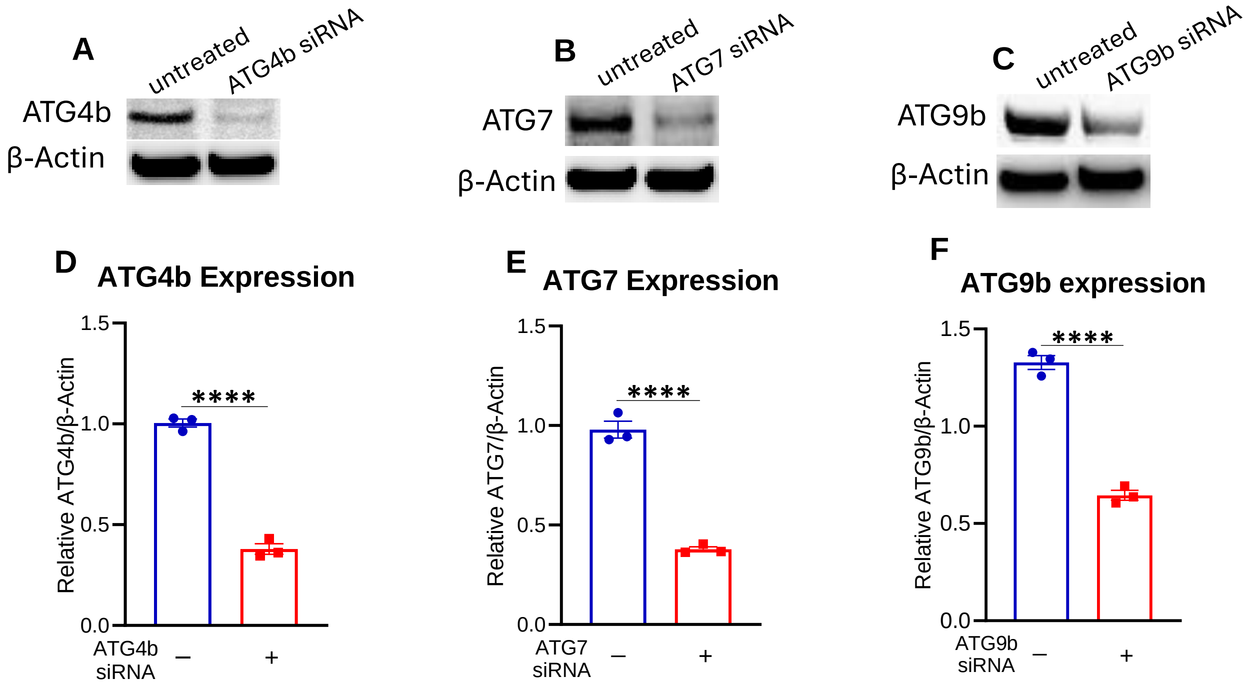


**Supplemental Figure 27:** Validation of ATG4b, ATG7, and ATG9b siRNAs. (**A-C**) Western blot analysis showing successful inhibition of (A) ATG4b, (B) ATG7, and (C) ATG9b via siRNA in mIDH1 mouse glioma NS with β-actin as a loading control. (**D-F**) ImageJ densitometric quantification of the western blot for (D) relative ATG4b/β-actin, (E) relative ATG7/β-actin, and (F) relative ATG9b/β-actin. Errors bars represent SEM from independent biological replicates (n = 3). ****$P < 0.0001$; unpaired t test.

**Supplemental Figure 28**

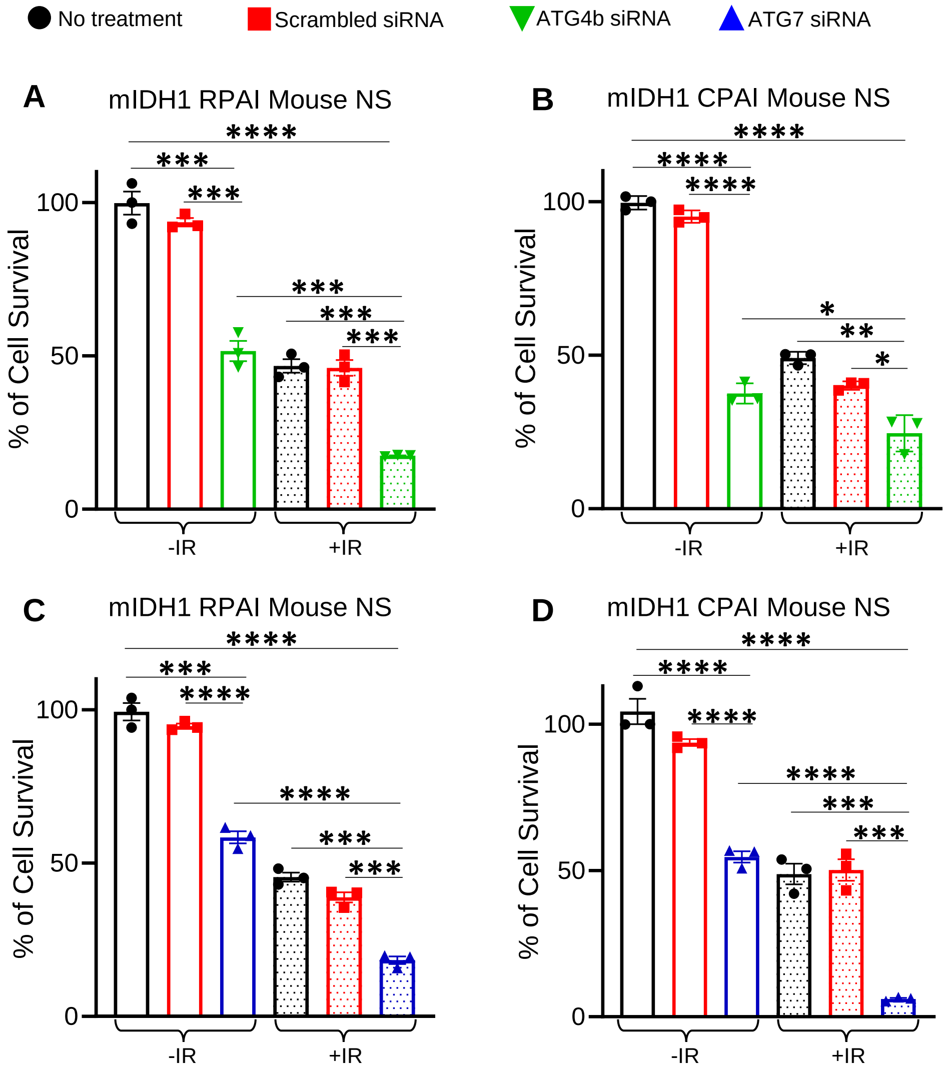


**Supplemental Figure 28:** Silencing autophagy induces cytotoxicity and restores radiosensitivity in two genetically engineered mouse glioma cell models: RPAI and CPAI. Cell Titer Glo assays (luminescence) show the impact of (**A-B**) ATG4b silencing (ATG4b siRNA) and (**C-D**) ATG7 silencing (ATG7 siRNA) on cellular cytotoxicity and radiosensitivity on (A, C) mIDH1 RPAI mouse NS, (B, D) mIDH1 CPAI mouse NS. Cells were treated with saline, scrambled siRNA, free ATG4b siRNAs (A-B), or free ATG7 siRNAs (C-D). The amount of ATG4b siRNA or ATG7 siRNA used was 100 nM. The amount of radiation used was 3 Gy. All experiments were performed in quadruplicate, and results are expressed as "% cell survival". Statistical analyses were conducted using student's t test and error bars depicting the SEM of technical replicates from 3 independent experiments. *$P < 0.1$, **$P < 0.01$, ***$P < 0.001$, ****$P < 0.0001$.

**Supplemental Figure 29**

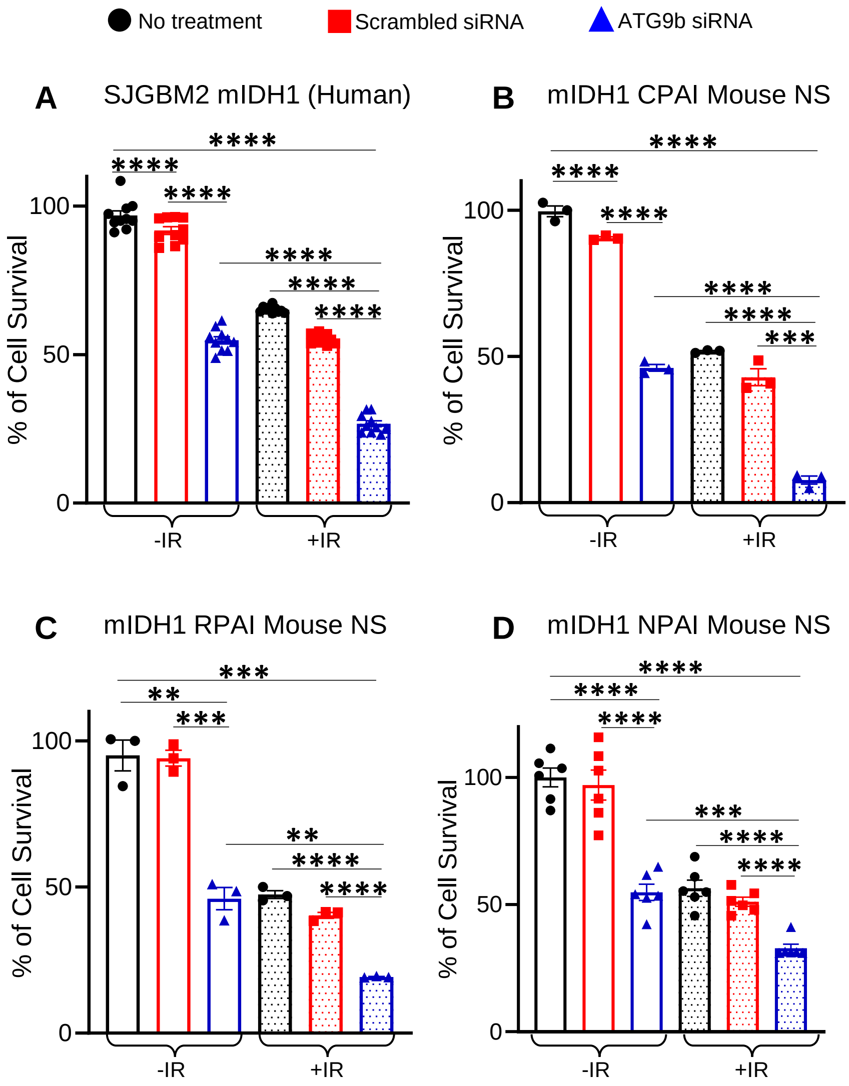


**Supplemental Figure 29:** Silencing autophagy induces cytotoxicity and restores radiosensitivity in both patient-derived glioma cells and genetically engineered mouse glioma cells. Cell Titer Glo assays (luminescence) show the impact of ATG9b silencing (ATG9b siRNA) on cellular cytotoxicity and radiosensitivity on (**A**) SJGBM2 mIDH1 human cells, (**B**) mIDH1 CPAI mouse NS, (**C**) mIDH1 RPAI mouse NS, and (**D**) mIDH1 NPAI mouse NS. Cells were treated with saline, scrambled siRNA, or free ATG9b siRNA. The amount of ATG9b siRNA used was 100nM for all mouse cells and 150nM for all human cells. The amount of radiation used was 5 Gy for SJGBM2 cells and 3 Gy for mouse NS. All experiments were performed in quadruplicate, and results are expressed as “% cell survival”. Statistical analyses were conducted using student’s t test and error bars depicting the SEM of technical replicates from 3 independent experiments. $^{**}P < 0.01$, $^{***}P < 0.001$, $^{****}P < 0.0001$.

**Supplemental Figure 30**

A

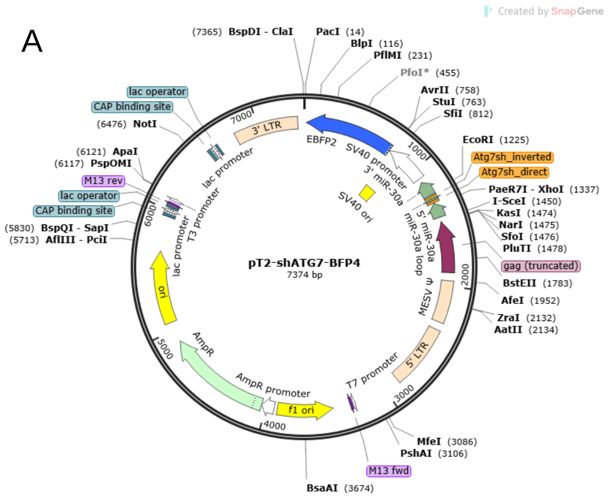


B

| shRNA | Sequences |
|---|---|
| sh2-ATG7 F | TCGAGAAGGTATATTGCTGTTGACAGTGAGCGCCAGAAGAAGTTGAACGAGTATATAGTGAAGCCACAGATGTATATACTCGTTCAACTTCTTCTGGTGCCTACTGCCTCGG |
| sh2-ATG7 R | AATTCCGAGGCAGTAGGCACCAGAAGAAGTTGAACGAGTATATACATCTGTGGCTTCACTATATACTCGTTCAACTTCTTCTGGCGCTCACTGTCAACAGCAATATACCTTC |

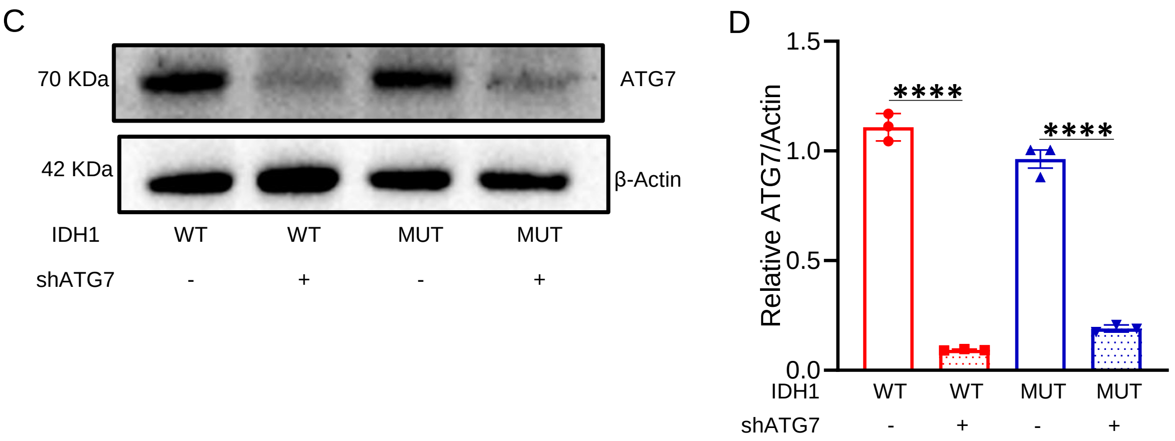


**Supplemental Figure 30:** Generation of autophagy deficient mIDH1 mouse glioma cells. (**A**) Sleeping Beauty (SB) plasmid pT2-shATG7-BFP4, containing a short hairpin targeting in *Atg7*, was used for in vitro and in vivo knockdown of *Atg7* expression in mIDH1 glioma using the SB transposon system. The plasmid map illustrates the final construction, which encodes for shATG7 (HP_603911) and enhanced blue fluorescent protein (EBFP). (**B**) Corresponding shRNA sequences highlighted in red for shAtg7. (**C, D**) Western blot analysis showing successful knockdown of ATG7 within wtIDH1 cells (WT) and mIDH1 cells (MUT) with β-actin as a loading control. ImageJ densitometric quantification of the western blot for ATG7 and β-actin. Errors bars represent SEM from independent biological replicates (n = 3). ****$p < 0.0001$; unpaired t test.

**Supplemental Figure 31**

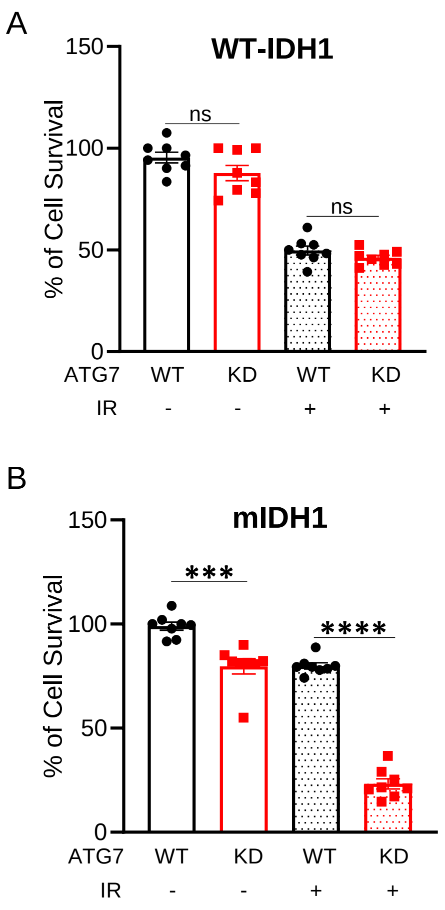


**Supplemental Figure 31:** Silencing autophagy restores radiosensitivity in mIDH1 mouse NS. Cell proliferation assays show the impact of ATG7 knockdown (ATG7 KD) on radiosensitivity in (**A**) NRAS/shP53/shATRX (NPA) mouse NS and (**B**) NRAS/shP53/shATRX/mIDH1 (NPAI) mouse NS, when treated with 3 Gy of radiation. All the experiments were performed in octuplicate (8 replicates), and the results are expressed in “% cell survival”. Statistical analyses were done with student’s t test and error bars depicting the SD of technical replicates from 3 independent experiments. ****P* < 0.001, *****P* < 0.0001, ns = not significant.

**Supplemental Figure 32**

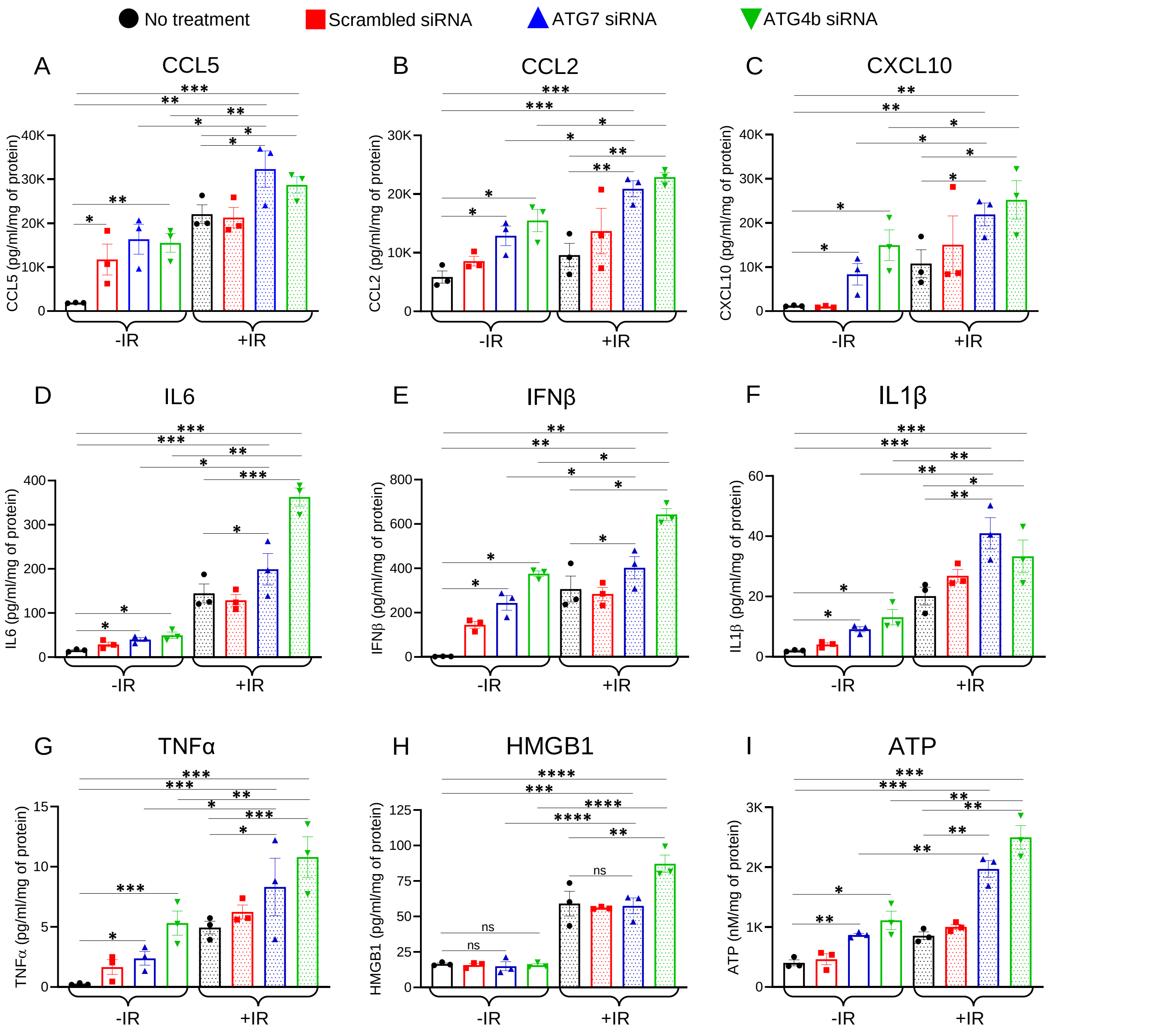


**Supplemental Figure 32:** Cytokine profiling in mIDH1 glioma mouse NS in response to autophagy inhibition and IR. The following cytokines were profiled in mIDH1 glioma mouse NS in response to autophagy inhibition and IR: Levels of DAMPs and Type-I IFNs i.e. (**A**) CCL5; (**B**) CCL2; (**C**) CXCL10; (**D**) IL6; (**E**) IFNβ; (**F**) IL1β; (**G**) TNFα; (**H**) HMGB1; and (**I**) ATP were determined through ELISA following treatment with 100nM of ATG7 and ATG4b siRNAs in combination with 3Gy ionizing radiation (IR). Conditioned media were collected and analyzed after 72h post-treatment. Statistical significances were determined using one-way ANOVA followed by Tukey's test. Significances denoted as follows: *$P$ < 0.05, **$P$ < 0.01, ***$P$ < 0.001, ****$P$ < 0.0001, ns = not significant. The bars represent the mean ± SEM (n = 3 biological replicates).

**Supplemental Figure 33**

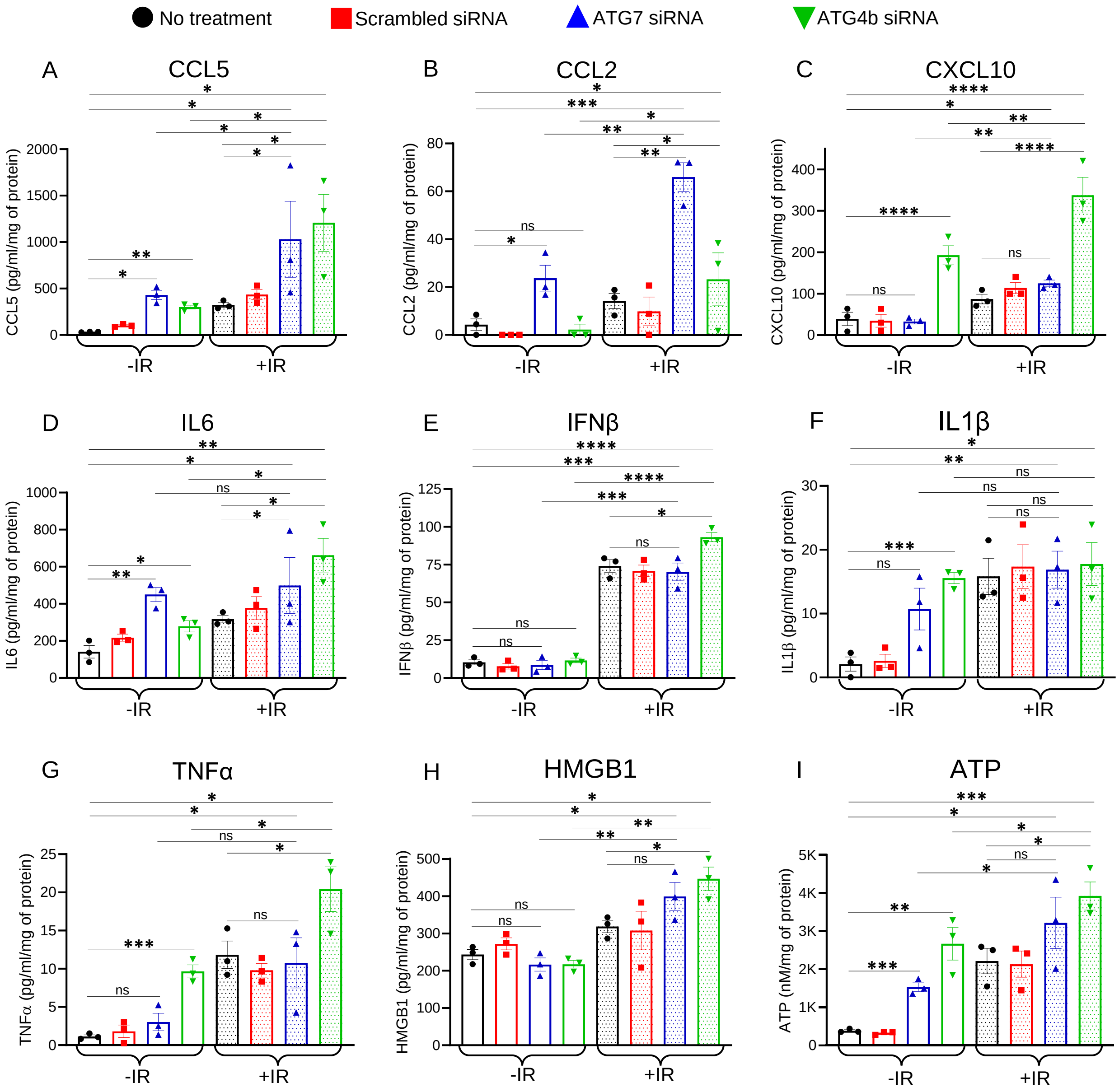


**Supplemental Figure 33:** Cytokine profiling in mIDH1 SJGBM2 cells in response to autophagy inhibition and IR. The following cytokines were profiled in human mIDH1 glioma cells in response to autophagy inhibition and IR: Levels of DAMPs and Type-I IFNs i.e. (**A**) CCL5; (**B**) CCL2; (**C**) CXCL10; (**D**) IL6; (**E**) IFNβ; (**F**) IL1β; (**G**) TNFα; (**H**) HMGB1; and (**I**) ATP were determined through ELISA following treatment with 150nM of ATG7 and ATG4b siRNAs in combination with 5Gy IR. Conditioned media were collected and analyzed after 72h post-treatment. Statistical significances were determined using one-way ANOVA followed by Tukey's test. Significances denoted as follows: *$P < 0.05$, **$P < 0.01$, ***$P < 0.001$, ****$P < 0.0001$, ns = not significant. The bars represent the mean ± SEM (n = 3 biological replicates).

**Supplemental Figure 34**

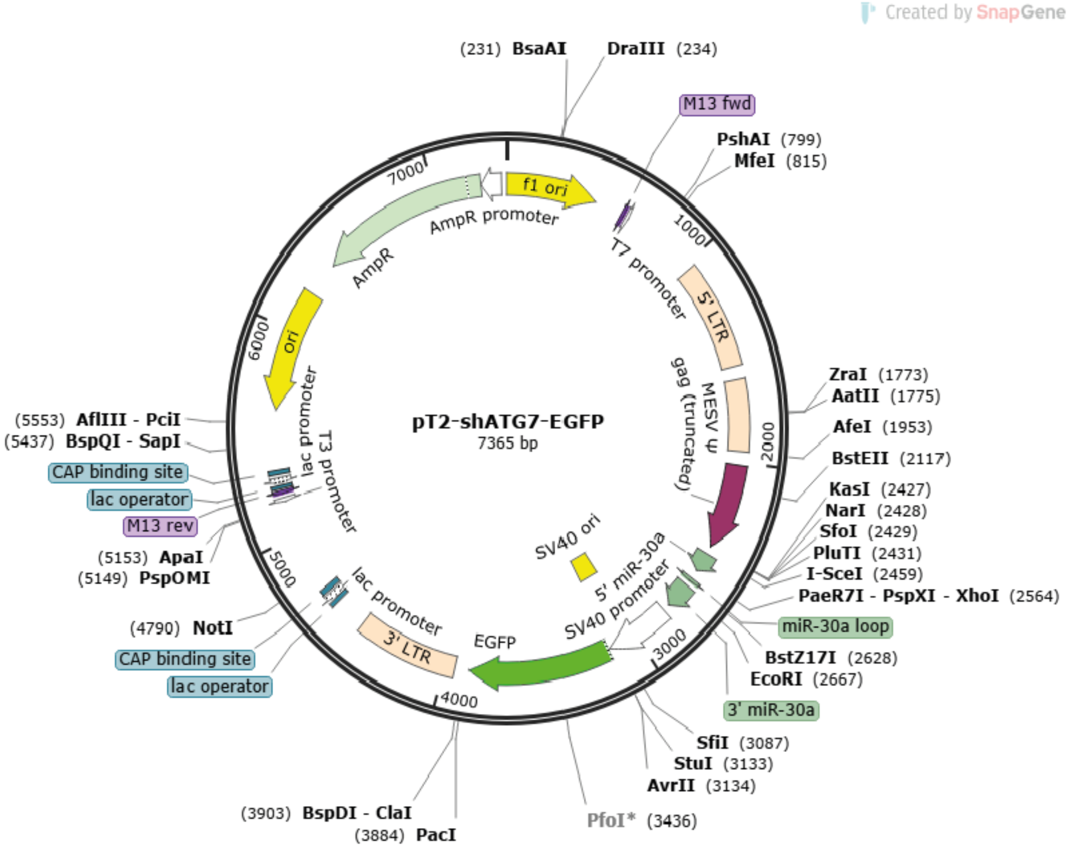


shATG7 (HP_603911):
CTCGAGTGCTGTTGACAGTGAGCGACCAGAAGAAGTTGAACGAGTATAGTGAAGCCACAGATGTATACTCTTCAACTTCTTT
GGGTGCCTACTCCTCGGAGAATTC

**Supplemental Figure 34:** Sleeping beauty plasmid pT2-shATG7-EGFP. Generation of an autophagy deficient mIDH1 mouse glioma model. Sleeping Beauty (SB) plasmid pT2-shAtg7-GFP, containing a short hairpin targeting in *Atg7*, was used for in vivo knockdown of *Atg7* expression in mIDH1 glioma using the SB transposon system. The plasmid map illustrates the final construction, which encodes for shATG7 (HP_603911) and EGFP.

**Supplemental Figure 35**

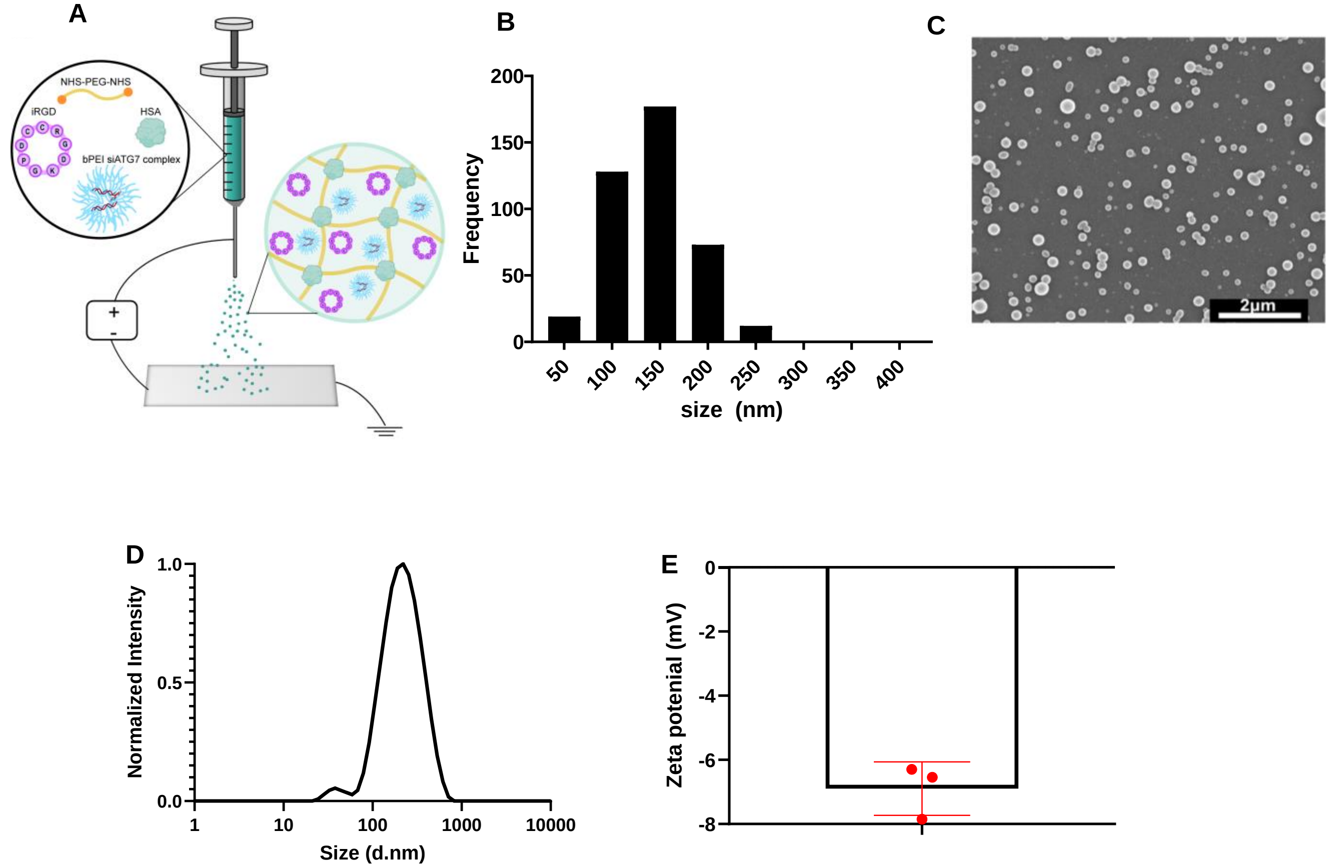


**Supplemental Figure 35:** ATG7i-SPNP fabrication and characterization. (**A**) Electrohydrodynamic jetting to produce ATG7i-SPNPs. All components (lysine-containing iRGD, branched PEI-siATG7 complex, PEG crosslinker, BSA Alexa Fluor 647 conjugate) are added in the initial solution. Solid SPNPs result on the plate and are crosslinked for 7 days. (**B**) Histogram of dry state SPNP diameter (mean diameter: 104 ± 42 nm) analyzed from Scanning Electron Microscopy of n > 400 individual SPNPs. (**C**) Scanning electron microscopy micrograph (scale bar = 2 µm). (**D**) Intensity-based dynamic light scattering spectra ($Z_{average}$=185 nm, peak diameter = 238 nm, PDI = 0.289). (**E**) Zeta potential (-6.84 ± 0.69 mV).

**Supplemental Figure 36**

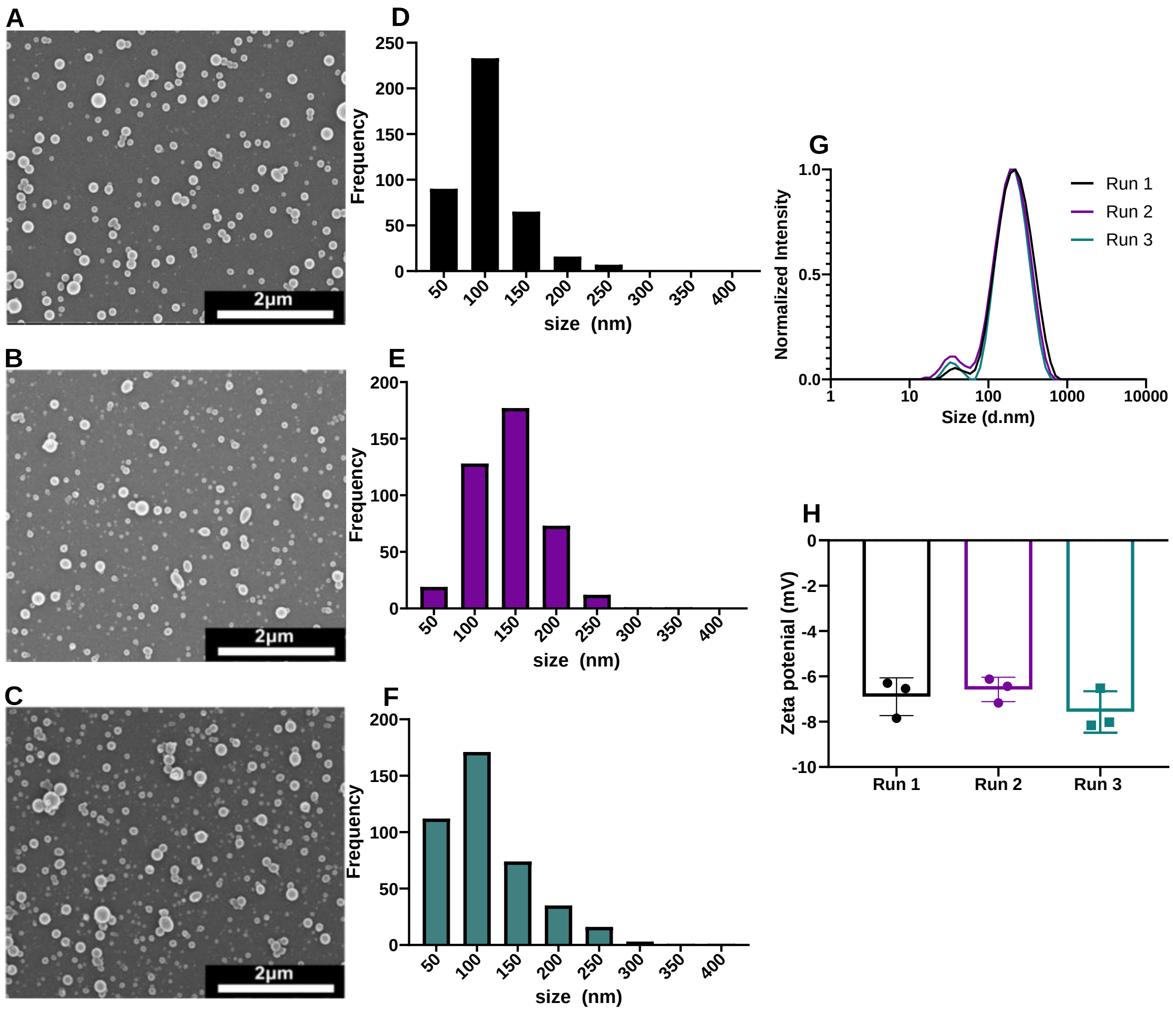


**Supplemental Figure 36:** Batch to batch production of ATG7i-SPNPs. Comparison of ATG7i-SPNPs characteristics that were fabricated and collected in three independent sessions. (**A-F**) A scanning electron microscope image (A-C) and the corresponding histogram (D-F) of the size distribution in the dry state of (A, D) run 1, (B, E) run 2, and (C, F) run 3. (**G**) Intensity dynamic light scattering spectra post-collection of the three runs. (**H**) Zeta potential.

**Supplemental Figure 37**

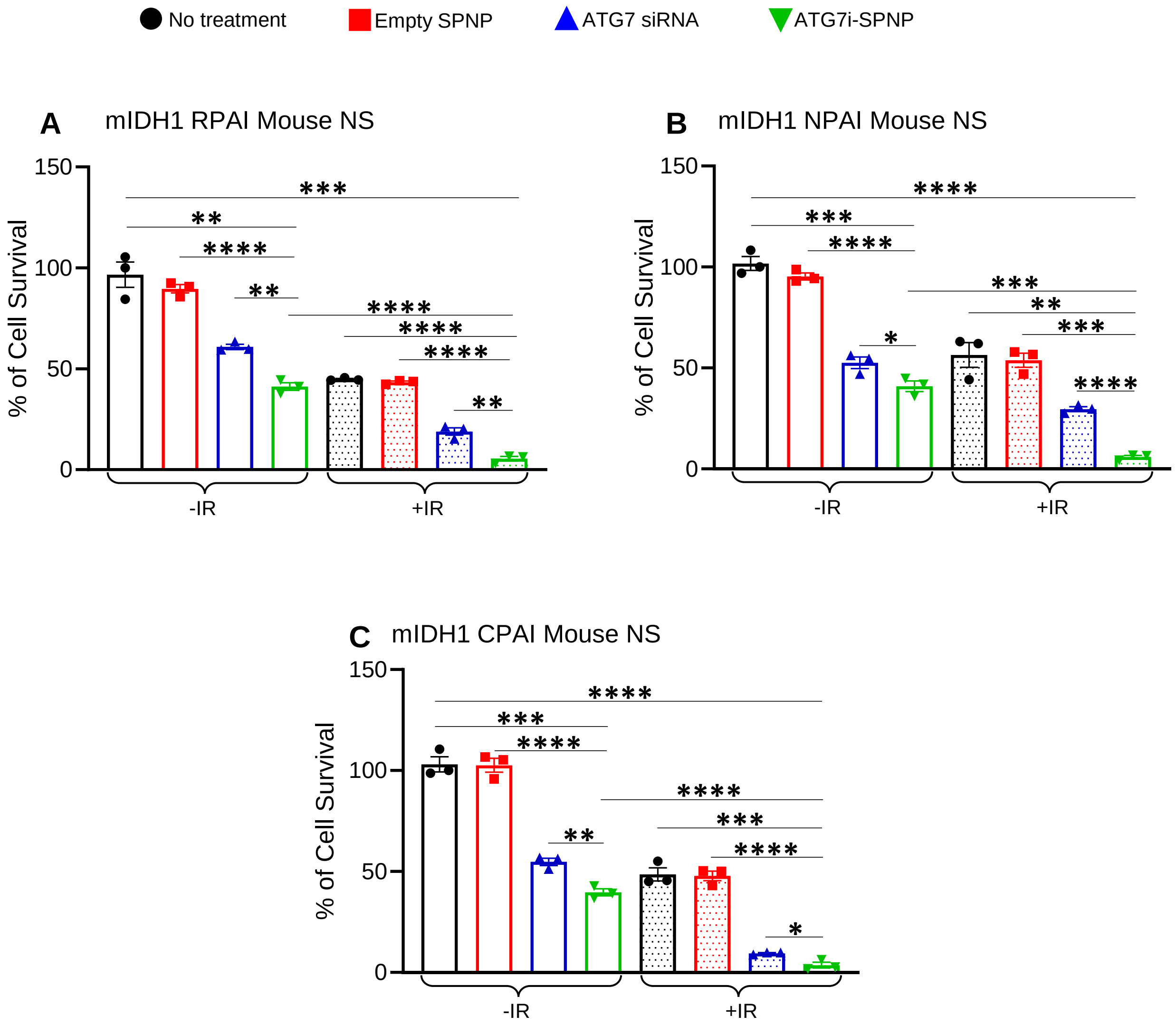


**Supplemental Figure 37:** Silencing autophagy induces cytotoxicity and restores radiosensitivity in genetically engineered mouse glioma cells. Cell Titer Glo assays (luminescence) show the impact of ATG7 silencing (ATG7 siRNA) on cellular cytotoxicity and radiosensitivity in (**A**) mIDH1 RPAI mouse NS, (**B**) mIDH1 NPAI mouse NS, and (**C**) mIDH1 CPAI mouse NS. Cells were treated with saline, empty nanoparticles (SPNPs), free ATG7 siRNAs, and ATG7 siRNA loaded nanoparticles (ATG7i-SPNPs). The amount of ATG7 siRNA used or loaded in the nanoparticle was 25µg. The amount of radiation used was 3 Gy. All experiments were performed in quadruplicate, and results are expressed as "% cell survival". Statistical analyses were conducted using student's t test and error bars depicting the SEM of technical replicates from 3 independent experiments. *$P < 0.1$, **$P < 0.01$, ***$P < 0.001$, ****$P < 0.0001$.

**Supplemental Figure 38**

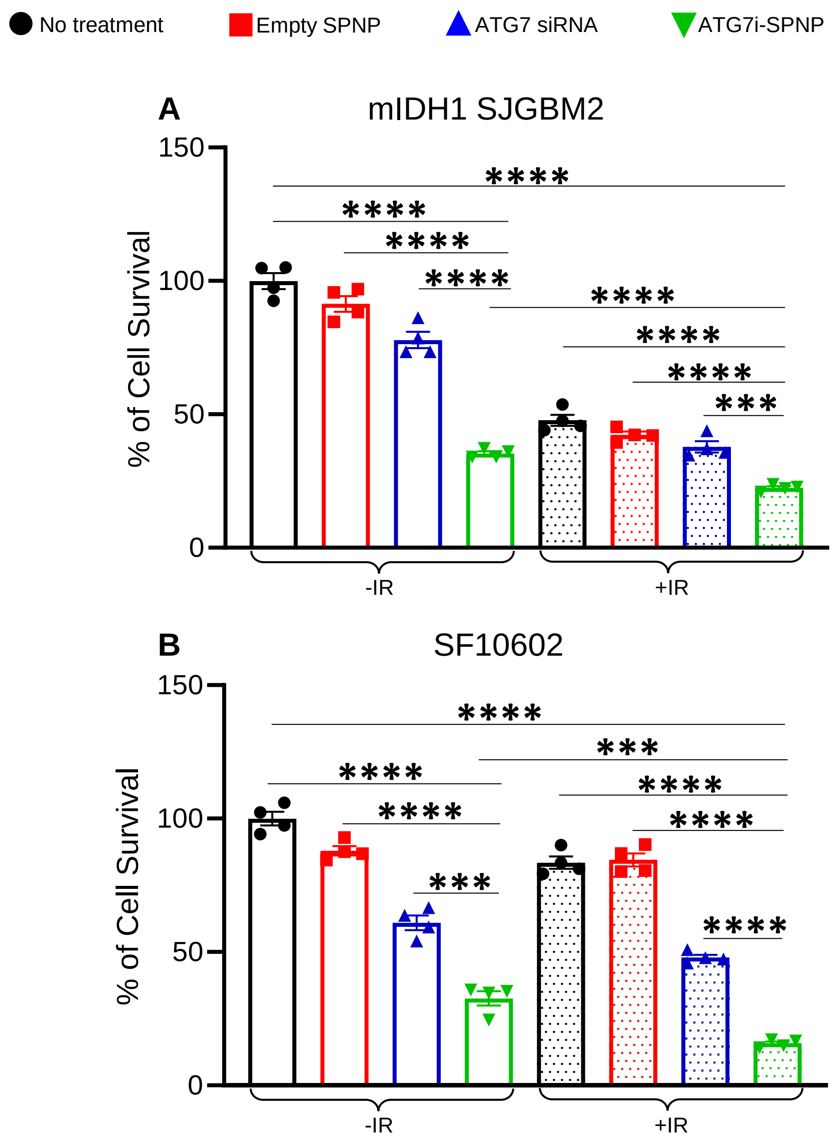


**Supplemental Figure 38:** Silencing autophagy induces cytotoxicity and restores radiosensitivity in mIDH1 human genetically engineered and patient-derived glioma cells. Cell Titer Glo assays (luminescence) show the impact of ATG7 silencing (ATG7 siRNA) on cellular cytotoxicity and radiosensitivity in (**A**) mIDH1 SJ-GBM2 following treatment with 10 Gy of irradiation. Similarly, the same luminescence assay was performed to evaluate the effect of ATG7 silencing on cellular cytotoxicity and radiosensitivity in (**B**) endogenous mIDH1-expressing human SF10602 cells, following 20 Gy of irradiation. Cells were treated with saline, empty nanoparticles (SPNPs), free ATG7 siRNAs, and ATG7 siRNA loaded nanoparticles (ATG7i-SPNPs). The amount of ATG7 siRNA used or loaded in the nanoparticle was 25µg. All experiments were performed in quadruplicate, and results are expressed as "% cell survival". Statistical analyses were conducted using student's t test and error bars depicting the SEM of technical replicates from 3 independent experiments. ***$P$ < 0.001, ****$P$ < 0.0001.

**Supplemental Figure 39**

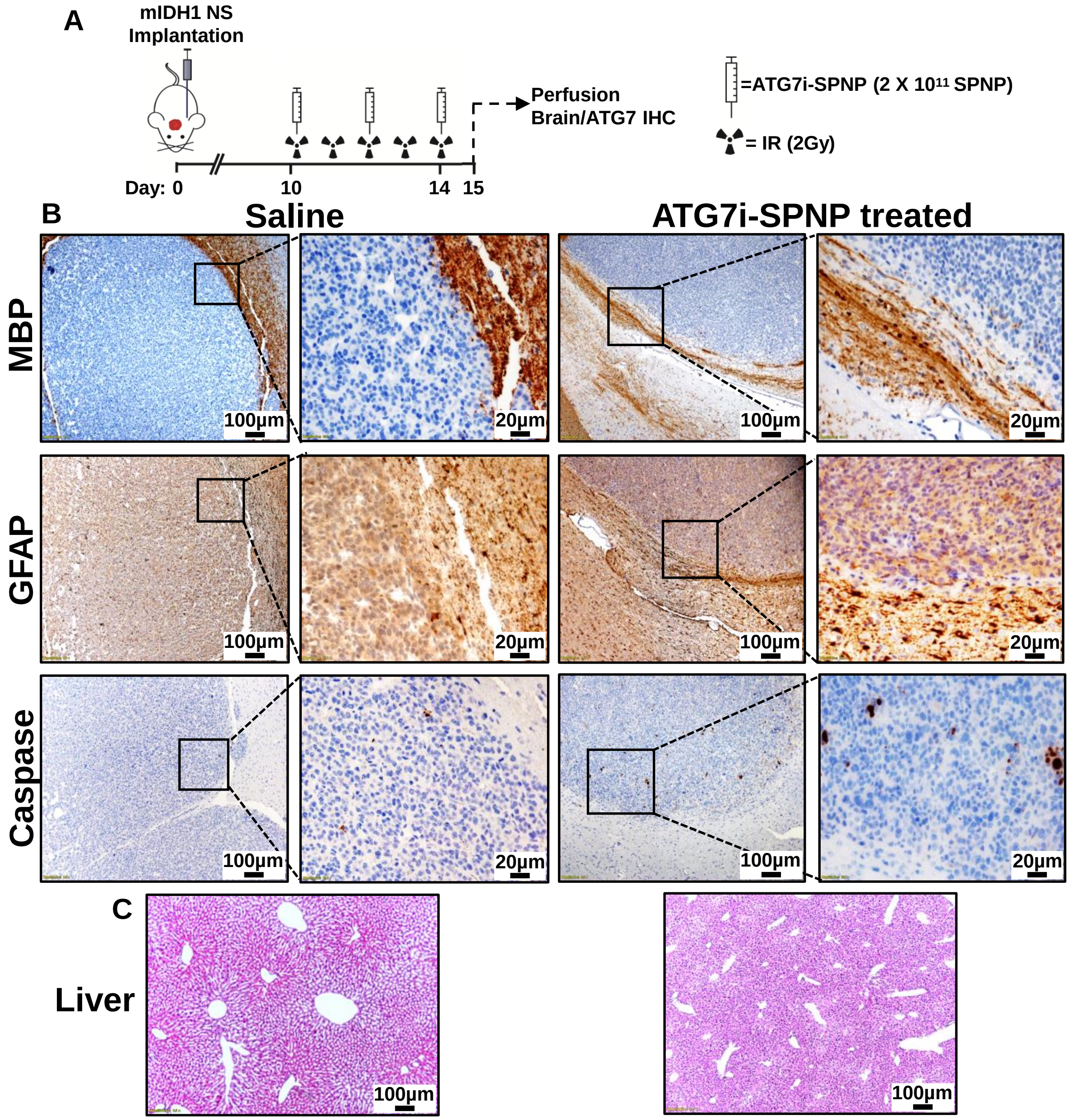


**Supplemental Figure 39:** Evaluation of brain and liver histopathology at an early time point following ATG7i-SPNP treatment in combination with IR in mIDH1 glioma-bearing mice. (**A**) Schematic of experimental design for the 14-day timed experiment: mIDH1 NS were implanted intracranially on Day 0. Mice were treated with ATG7i-SPNP on days 10, 12, and 14. Subsequently, mice were perfused at the end of Day 14 and analyzed. (**B**) Representative Immunohistochemical (IHC) staining of 5 µm paraffin-embedded brain sections from saline and ATG7i–SPNP treated groups, stained for MBP (myelin basic protein), GFAP (glial fibrillary acidic protein), and cleaved Caspase-3. MBP and GFAP staining are markers for regional demyelination and astrogliosis, respectively. Increased cleaved Caspase-3 staining in treated brains suggests enhanced apoptosis. Low magnification (10×) panels (black scale bar = 100 µm), High magnification (40×) panels (black scale bar = 20 µm) indicate positive staining for the areas in the low-magnification panels (**C**) Representative H&E-stained liver sections from both groups show no histopathological abnormalities, suggesting no systemic toxicity due to ATG7i-SPNP treatment.

**Supplemental Figure 40**

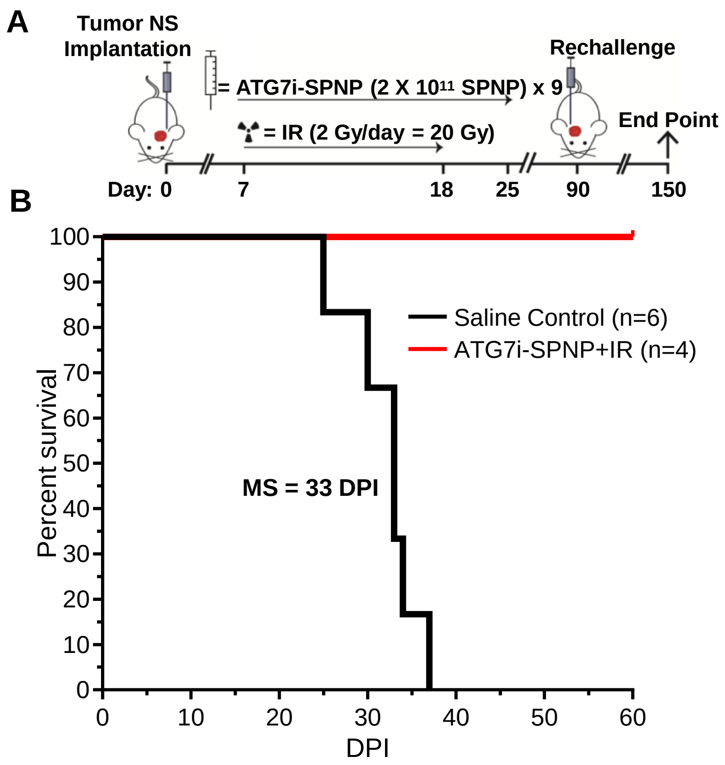


**Supplemental Figure 40:** Long-term survivors were rechallenged with genetically engineered mIDH1 mouse glioma NS. (**A**) Experimental design detailing when mice treated with ATG7i-SPNPs were considered long-term survivors, at which point they were rechallenged. (**B**) Kaplan-Meier curve showing the survival of long-term survivor mice that were rechallenged with genetically engineered mouse mIDH1 glioma NS.

**Supplemental Figure 41**

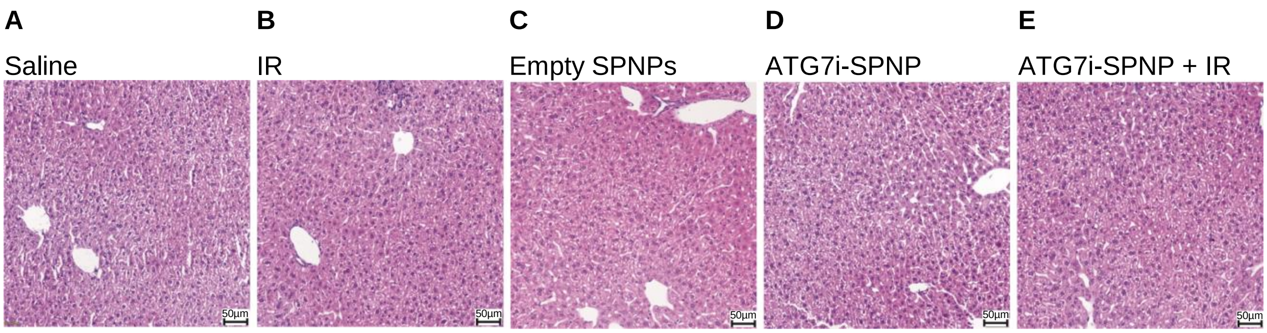


**Supplemental Figure 41:** Histopathological assessment of livers from tumor bearing mice treated with ATG7i-SPNPs + IR. H&E staining of 5µm paraffin embedded liver sections from (**A**) Saline (20X), (**B**) IR (20X), (**C**) empty SPNPs (20X), (**D**) ATG7i-SPNPs (20X) and (**E**) ATG7i-SPNP + IR (20X) treatment groups. Histology performed on resected livers following complete treatment of NPAI tumor bearing mice. Representative images from an experiment consisting of independent biological replicates are displayed. Black scale bars = 50µm.

**Supplemental Figure 42**

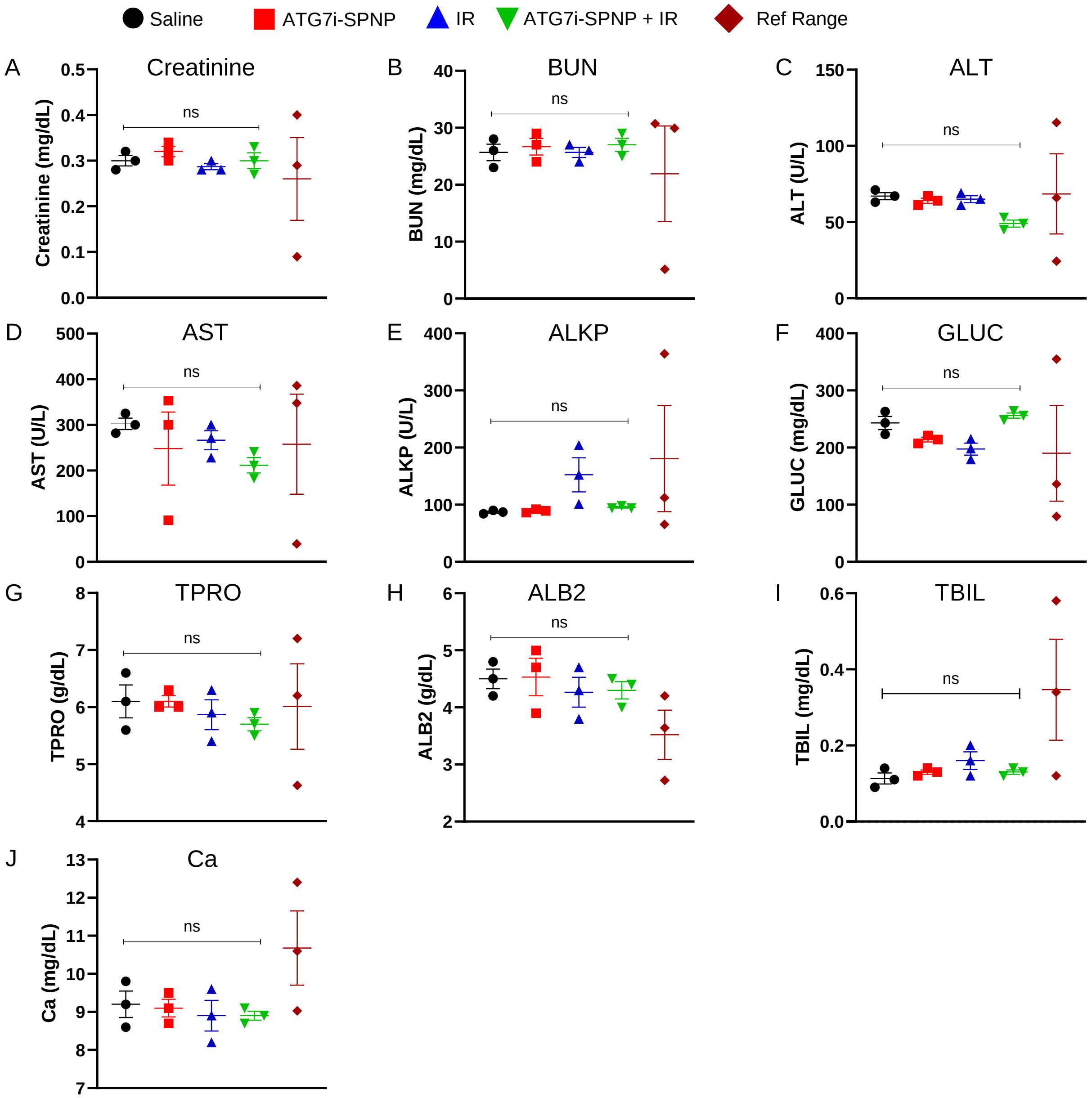


**Supplemental Figure 42:** Mouse serum biochemical analysis following intravenous ATG7i-SPNP in combination with IR treatment. NPAI OVA tumor bearing mice treated with ATG7 inhibitor ATG7i-SPNPs in combination with IR exhibit normal serum biochemical parameters compared with saline treated control. Serum was collected from tumor bearing mice treated with saline, ATG7i-SPNPs, IR, or ATG7i-SPNP + IR at 23 DPI. For each treatment group levels of (**A**) Creatinine, (**B**) BUN, (**C**) ALT, (**D**) AST, (**E**) ALKP, (**F**) GLUC, (**G**) TPRO, (**H**) ALB2, (**I**) TBIL, and (**J**) Ca were quantified. The levels of different serum biochemical parameters between the treatment groups were compared and were found non-significant (n = 3 biological replicates).

Supplemental Figure 43

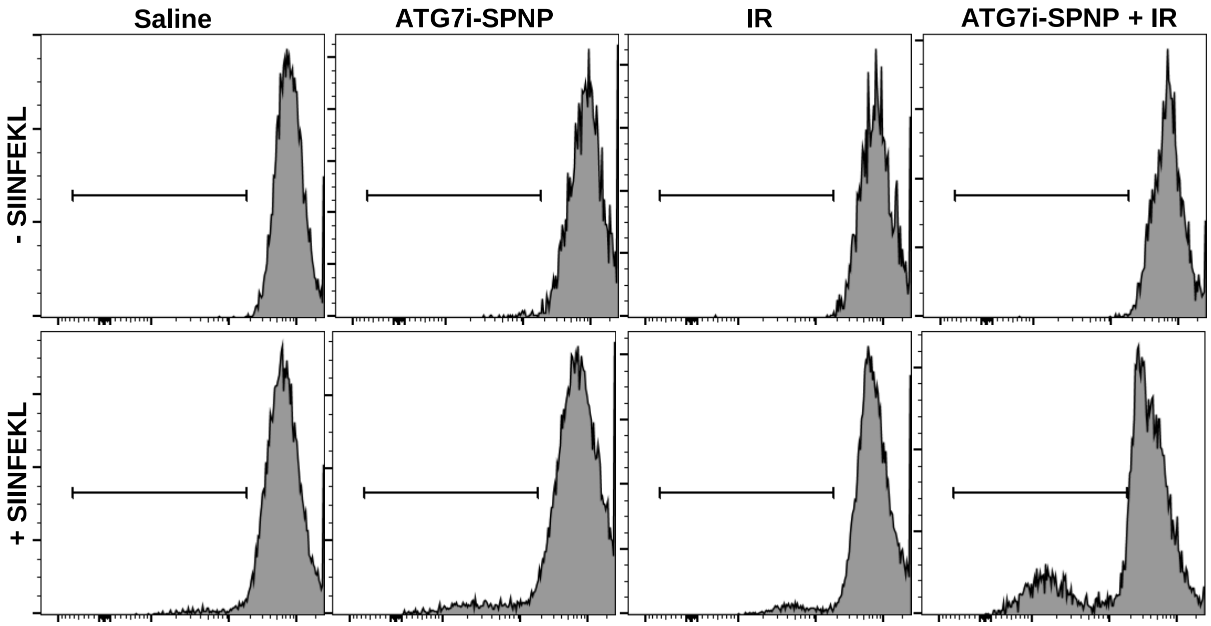


**Supplemental Figure 43:** ATG7i-SPNP treatment + IR leads to enhanced T cell proliferation and activation in response to SIINFEKL stimulation of splenocytes. T cell proliferation was measured as the reduction of CFSE staining in the CD45+/CD3+/CD8+ population. Upper panel: Representative histograms show CFSE stains of unstimulated splenocytes (inactivated T cells) and Lower panel: Representative histograms show 100nM SIINFEKL-induced T cell proliferation from Saline, ATG7i-SPNP, IR and ATG7i-SPNP + IR treated groups.

**Supplemental Figure 44**

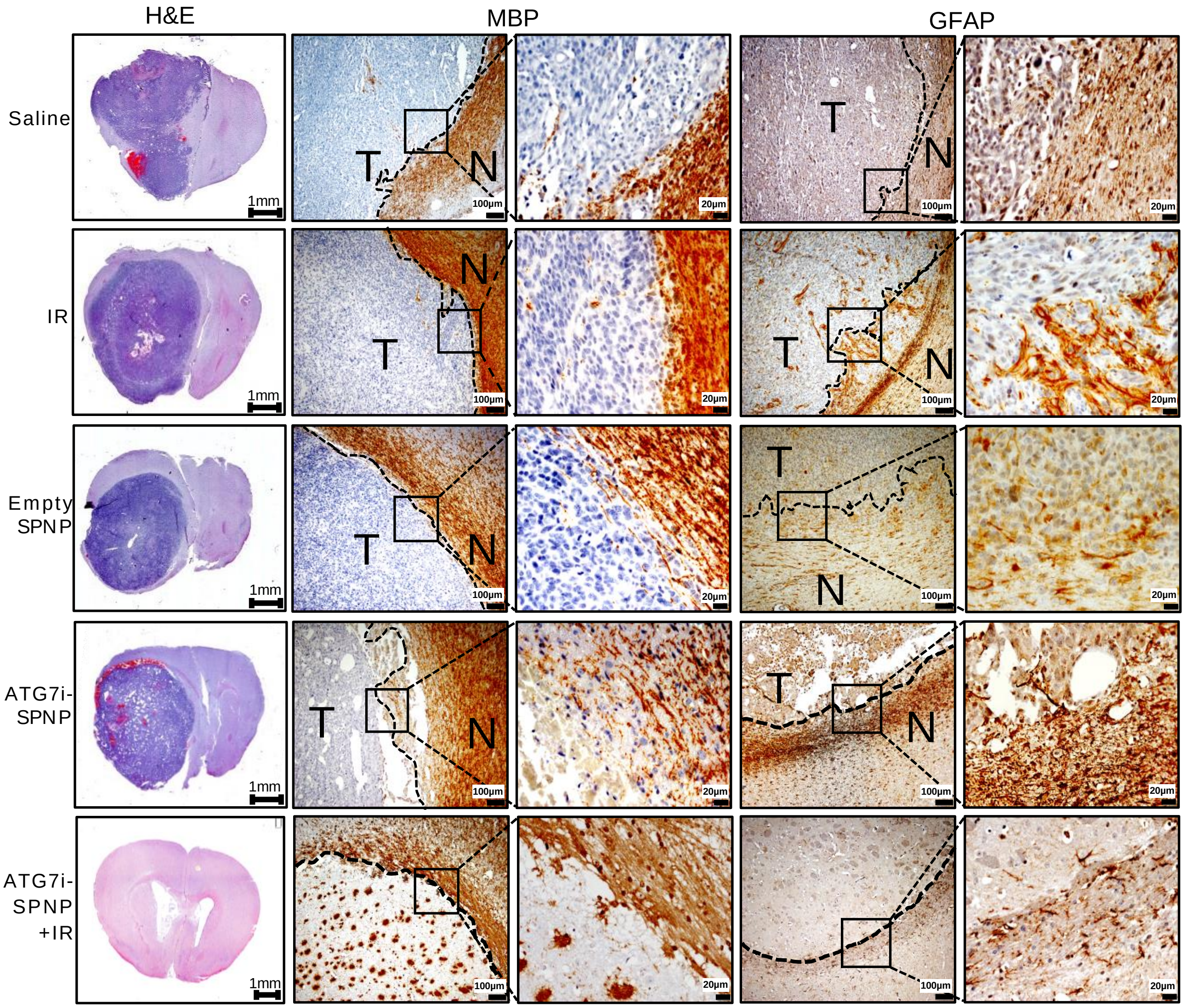


**Supplemental Figure 44:** Histopathological assessment of brains from tumor bearing mice treated with ATG7i-SPNP +IR. H&E staining of 5µm paraffin embedded brain sections from saline, IR, Empty SPNP, ATG7i-SPNP, and long-term survivors from ATG7i-SPNP+IR treatment groups (scale bar=1mm). Paraffin embedded 5µm brain sections for each treatment group were stained for myeline basic protein (MBP), and glial fibrillary acidic protein (GFAP). Low magnification (10X) panels show normal brain (N) and tumor (T) tissue (black scale bar = 100 µm). High magnification (40X) panels (black scale bar = 20 µm) indicate positive staining for areas delineated in low-magnification panels. Representative images from a single experiment consisting of independent biological replicates are displayed.

**Supplemental Figure 44**

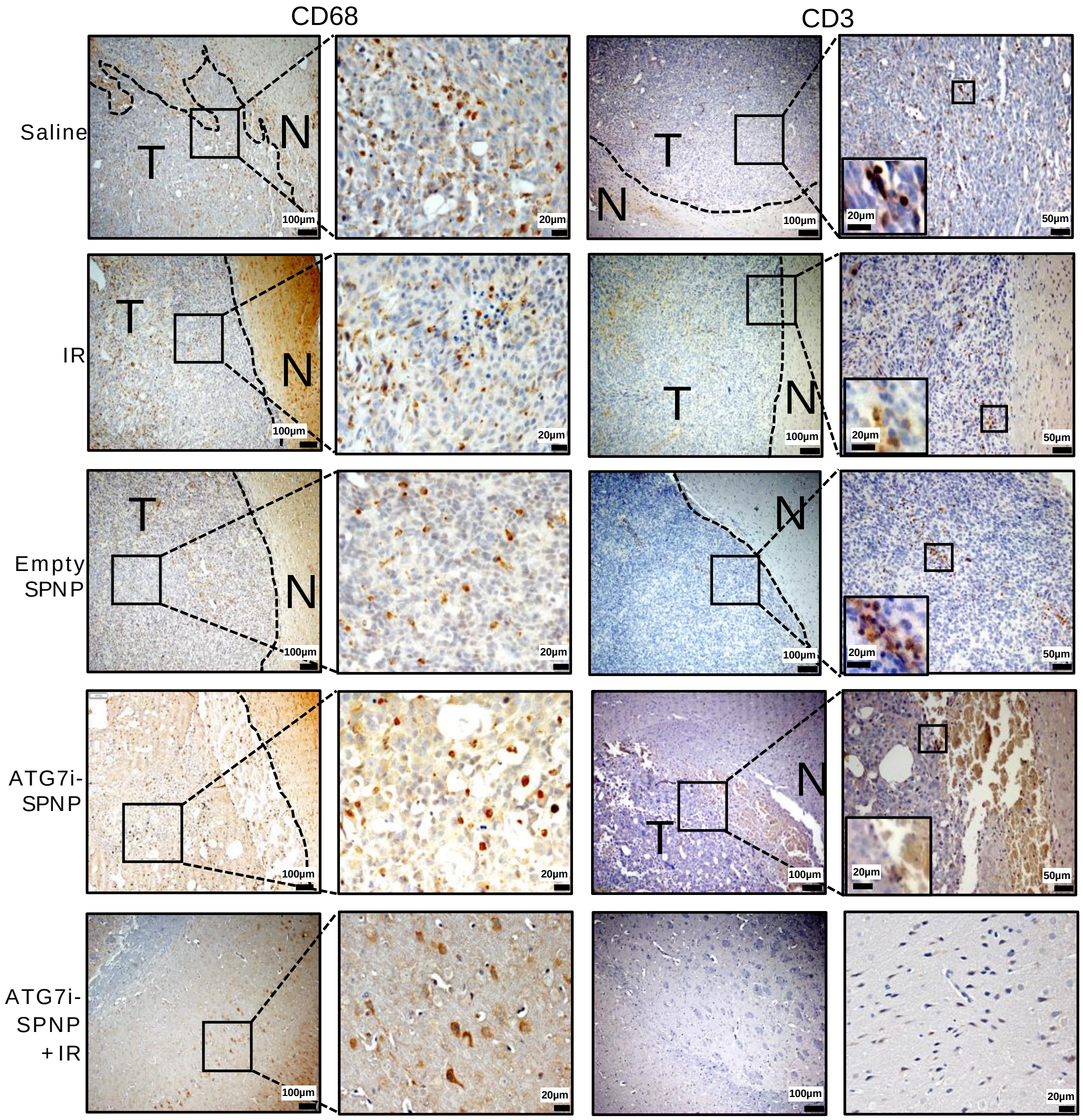


**Supplemental Figure 44:** Histopathological assessment of brains from tumor bearing mice treated with ATG7i-SPNP+IR. Paraffin embedded 5µm brain sections of the following treatment groups: saline, IR, Empty SPNP, ATG7i-SPNP, and long-term survivors from ATG7i-SPNP+IR were stained for CD68 and CD3. Low magnification (10X) panels show normal brain (N) and tumor (T) tissue (black scale bar = 100 µm). High magnification (40X) panels (black scale bar = 50 µm for saline, IR, Empty SPNP, and ATG7i-SPNP and scale bar = 20 µm for ATG7i-SPNP+IR long-term survivors) indicate positive staining for areas delineated in low-magnification panels. Representative images from a single experiment consisting of independent biological replicates are displayed.

**Supplemental Table 3**

| siRNA | Company | Catalog # | Sequence |
|---|---|---|---|
| ATG4b (Mouse) | Origene | SR413163 | rUrArGrCrCrArArUrArArArGrUrArGrUrGrGrGrCr ArCrUrGrUrUrGrUrUrGrUrUrUrUrCrUrArCrUrGr ArArArCrUrUrArUrUrUrArUrCrArGrArGrUrArUr CrArUrArUrGrUrCrArArArGrUrGrGrCrUrGrCrGrU |
| ATG7 (Mouse) | Origene | SR427399 | rCrUrUrGrArUrCrArGrUrArCrGrArGrCrGrArGrAr ArGrGATrCrArUrCrArArGrGrGrCrUrArUrUrArCr UrArCrArArUrGGTrGrCrUrCrUrGrArArCrUrCrAr ArUrArArUrArArCrCrUrUGG |
| ATG4b (Human) | Origene | SR323518 | rArGrArUrGrGrArCrGrCrArGrCrUrArCrUrCrUrGr ArCrCTArGrCrArUrUrGrArArGrArCrArUrArGrUr GrUrArUrUrUrCCTrGrUrCrCrArCrArUrUrGrCrAr ArUrGrGrArCrArArCrArCTG |
| ATG7 (Human) | Origene | SR323157 | rGrArGrUrCrArUrCrArGrUrGrGrArUrCrUrArArAr UrCrUCArGrCrCrGrUrGrGrArArUrUrGrArUrGrGr UrArUrCrUrGrUTTrGrUrGrArArUrArUrCrArArAr UrArCrCrArArUrCrUrUrAGA |
| ATG9b (Mouse) | Origene | SR421223 | rCrArArUrGrArUrUrGrArArArArUrGrArGrArCrUr UrGrUrUrCrArA |
| ATG9b (Human) | Origene | SR317285 | rCrCrUrGrGrArGrArUrUrArUrCrGrArCrUrUrUrUrU rUrCAT |

**Supplemental Table 3:** ATG4b, ATG7, and ATG9b mouse and human siRNA respective sequences, company origin, and catalog number.

**Supplemental Table 4**

| Group | MS Change | P value |
|---|---|---|
| Saline (MS: 33 DPI) vs. Empty-SPNP (MS: 35 DPI) | + 2d | P = 0.0042 |
| Saline (MS: 33 DPI) vs. IR (MS: 37 DPI) | + 4d | P = 0.0653 |
| Saline (MS: 33 DPI) vs. ATG7i-SPNP (MS: 26 DPI) | - 7d | P = 0.0242 |
| Saline (MS: 33 DPI) vs. ATG7i-SPNP+IR (MS: 60 DPI) | + 27d | P < 0.0001 |
| Empty-SPNP (MS: 35 DPI) vs. IR (MS: 37 DPI) | + 2d | P = 0.07025 |
| Empty-SPNP (MS: 35 DPI) vs. ATG7i-SPNP (MS: 26 DPI) | - 9d | P = 0.0003 |
| Empty-SPNP (MS: 35 DPI) vs. ATG7i-SPNP+IR (MS: 60 DPI) | + 25d | P = 0.0001 |
| IR (MS: 37 DPI) vs. ATG7i-SPNP (MS: 26 DPI) | - 11d | P = 0.0064 |
| IR (MS: 37 DPI) vs. ATG7i-SPNP+IR (MS: 60 DPI) | + 23d | P = 0.0003 |
| ATG7i-SPNP (MS: 26 DPI) vs. ATG7i-SPNP+IR (MS: 60) | + 34d | P < 0.0001 |

**Supplemental Table 4:** Log-rank (Mantel-Cox) Kaplan Meier detailed survival analysis of animals implanted with genetically engineered mIDH1 glioma mouse cells treated with ATG7i-SPNP in combination with IR.
